# A multi-tissue epigenomic atlas links the sheep non-coding genome to domestication and complex traits

**DOI:** 10.64898/2026.08.22.746371

**Authors:** Zhu Meng, Shiwen Zhang, Ziyang Zhuang, Jiayi He, Fei Liu, Lingyun Wu, Xiangrong Sun, Jingsheng Lu, Xinyi Li, Hao Yang, Mian Gong, Xiaolong Du, Qiao Mei, Hao Li, Guoqing Zhang, Xiaoyun He, Jianning He, Xiaofei Guo, Sijia Nie, Xiaoxu Zhang, Mingzhu Shan, Jianqi Yang, Xinyue Li, Huaiqiang Yang, Yangshuo Hu, Yuchen He, Shuo Tian, Xianfeng Wu, Xudong Wu, Kai Zhan, Mingshan Wang, Xuemei Lu, Huaijun Zhou, Guiping Zhao, Pengju Zhao, Yinghui Ling, Chunhuan Ren, Youbing Yang, Xiaosheng Zhang, Yu Jiang, Weiwei Wu, Ruidong Xiang, Richard P. M. A. Crooijmans, Ole Madsen, David E. MacHugh, Kevin G. Daly, Emily L. Clark, Lingzhao Fang, Zijun Zhang, Mingxing Chu, Dailu Guan, Zhangyuan Pan

## Abstract

Non-coding regulatory variation drives complex traits, domestication, and evolutionary adaptation, yet the sheep genome lacks high-resolution functional annotation. Here we present SheepEpimap, a multi-tissue regulatory atlas harmonizing 516 CUT&Tag histone modifications, ATAC-seq, and RNA-seq datasets across 43 adult tissues in sheep. We annotated 2.93 million *cis*-regulatory elements, yielding 557,441 enhancer–gene pairs and 145,407 variants with allele-specific effects. By training a sequence-to-function deep-learning model, we decoded the base-pair syntax of chromatin accessibility, annotated transcription factor motif instances genome-wide, and constructed 12,210 tissue-specific gene regulatory networks (GRNs). Integrating this resource with multi-tissue expression quantitative trait loci, selection sweeps, and genome-wide association studies prioritized non-coding variants driving domestication and complex traits. Finally, cross-species analysis revealed that sequence-conserved, tissue-matched enhancers were significantly enriched in heritability of complex traits and diseases in humans. In summary, SheepEpimap (https://genome.ucsc.edu/s/mengzhu/SheepEpimap) provides an open-access foundational ecosystem for sheep functional genomics, precision breeding, and comparative biology.

## Introduction

Sheep (*Ovis aries*) were among the earliest domesticated livestock species, serving as a cornerstone of human societies since the Neolithic transition by offering meat, milk, wool, and hides^1,2^. Through global dispersal, extensive artificial selection, and diverse environmental pressures, sheep have evolved specific adaptations in metabolic specialization, reproductive biology, and extreme climate resilience^3,4^. Globally vital sheep farming boasted 1.363 billion heads and $73.95 billion farm output in 2024, with 2,628 sheep breed records across 174 countries^5,6^. Improving complex traits such as growth efficiency, fertility, wool quality, and disease resistance in sheep is thus central to sustainable global production and environment. However, like most complex traits in mammals, these phenotypes are predominantly shaped not by protein-coding alterations, but by non-coding regulatory variants that modulate gene expression in highly tissue-specific and spatiotemporal contexts^7–10^. Beyond agriculture, the physiological compatibility, comparable organ size, and developmental tempo of sheep have established them as potential large-animal models for human biomedical research, particularly in cardiovascular, musculoskeletal, and neurodegenerative disorders^11–13^. Consequently, a high-resolution, tissue-resolved annotation of the sheep regulatory genome is crucial to decrypting the mechanisms underlying domestication, complex traits, and translational biomedicine.

Deciphering the functional syntax of the non-coding genome remains a fundamental challenge in modern biology. In human genomics, the field has transitioned from descriptive consortia catalogs (e.g., ENCODE and NIH Roadmap Epigenomics) to advanced predictive frameworks like AlphaGenome and ChromBPNet, which model how DNA sequence variation can influence molecular phenotypes^14–16^. Livestock functional genomics has recently begun to adopt these integrative paradigms through the international Functional Annotation of Animal Genomes (FAANG) and Farm Animal Genotype-Tissue Expression (FarmGTEx) projects^17,18^. For instance, pig and chicken FAANG resources have progressed beyond simple peak catalogs by defining multi-tissue *cis-*regulatory elements and showing that trait- and domestication-associated variants are enriched in active regulatory elements^19,20^. Complementing these epigenomic maps, FarmGTEx-scale analyses in pig^21^, cattle^22^ and chicken^23^ have charted multi-tissue regulatory-variant landscapes and linked them to agronomic phenotypes. A recent bovine atlas further integrated multi-tissue regulatory maps with sequence-to-function models for variant prioritization and GWAS fine-mapping^24^. Despite these milestones, progress in small ruminant genomics has lagged. Although previous epigenomic studies in sheep established an important foundation by reporting tissue-specific *cis*-regulatory elements (CREs) and baseline chromatin marks, these resources remain constrained by either narrow tissue representation, a limited repertoire of epigenetic assays, or the absence of gene regulatory network inference^25,26^. More recently, our SheepGTEx project made a major leap forward by detecting hundreds of thousands of genetic variants associated with a diverse range of molecular phenotypes (molQTL)^27^. However, the precise mechanistic impact of these non-coding variants on local enhancer activity, promoter usage, and allele-specific regulatory imbalances has not been systematically dissected. No single unified sheep resource has yet integrated high-density tissue coverage, multi-state chromatin modeling, physical enhancer-to-gene topologies, sequence-driven deep learning network inference, and multi-layer allele-specific imbalances. Such a holistic framework is urgently required to mechanistically interpret molQTL, resolve complex trait GWAS loci, and decode evolutionary selection sweeps in small ruminants.

To address these limitations, we present SheepEpimap, a multi-tissue atlas of the sheep regulatory genome by generating and integrating 516 CUT&Tag, ATAC-seq and RNA-seq datasets across 43 adult tissues in sheep (**Fig. 1a,b**). SheepEpimap substantially expands both the number of datasets and tissue coverage relative to previous sheep regulatory resources^25,26^. We use an 11-state chromatin model to annotate CREs and combine motif analysis with deep learning sequence models to characterize regulatory sequence grammar and infer candidate tissue-associated gene regulatory networks. We map allele-specific variation across chromatin accessibility, histone modifications and transcription to examine relationships between sequence variation and chromatin or gene-expression imbalance (**Fig. 1c**). We integrate SheepEpimap with SheepGTEx, GWAS results for 34 complex traits and selection signatures to prioritize candidate effector genes and putative non-coding variants related to growth, reproduction, wool traits, immunity and domestication. Cross-species projection assesses the contribution of sequence-conserved, tissue-matched enhancers to human complex-trait and disease heritability (**Fig. 1d**). The open-access SheepEpimap supports mammalian regulatory genomics, complex-trait genetics, selective breeding, comparative biology and translational biomedicine.

**Fig. 1.**
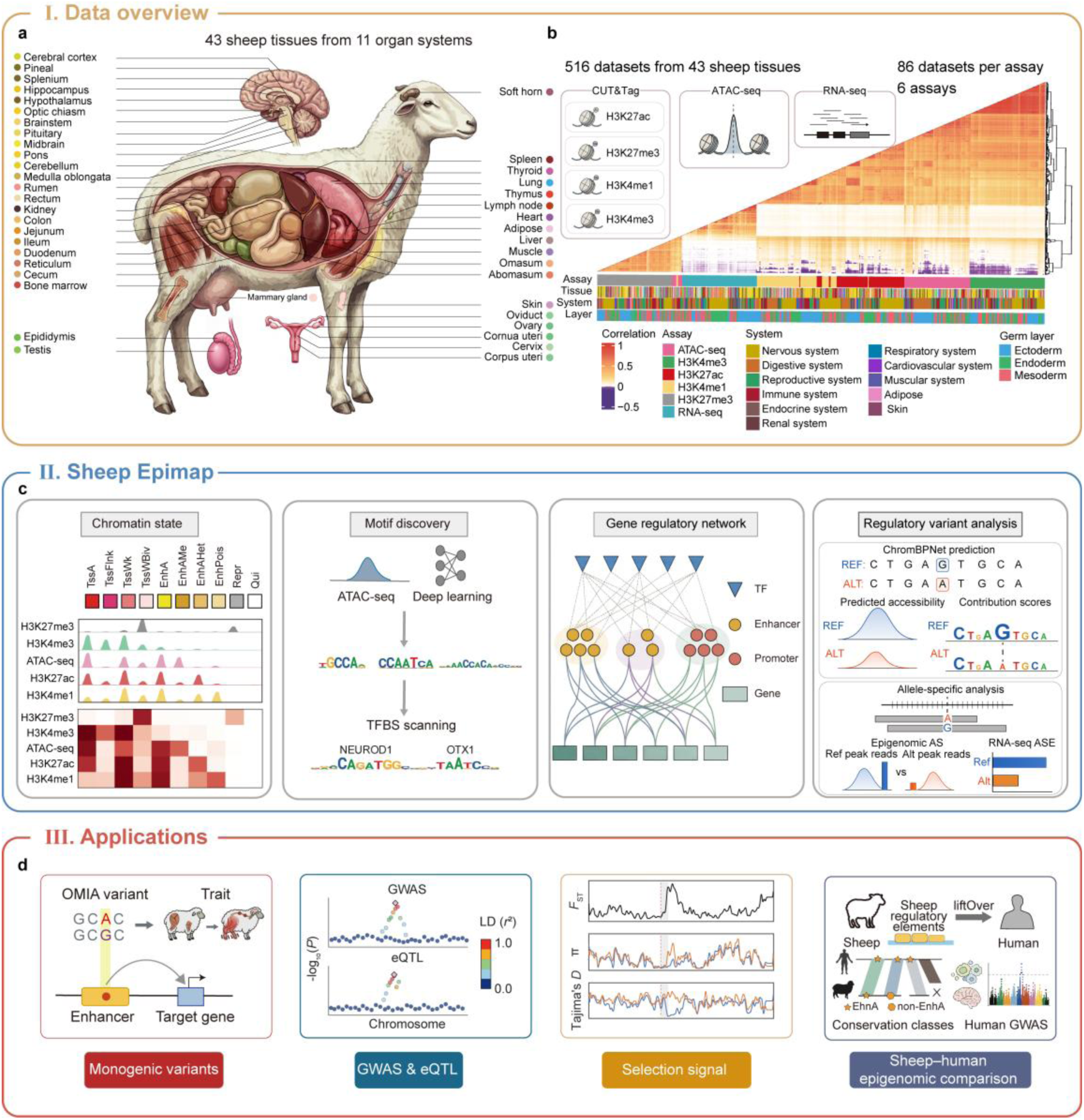
Sheep multi-omics regulatory atlas. a, Overview of 43 sampled adult sheep tissues from 11 organ systems. Tissue identities and colour coding are shown below the illustration. b, Summary of 516 matched multi-omics datasets generated across 43 tissues and six assays: RNA-seq, ATAC-seq and CUT&Tag profiling of H3K27ac, H3K27me3, H3K4me1 and H3K4me3. The heatmap shows pairwise Pearson correlations among datasets; annotation bars indicate assay, tissue group and germ-layer origin. c, Overview of SheepEpimap construction, including chromatin-state annotation, deep-learning-based motif discovery, transcription factor-binding-site scanning, gene regulatory network construction and regulatory variant analysis using ChromBPNet prediction and allele-specific analysis d, Representative applications of the atlas for interpreting monogenic variants, GWAS and eQTL signals, population selection signatures and sheep–human regulatory conservation. ATAC-seq, assay for transposase-accessible chromatin using sequencing; CUT&Tag, Cleavage Under Targets and Tagmentation; RNA-seq, RNA sequencing; TF, transcription factor; TFBS, transcription factor-binding site; SNP, single-nucleotide polymorphism; OMIA, Online Mendelian Inheritance in Animals; GWAS, genome-wide association study; eQTL, expression quantitative trait locus; LD, linkage disequilibrium; *F*_ST_, fixation index; π, nucleotide diversity.

## Results

### Overview of functional genomic data in the SheepEpimap project

To establish a comprehensive, high-resolution map of the sheep regulatory genome, we systematically generated 516 multi-omic datasets spanning 43 adult tissues that represent 11 distinct physiological systems. This cohort comprises 344 CUT&Tag profiles mapping four histone modifications (*i.e.* H3K4me1, H3K4me3, H3K27ac, and H3K27me3), together with 86 ATAC-seq profiles and 86 matching RNA-seq datasets (**Fig. 1a-b**). The sequencing experiment yielded 14.88 billion raw reads, of which 13.99 billion (94.02%) were uniquely mapped and retained after alignment and filtering (**Supplementary Table 1**). Following quality control, experimental tracks with a FRiP score below 5% or fewer than 1,000 peaks were excluded. Missing or low-quality tracks were subsequently imputed using ChromImpute^28^, yielding 73 imputed data tracks and substantially improving tissue and assay completeness across the atlas (**Supplementary Fig. 1 and 2**).

Following a stringent peak-calling pipeline (**Methods**), we identified 1,188,652 H3K27ac, 1,921,007 H3K27me3, 3,383,867 H3K4me1, 967,269 H3K4me3 and 1,704,580 ATAC-seq non-redundant peaks, covering 8.91%, 18.72%, 28.18%, 4.73%, and 9.47% of the sheep genome, respectively (**Extended Data Fig. 1a-d; Supplementary Table 1**). Cross-assay correlation analyses revealed that active chromatin features (including H3K4me3, H3K27ac, H3K4me1, and ATAC-seq signals) were positively correlated with each other and with gene expression. In contrast, these active marks were negatively correlated with H3K27me3-marked regions, mirroring the canonical regulatory coordination between active *cis*-elements and Polycomb-associated repressive chromatin states^7^ (**Fig. 1b; Supplementary Fig. 3-4**). As an example, *RXFP2* showed highly tissue-specific expression in soft-horn tissue together with concordant H3K27ac enrichment and chromatin accessibility. Previous genetic and transcriptomic studies have linked this locus to horn development and morphology in sheep^29–31^ (**Extended Data Fig. 1e; Supplementary Fig. 5**). These multilayer signals illustrate the ability of SheepEpimap to resolve tissue-specific regulatory activity across the sheep genome. **Construction of a sheep regulatory element atlas**Leveraging the multi-omic datasets described above characterize, we sought to systematically partition the sheep non-coding genome into discrete functional domains. By integrating our coordinated histone modification and chromatin accessibility profiles via ChromHMM^32^, we defined an 11-state core chromatin model to annotate the sheep regulatory genome^33^ (**Supplementary Fig. 6a**). These 11 chromatin states were consolidated into four primary functional classes: promoter states (termed TssA, TssFlnk, TssWk, and TssWBiv), which collectively spanned 3.12% of the genome; enhancer states (EnhA, EnhAMe, EnhAHet and EnhPois), covering 4.97%; a repressed state (Repr), occupying 4.20%; and quiescent states (QuiW and Qui), which accounted for the remaining 87.72% of the genome (**Fig. 2a; Supplementary Table 3**). Across all 43 tissues, we identified 2,931,347 non-redundant regulatory elements after excluding QuiW and Qui, including 675,096 promoters, 2,004,542 enhancers and 251,709 repressed regions (**Supplementary Fig. 6b-h; Supplementary Table 3**). The annotated chromatin states showed the expected enrichment profiles for DNA methylation, open chromatin accessibility, evolutionary sequence conservation, and significant overlapping cross-validation with experimentally verified mammalian VISTA enhancer sequences^19,20,24,34^ (**Fig. 2b,c; Supplementary Figs. 7** and 8).

**Fig. 2.**
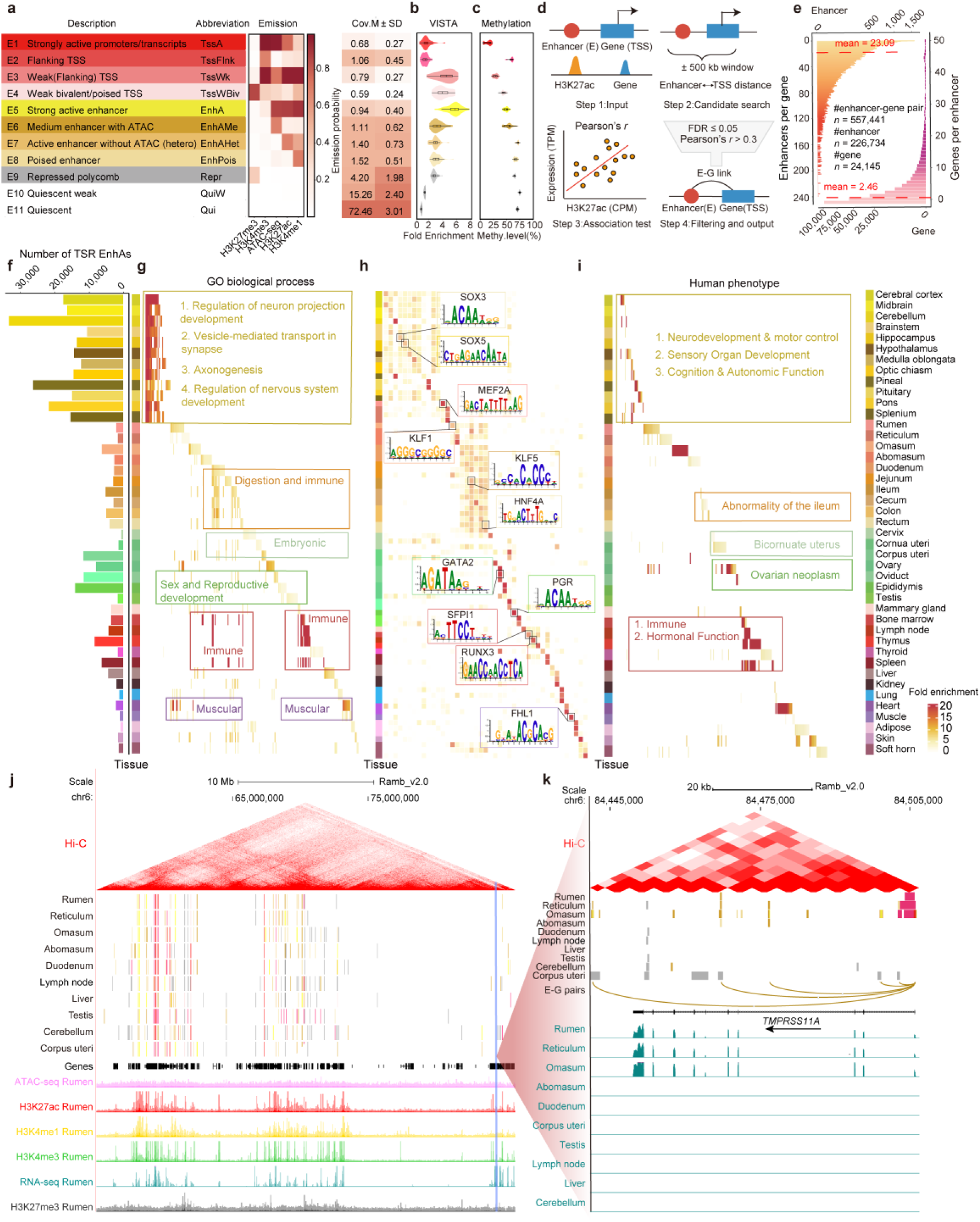
The regulatory element atlas across 43 tissues in sheep. a, Eleven chromatin states inferred from H3K27ac, H3K4me3, ATAC-seq, H3K4me1 and H3K27me3 signals across sheep tissues, showing state annotations, abbreviations, emission probabilities and genomic coverage. The heatmap shows emission probabilities for each epigenomic feature. b, Fold enrichment of chromatin states in VISTA enhancer annotations. c, DNA methylation levels across chromatin states (Methods). d, Schematic workflow for enhancer–gene association analysis based on enhancer H3K27ac signals, gene expression and genomic distance to transcription start sites. e, Distribution of enhancer–gene associations, showing the number of enhancers linked to each gene and the number of genes linked to each enhancer. f, Distribution of tissue-specific active enhancers across 43 sheep tissues. Colors denote tissue categories. g, Gene Ontology biological process enrichment of genes associated with tissue-specific active enhancers. h, Transcription factor motif enrichment of tissue-specific active enhancers, with representative sequence logos. i, Human phenotype enrichment based on human orthologs of genes associated with tissue-specific active enhancers. j, Genome-browser view integrating Hi-C interactions, chromatin states, gene annotations and multi-omics signals (Methods). k, Representative *TMPRSS11A* locus showing Hi-C interactions, chromatin states, predicted enhancer–gene associations and RNA-seq signals across tissues.

Furthermore, we used the ROSE algorithm to identify super-enhancers^35^, resulting in 141,809 super-enhancers across 43 tissues, ranging from 2,158 in the pineal gland to 4,727 in the uterine horns (**Extended Data Fig. 2; Supplementary Table 5**). To link enhancers to putative target genes, we correlated H3K27ac signal intensity with gene expression (**see Methods,** **Fig. 2d**). This analysis linked 226,734 unique enhancers to 24,145 genes, yielding 557,441 enhancer–gene pairs (**Fig. 2e**). Compared with EnhA elements, super-enhancers showed a significantly higher number of putative target genes per element and a significantly lower number of elements per target gene (**Extended Data Fig. 2g; Extended Data Fig. 3; Supplementary Fig. 9**). Collectively, these results nominate candidate connections between CREs and target loci, providing a tissue-resolved regulatory map of the sheep genome.

## Functional characteristics of tissue-specific chromatin states

Tissue-specific CREs are key components of regulatory programs that establish cellular identity and support tissue-specific functions^7,36,37^. Across the SheepEpimap atlas, we identified 1,917,868 tissue-specific regulatory elements. This includes 338,228 tissue-specific EnhA elements, ranging from 564 in cervix tissue to 33,601 in the cerebellum (**Fig. 2f; Supplementary Fig. 10; Supplementary Table 7**). We next investigated their functional relevance through Gene Ontology (GO), transcription factor (TF) motif and phenotype enrichment analyses using their putative target genes (see **Methods**). GO enrichment patterns were broadly concordant with the expected physiological functions of the corresponding tissues. For example, enhancers specific to nervous-system tissues were linked to genes enriched for neurite-development processes, while muscle-specific enhancers were linked to genes involved in muscle-organ and muscle-cell development (**Fig. 2g; Supplementary Table 8**). Consistent with these observations, motif analyses highlighted lineage-associated TF signatures within tissue-specific EnhA elements across organ systems. For example, nervous-system-specific EnhA elements were enriched for SOX family members, involved in neural lineage specification and neuronal differentiation^38^, while heart-specific EnhA elements were enriched for MEF2A, a conserved DNA-binding transcription factor involving in muscle development and differentiation^39^ (**Fig. 2h; Supplementary Table 9**). Phenotype enrichment based on human and mouse orthologues of enhancer-linked genes also identified terms related to the corresponding tissues. Nervous-system- and ovary-specific elements were associated with neurodevelopment and ovarian neoplasm, respectively, whereas intestine- and uterus-specific elements were associated with abnormal intestine morphology and abnormal uterus morphology, respectively (**Fig. 2i; Supplementary Fig. 11a; Supplementary Tables 10, 11**). As a representative example, interactive visualization of the *TMPRSS11A* locus highlighted concordant patterns of chromatin state, enhancer-gene associations, gene expression activity (**Fig. 2j,k**). *TMPRSS11A* encodes a type II transmembrane serine protease of the HAT/DESC cluster, a group of epithelial proteases implicated in epithelial proteolytic regulation in a tissue-specific manner^40,41^. Together, these analyses suggest that tissue-specific EnhA elements are associated with tissue-associated TF activity and functionally coherent gene-expression programs.

## ChromBPNet reveals sequence syntax underlying chromatin accessibility

To transition from a descriptive catalog of chromatin features to a predictive understanding of the ovine non-coding genome, we sought to extract the primary sequence grammar governing tissue-specific nuclear architecture. We trained tissue-specific ChromBPNet deep-learning models directly on our multi-tissue ATAC-seq profiles across all 43 tissues^42,43^ (**Fig. 3a**). Models with Pearson’s *r* > 0.5 between predicted and observed accessibility in held-out test regions were retained (**Fig. 3b; Extended Data Fig. 4a; Supplementary Table 12**). The retained models showed concordance with independent datasets and observed ATAC-seq signals, yielding high-quality models for 38 tissues (**Fig. 3b; Extended Data Fig. 4b–d; Supplementary Table 13**).

**Fig. 3.**
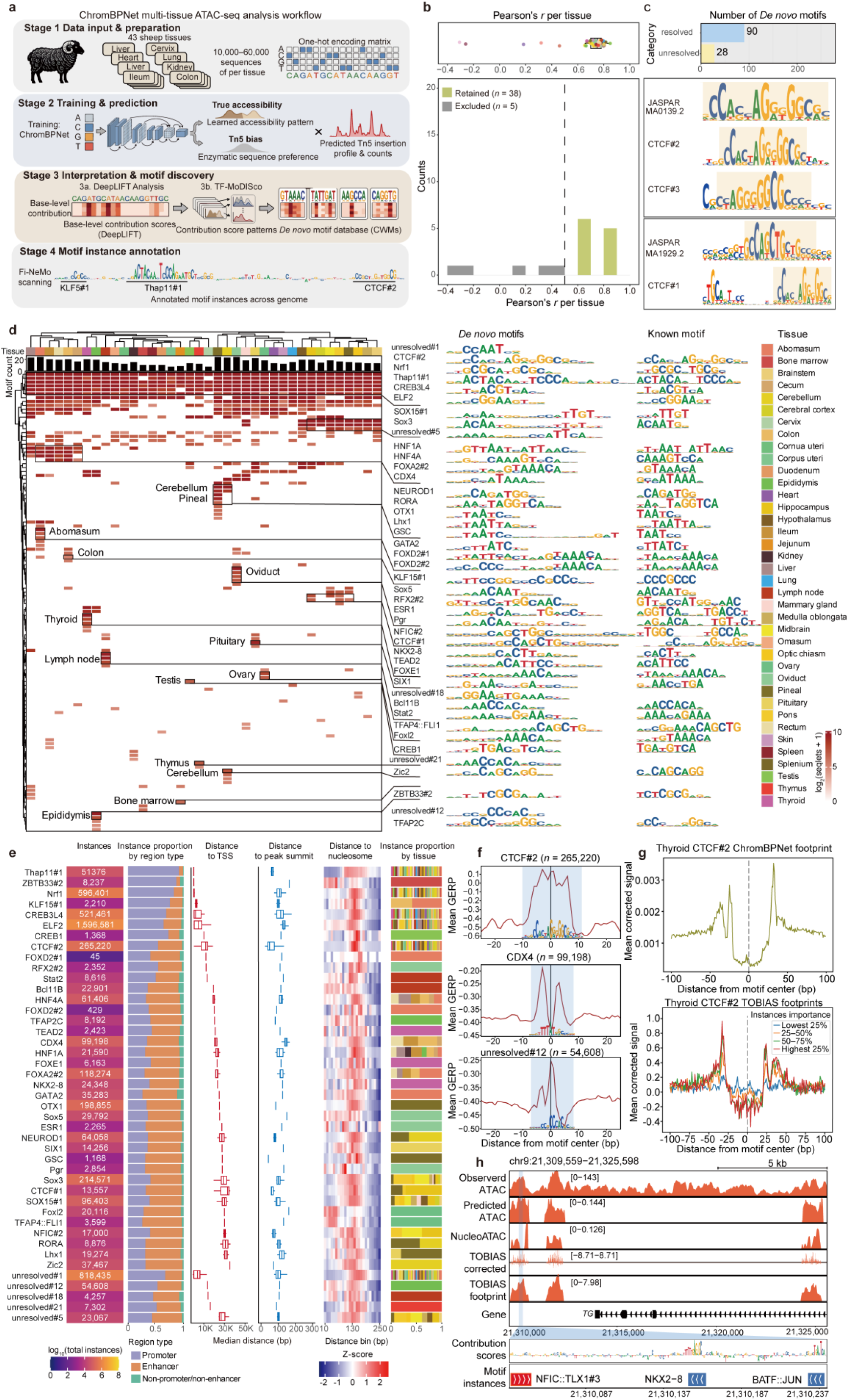
ChromBPNet analysis of sequence features underlying chromatin accessibility across sheep tissues. a, Workflow for training and interpreting tissue-specific ChromBPNet models using multi-tissue sheep ATAC-seq data. b, ChromBPNet model performance across tissues, measured by Pearson’s *r* between predicted and observed log-transformed counts on held-out data. c, Summary of *de novo* motifs identified from ChromBPNet contribution scores, including resolved and unresolved motifs and representative CTCF motif comparisons. d, Tissue-level atlas of *de novo* motifs identified by TF-MoDISco, shown as log_2_(seqlets + 1), with corresponding motif logos and closest known motif matches. e, Genome-wide properties of predicted motif instances, including instance abundance, genomic annotation, distances to TSSs, ATAC-seq peak summits and nucleosome dyads, and tissue distribution. f, Centered mean GERP conservation profiles for selected motifs. g, Footprint profiles for thyroid CTCF#2 motif instances derived from ChromBPNet predictions and TOBIAS-corrected ATAC-seq signals. Motif instances are stratified by model-derived importance score. h, Representative genome-browser view showing observed and predicted (ChromBPNet) ATAC-seq signals, nucleosome occupancy, TOBIAS-corrected and footprint signals, gene annotations, base-level contribution scores and predicted motif instances.

To extract mechanistic insights from these deep-learning structures, we leveraged backpropagation contribution scores to chart a comprehensive *de novo* motif atlas of sheep chromatin accessibility with TF-MoDISco^44^. This analysis identified 118 *de novo* motifs, of which 90 matched known motifs, and 28 remained unannotated. A representative CTCF-associated contribution weight matrix (CWM) closely matched the corresponding JASPAR^45^ CTCF motif, indicating that ChromBPNet recovered canonical TF binding motifs from primary DNA sequences (**Fig. 3c; Supplementary Table 14**).

Crucially, rather than identifying redundant household sequence patterns, our deep-learning approach identified motif configurations associated with tissue-specific functions. The models differentiated broadly shared structural landmarks from highly specialized, tissue-restricted motifs, such as liver-enriched HNF4A, intestinal-enriched CDX4, and an epididymis-restricted, unannotated unresolved#12 motif (**Fig. 3d; Supplementary Fig. 12a; Supplementary Table 14).** To localize these motifs within open chromatin regions, we then used Fi-NeMo^15^ to map motif instances of all 118 *de novo* motifs within open chromatin regions, identifying an average of 91,516 instances per motif. These instances were broadly distributed across ATAC peaks and showed motif-specific preferences for promoters or enhancers, as well as distinct distances to transcription start sites (TSSs), peak summits and nucleosome dyads (**Fig. 3e; Extended Data Fig. 4e–h**). For instance, evaluating the abomasum-active GATA2 motif demonstrated that localized nucleotide contribution scores were heavily concentrated at the exact core consensus positions required for spatial transcription factor engagement (**Extended Data Fig. 4i**). To examine evolutionary constraint at these model-derived motifs, we retrieved precomputed GERP scores^46^ for genomic positions flanking motif centers. Genomic positions flanking the core centers of the ubiquitous CTCF#2, CDX4 and the tissue-restricted unresolved#12 motif showed higher GERP scores, consistent with stronger evolutionary constraint (**Fig. 3f**). Moreover, we cross-examined our deep-learning predictions by ChromBPNet with physical *in vivo* molecular footprints with TOBIAS-based footprinting^47^. Both orthogonal approaches recovered similar footprint patterns for CTCF#2, and motif instances with higher model importance showed stronger footprint signals (**Fig. 3g; Supplementary Fig. 13a–c; Supplementary Tables 16, 17).**

Locus-specific inspection showed that ChromBPNet-predicted accessibility, sequence contribution scores and motif instances jointly highlighted tissue-specific regulatory regions, although their local maxima did not always coincide; predicted accessibility profiles were broadly consistent with the observed tissue-specific ATAC-seq tracks (**Fig. 3h; Supplementary Fig. 12b; Extended Data Fig. 4j**). For example, a highly specialized open chromatin region flanking the thyroglobulin (*TG*) gene in thyroid tissue contained dense clusters of high-contribution thyroid-specific motifs, including *NKX2-8*, a key homeobox master regulator critical for thyroid organogenesis and metabolic specialization^48^ (**Fig. 3h**). These analyses were used to generate a genome-wide annotation of TF motif instances, which is available through the SheepEpimap browser (https://genome.ucsc.edu/s/mengzhu/SheepEpimap).

## Construction of a sheep gene regulatory network

We constructed multi-tissue gene regulatory networks (GRN) by integrating enhancers, promoters, TF sequence motifs by ChromBPNet, and target genes to dissect gene regulation in a systems genetic manner (**Fig. 4a**). Across tissues, we reconstructed 12,210 GRNs, with an average of 407 GRNs per tissue. Collectively, these GRNs involved 75 TFs, 1,946 enhancers, 619 promoters and 560 genes (**Fig. 4b-d; Supplementary Tables 18–20**). Network clustering revealed that CREs and target genes were non-randomly organized into distinct tissue-associated modules rather than being uniformly distributed across the global network (**Fig. 4c**). Notably, tissues from the nervous system, including the cerebellum, cerebral cortex, midbrain, hypothalamus, brainstem, and optic chiasm, formed prominent regulatory modules. Representative subnetworks further demonstrated the utility of this framework for identifying tissue-specific regulatory modules (**Fig. 4e**).

**Fig. 4.**
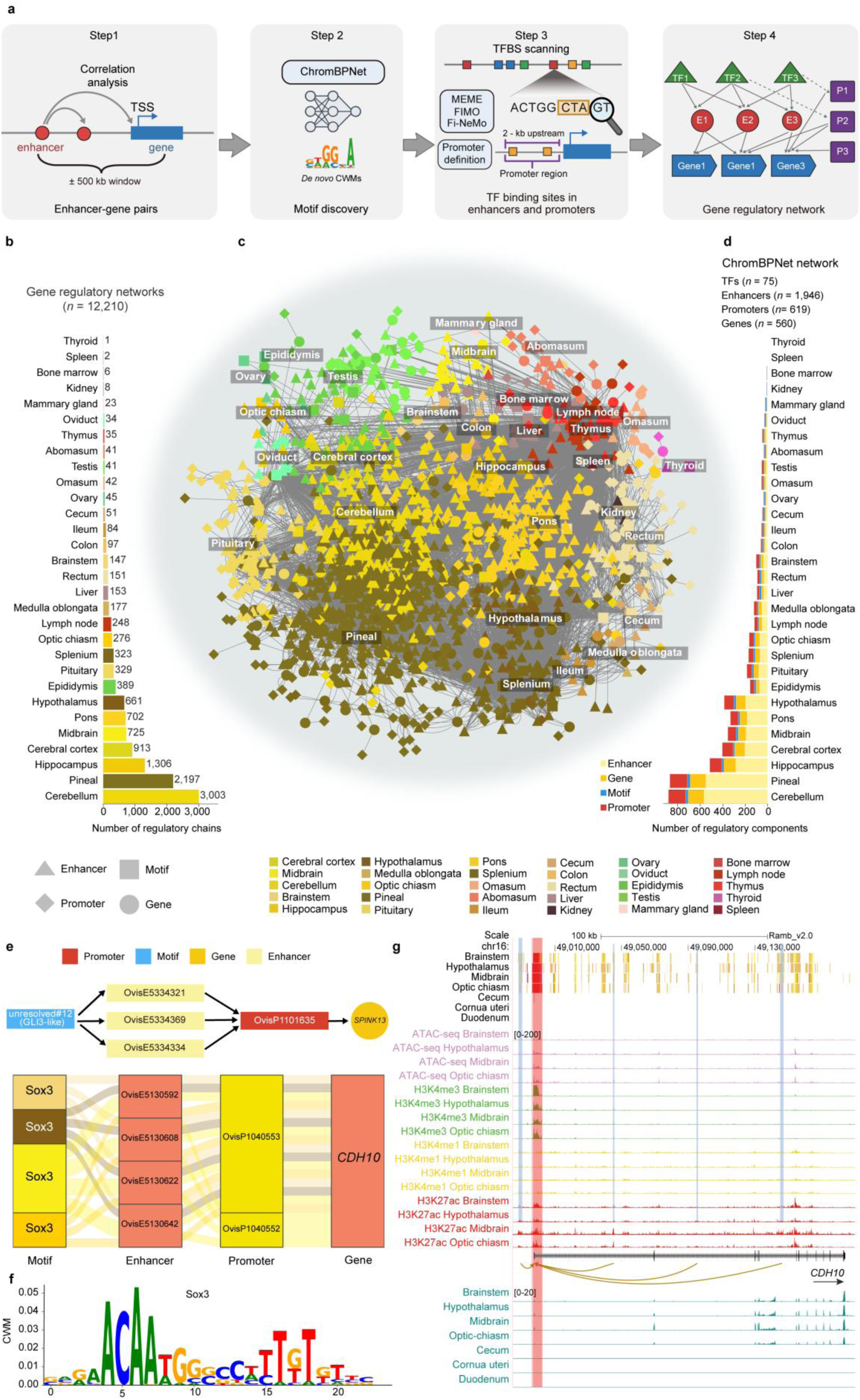
ChromBPNet-derived gene regulatory networks across sheep tissues. a, Schematic overview of network construction, including enhancer–gene pairing, *de novo* contribution weight matrix discovery, transcription factor-binding-site scanning in enhancers and promoters, and network assembly. b, Number of gene regulatory networks identified in each tissue. c, Global ChromBPNet-derived gene regulatory network arranged by tissue module. Node colors indicate tissue modules, and node shapes denote enhancers, motifs, promoters and genes, as indicated in the legends. d, Summary of the overall network composition and the distribution of node classes across tissue modules. e, Representative subnetworks involving unresolved#12, a motif with similarity to GLI3 (GLI3-like) and *SPINK13* (top), and Sox3 motifs and *CDH10* (bottom). Enhancer and promoter identifiers are shown within the corresponding nodes. f, Sequence logo of the Sox3 contribution weight matrix. g, Genome-browser view of the *CDH10* locus on chromosome 16, showing chromatin states across selected tissues, brain-tissue ATAC-seq, H3K4me3, H3K4me1 and H3K27ac profiles, enhancer–gene pairs, gene annotation and RNA-seq profiles across selected tissues. CWM, contribution weight matrix; TFBS, transcription factor-binding site.

Within the brain-associated module, the Sox3 motif was connected to multiple candidate enhancers, including OvisE5130592, OvisE5130608, OvisE5130622, and OvisE5130642, which were further linked to *CDH10*-associated promoters (OvisP1040553 and OvisP1040552). Given the established roles of Sox3 in neuronal development, and the *CDH10* in synapse formation and axon growth, this Sox3–enhancer/promoter–*CDH10* module may play a specific role in the neuronal system (**Fig. 4e-g**)^49–51^.

In an epididymis-associated module, an unresolved#12 motif, which showed a similarity to a GLI3 motif (*q*-value = 0.079), was connected to multiple candidate enhancers (OvisE5334321, OvisE5334369, and OvisE5334334) and linked to the *SPINK13* promoter OvisP1101635, forming a candidate GLI3-like–enhancer–promoter–*SPINK13* regulatory axis **(Fig. 4e; Extended Data Fig. 5a**). Considering the reported roles of *SPINK13* in male reproductive tract function, sperm maturation, and male fertility, this regulatory module provides a candidate mechanism through which a GLI3-like motif may contribute to tissue-specific regulation of male reproductive genes^52,53^.

Overall, this analysis provides a multi-omic framework for prioritizing tissue-associated GRNs, although the inferred links remain correlative and require perturbation or reporter assays for causal validation.

## Allele-specific regulatory variants

Allele-specific (AS) regulatory effects provide a direct approach for linking genetic variation with *cis-*regulatory activity while minimizing confounding from *trans-*acting effects and technical variation^54,55^. To systematically characterize *cis*-regulatory effects in the sheep genome, we performed AS analyses at 13,366,709 unique heterozygous SNPs across chromatin accessibility, histone modifications and transcriptomes. In total, we identified 145,407 SNPs exhibiting significant allelic imbalances (BH-adjusted *P* < 0.1 after binomial testing), including 48,536 from ATAC-seq, 4,295 from H3K4me1, 28,968 from H3K4me3, 7,183 from H3K27ac, 258 from H3K27me3 and 56,167 from RNA-seq data (**Fig. 5a; Supplementary Fig. 14a–c; Supplementary Table 21**). These AS SNPs were preferentially located near TSSs, upstream regions, 5′ UTRs, and exons (**Fig. 5b; Supplementary Fig. 14d-g**).

**Fig. 5.**
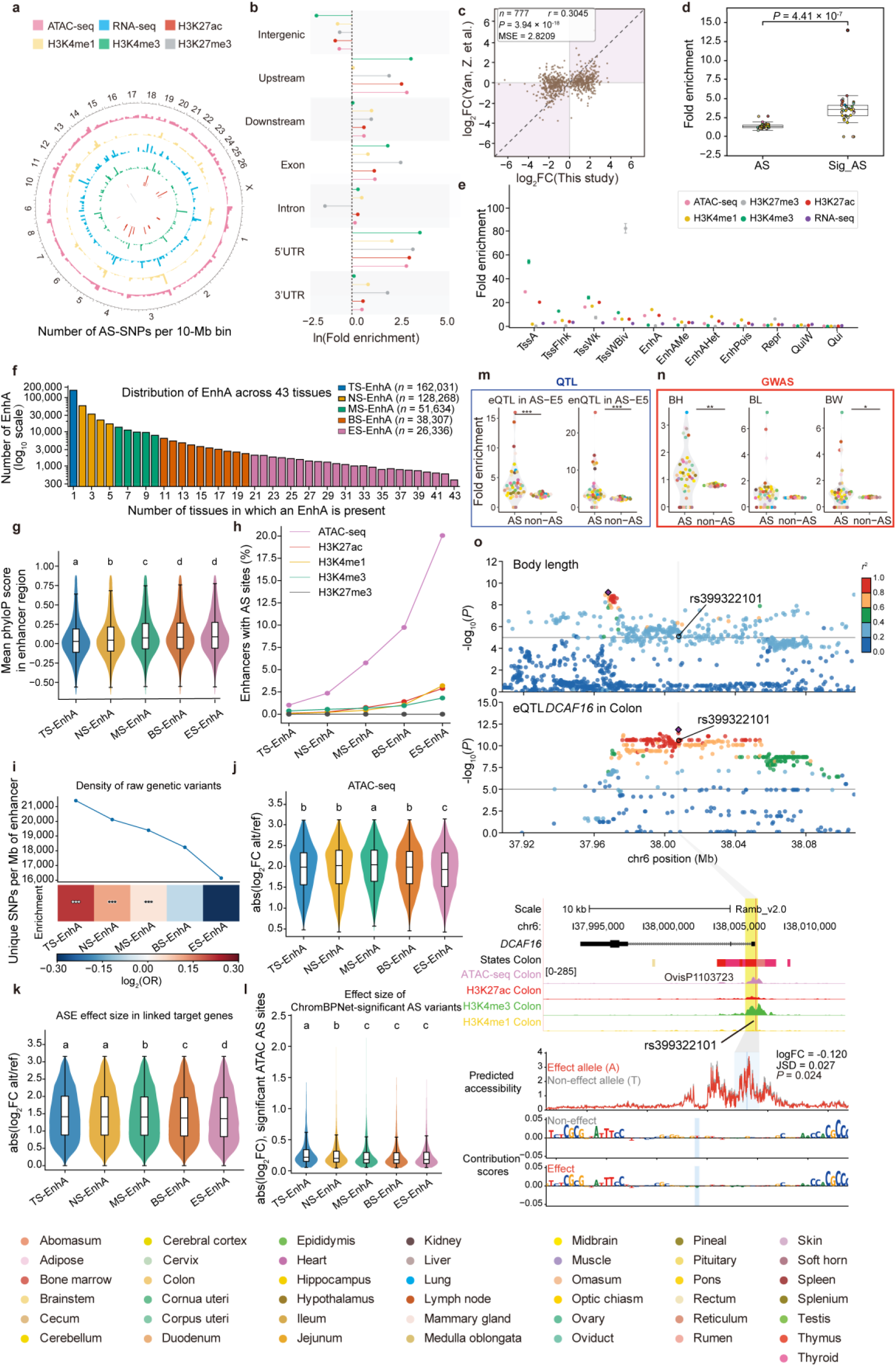
Genomic distribution, regulatory enrichment and trait relevance of allele-specific signals. a, Genome-wide distribution of allele-specific signals across assays. b, Enrichment of allele-specific variants across functional genomic annotations. c, Concordance of allelic effect sizes between this study and a published dataset. d, Enrichment comparison between all allele-specific variants and ChromBPNet-significant allele-specific variants. e, Enrichment of allele-specific signals across chromatin states. f, Distribution of active enhancers according to tissue-sharing breadth. g, Mean phyloP conservation scores across active-enhancer groups. h, Proportion of enhancers containing detected allele-specific sites across enhancer-sharing classes and molecular assays. i, Density and relative enrichment of raw genetic variants across active-enhancer groups. j, Effect-size distribution of ATAC-seq allele-specific variants across active-enhancer groups. k, Effect-size distribution of allele-specific expression sites in linked target genes across active-enhancer groups. l, Effect-size distribution of ChromBPNet-significant ATAC-seq allele-specific variants across active-enhancer groups. m–n, Enrichment of eQTLs, enQTLs and GWAS loci in allele-specific and non-allele-specific regulatory elements. o, Representative candidate regulatory variant at the *DCAF16* locus, integrating GWAS, eQTL, chromatin-state, epigenomic and ChromBPNet variant-effect tracks.

Allelic effects detected in RNA-seq data were positively correlated with those observed in ATAC-seq, H3K4me3 and H3K27ac profiles, indicating coordinated AS regulatory effects across molecular layers (**Supplementary Fig. 14h**). Significant correlations of signed allelic effect sizes between liver datasets from an independent sheep multi-omics study^56^ and this study support the reproducibility of the identified AS regulatory signals (**Fig. 5c**). ChromBPNet-based prediction further revealed that AS SNPs with significant predicted effects were preferentially enriched within TF motifs, suggesting that motif-disrupting variation contributes to AS regulatory activity^43^ (**Fig. 5d**). Chromatin state enrichment analysis showed that H3K4me3 AS SNPs were preferentially localized to promoter states, whereas H3K4me1 and H3K27ac AS SNPs showed enrichment in both proximal promoters and distal active enhancers (**Fig. 5e**).

We further classified EnhA elements into five groups according to the number of tissues in which they were active: TS-EnhA (1 tissue), NS-EnhA (2–5 tissues), MS-EnhA (6–10 tissues), BS-EnhA (11–20 tissues), and ES-EnhA (21–43 tissues) (**Fig. 5f; Extended Data Fig. 6a–c**). We observed a distinct pattern of regulatory variation associated with enhancer sharing. Although broadly shared enhancers exhibited an increased proportion of detectable AS SNPs, they showed reduced overall population-level SNP density and genomic variation, consistent with stronger purifying constraint on broadly pleiotropic regulatory elements^57,58^ (**Fig. 5g-i; Extended Data Fig. 6d–h**). Consistently, allelic effect sizes detected in ATAC-seq, H3K4me1 and H3K27ac profiles were lowest in ES-EnhA, and both ChromBPNet-predicted regulatory effects and AS expression effects decreased with increasing enhancer-sharing breadth (**Fig. 5j-l; Extended Data Fig. 6i–m**). Combined, these findings indicated that broadly active enhancers are more constrained and harbor variants with modest regulatory effects, whereas tissue-restricted enhancers are enriched for stronger context-dependent regulatory effects^8,59^.

Finally, to evaluate the relevance of AS regulatory variants for complex traits, we tested enrichment of molQTL and GWAS-associated loci within AS and non-AS regulatory elements. AS regulatory elements showed significant enrichment for eQTL and enQTL in SheepGTEx^27^, as well as GWAS loci associated with 34 complex traits^27^ (**Fig. 5m,n; Supplementary Fig. 15**). For example, the body-length-associated variant rs399322101 was also a colon eQTL near *DCAF16*, a CRL4 E3 ligase substrate receptor within the growth-associated *DCAF16– NCAPG–LCORL* locus in sheep and cattle^60,61^. ChromBPNet-based variant effect prediction further suggests that the effect allele A reduced local chromatin accessibility relative to the alternative allele T, providing a potential regulatory mechanism underlying this association (**Fig. 5o**).

## Integrating SheepEpimap with molQTL and GWAS signals in sheep

To evaluate the utility of the SheepEpimap for interpreting genetic associations, we integrated the atlas with 38 monogenic variants in OMIA^62^, 6,937 trait-associated QTL from AnimalQTLdb^63^, 19,054,283 SheepGTEx eQTL/enQTL^27^, and 21,531 GWAS signals associated with 34 complex traits^27^ (**Fig. 6a**). We found that 25 OMIA loci overlapped annotated regulatory elements, representing 65.8% of the analyzed loci, whereas 3,759 QTL from AnimalQTLdb overlapped regulatory elements, corresponding to 54.2% of all examined QTL (**Fig. 6b; Supplementary Tables 22 and 23**). For example, the 2-bp deletion chr2:204020069– 204020070delAG in OMIA overlapped a brain-specific enhancer, predicted to regulate *ALS2*, a gene whose loss of function causes juvenile-onset motor neuron disorders (**Fig. 6c**)^64,65^. This observation illustrates how SheepEpimap can provide regulatory context for interpreting trait-associated loci.

**Fig. 6.**
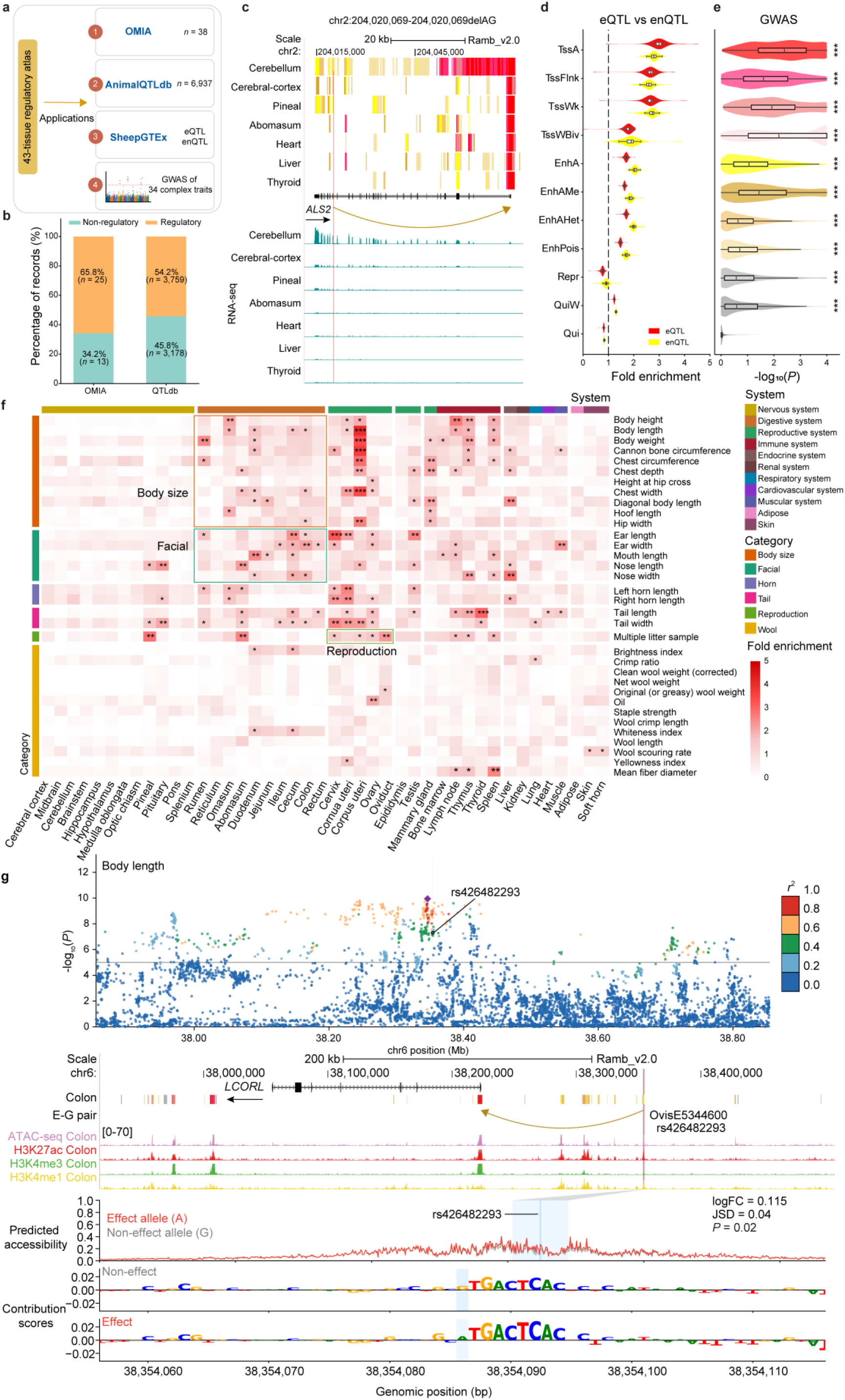
Application of the sheep regulatory atlas to variant and trait interpretation. a, Schematic overview of downstream applications of the 43-tissue regulatory atlas, including OMIA variants, Animal QTLdb records, SheepGTEx eQTLs and enQTLs, and GWAS signals for complex traits. b, Proportion of OMIA and QTLdb records overlapping regulatory and non-regulatory regions. c, Chromatin states and RNA-seq signals surrounding the candidate *ALS2* deletion associated with segmental axonopathy. d, Enrichment of sheepGTEx eQTLs and enQTLs across chromatin states. e, GWAS enrichment significance across chromatin states for complex traits. Asterisks denote nominal significance levels: \**P* < 0.05, \*\**P* < 0.01 and \*\*\**P* < 0.001. f, Heatmap showing GWAS enrichment in tissue-specific active enhancers across tissues and traits. g, Regulatory annotation and ChromBPNet allele-effect prediction for the body-length-associated variant rs426482293 at the *LCORL* locus, showing regional GWAS signals, chromatin-state annotation, enhancer–gene pairs, colon epigenomic signals, predicted accessibility and base-resolution contribution scores.

MolQTL and GWAS signals further supported the functional relevance of the SheepEpimap regulatory annotations. The SheepGTEx eQTL were preferentially enriched in promoter states, and enQTL were preferentially enriched in enhancer states (**Fig. 6d**). Integration of GWAS signals for 34 complex traits with the 11-state chromatin annotation revealed significant enrichment across multiple promoter and enhancer states (**Fig. 6e**). Focusing on tissue-specific EnhA elements, we observed multiple trait–tissue enrichment patterns that were consistent with known biological functions. For example, GWAS signals for body size and facial size were enriched in digestive tissues, signals for litter size were enriched in the uterus, ovary and oviduct, and associations for wool scouring rate were enriched in skin (**Fig. 6f; Supplementary Table 24**).

As a representative body-length locus, the associated variant rs426482293 was located within a colon enhancer (OvisE5344600) at the *LCORL* locus (**Fig. 6g**). *LCORL* has been repeatedly implicated in body size, body weight and growth-related traits in sheep and other livestock species^66,67^. ChromBPNet-based prediction indicated that the A allele was associated with increased local chromatin accessibility compared with the G allele, suggesting that rs426482293 represents a candidate regulatory variant potentially acting through a colon enhancer–*LCORL* axis (**Fig. 6g**). Similarly, we identified a wool-scouring-rate-associated variant, rs404999101, located within a skin enhancer (OvisE5072006) near *PIK3CD* (**Extended Data Fig. 7**). Together, these examples demonstrate how tissue-resolved regulatory annotations can refine the interpretation of trait-associated loci by nominating candidate regulatory elements, tissues, target genes and putative functional variants.

## SheepEpimap prioritizes candidate regulatory variants within selection signals

To investigate the regulatory architectures shaped by sheep domestication and environmental adaptation, we integrated SheepEpimap with candidate selective-sweep regions derived from SheepGTEx^27^ genotype data and Li et al^68^. across four historical and demographic comparisons. These included: modern Central East Asian (*n* = 1,104) versus modern European (n = 710) sheep; ancient CEA (n = 21) versus modern CEA (n = 1,104) sheep; ancient EUR (n = 31) versus modern EUR (n = 710) sheep; and domestic sheep (n = 738) versus wild Ovis orientalis (n = 72) (Fig. 7a). For the three domestic-sheep comparisons, candidate selective-sweep regions were defined using the top 1% of genome-wide empirical F_ST values. For domestic sheep versus wild Ovis orientalis, regions in the top 1% of both XP-CLR and *π*-ratio values were retained instead (Methods). Collectively, these regions contained 6,986 candidate variants, of which 98% were noncoding and 87% overlapped or flanked at least one tissue-specific regulatory region (**Fig. 7b**).

**Fig. 7.**
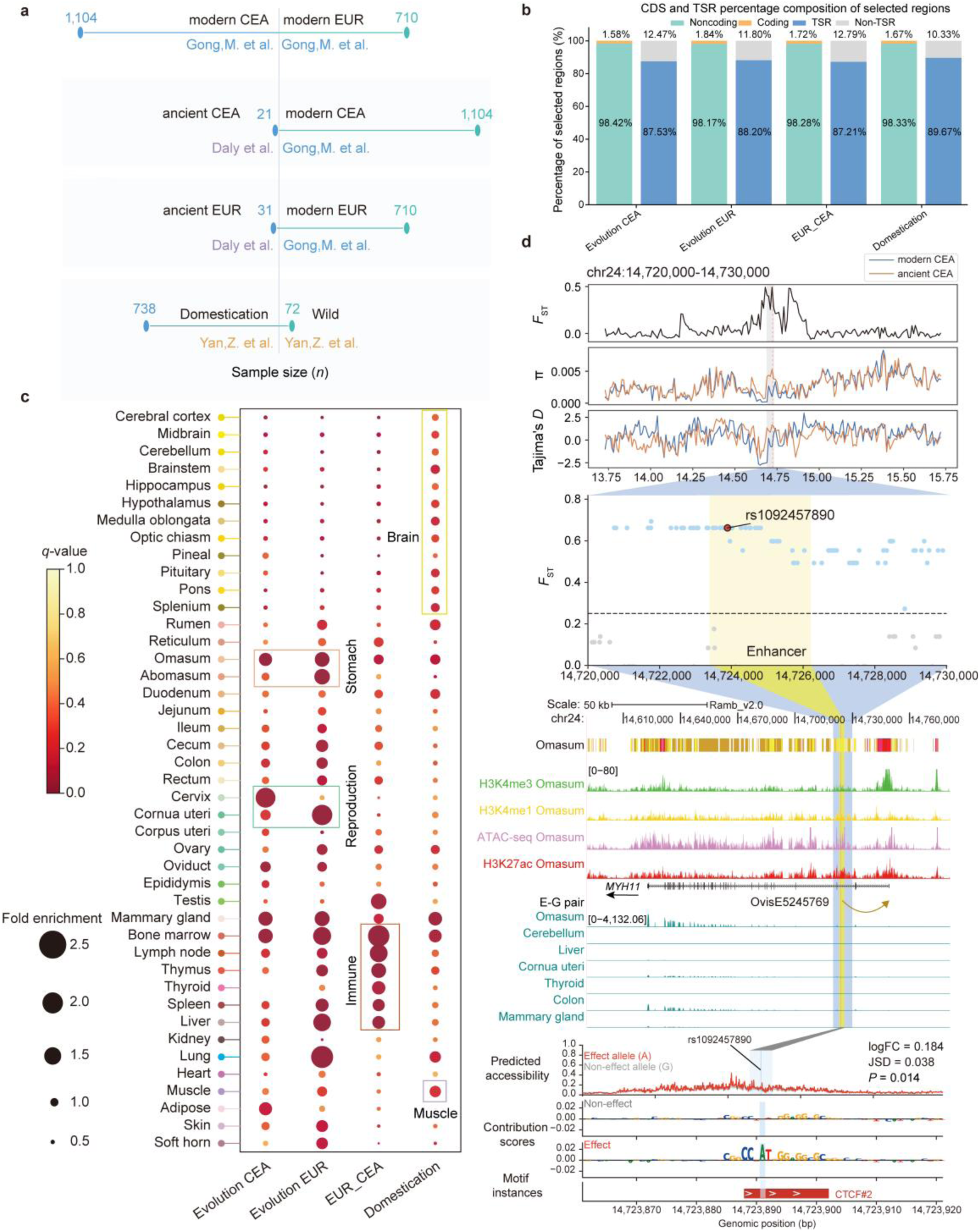
Regulatory annotation of candidate selected variants using the sheep regulatory atlas. a, Population groups, sample sizes and data sources used for four selection-signal comparisons: ancient versus modern Central and East Asian sheep, ancient versus modern European sheep, modern European versus modern Central and East Asian sheep, and domesticated versus wild sheep. b, Percentage composition of candidate selected regions according to coding status and overlap with tissue-specific regulatory regions across the four selection-signal classes. c, Enrichment of tissue-specific active enhancers (EnhA) across tissues and selection-signal classes. Dot size indicates fold enrichment, and color denotes the *q*-value. d, Regulatory annotation of the candidate selected variant rs1092457890 at the *MYH11* locus. Regional tracks show fixation index (*F*_ST_), nucleotide diversity (π) and Tajima’s D, followed by local *F*_ST_ values across the candidate enhancer OvisE5245769. Genome-browser tracks show chromatin states, epigenomic signals, enhancer–gene pairs, gene annotation and RNA-seq profiles. The lower panels show ChromBPNet-predicted accessibility, base-resolution contribution scores and motif instances for the effect and non-effect alleles. CEA, Central and East Asian; EUR, European; TSR, tissue-specific regulatory region.

We examined the enrichment of candidate selected variants within tissue-specific enhancers. Active enhancer (EnhA) states showed distinct enrichment patterns across tissues (**Fig. 7c**). Ancient-to-modern comparisons showed preferential enrichment in reproductive and digestive tissue enhancers, suggesting continued selection on physiological traits associated with fertility and metabolism during sheep improvement. In contrast, modern CEA–EUR differentiation was enriched in immune-related enhancers, consistent with divergent pathogen exposure and environmental adaptation between geographic populations. Domestic-versus-wild ancestral comparisons showed strong enrichment patterns in neural, neuroendocrine, and muscle-associated enhancers, consistent with the “domestication syndrome” hypothesis, in which early domestication involved selection on behavior, nervous system traits, and muscle-related phenotypes ^1,69,70^.

We next sought to connect these selective sweep signals with tissue-specific regulatory mechanisms. In the ancient-to-modern Asian comparison, selected variants were enriched in omasum-associated regulatory networks. The variant rs1092457890 overlapped an omasum enhancer predicted to interact with *MYH11*, encoding smooth-muscle myosin heavy chain 11, a key contractile protein involved in visceral smooth-muscle function and gastrointestinal motility^71^. Another selected variant, rs595082469 was located in the *GAA* promoter and overlapped an omasum eQTL in which *GAA* was the target gene. *GAA* encodes lysosomal acid α-glucosidase, an enzyme involved in lysosomal glycogen degradation, and mutations in this gene cause Pompe disease characterized by skeletal and cardiac muscle dysfunction^72^ (**Fig. 7d; Extended Data Fig. 8a**). Together with the genome-wide enrichment of highly shared E5 enhancers within the omasum, these observations suggest that regulatory variation affecting forestomach function may have contributed to dietary and metabolic adaptation during sheep domestication in Asian populations.

Similar regulatory patterns were observed in the European selection comparisons. In the ancient-to-modern European lineage comparison, three highly differentiated variants clustered within an abomasum enhancer linked to *CAPN5*. These variants overlapped with abomasum *CAPN5* eQTL, and ChromBPNet modeling predicted allele-dependent differences in local chromatin accessibility associated with these variants. *CAPN5* encodes calpain-5, a calcium-dependent cysteine protease involved in targeted cellular proteolysis and tissue remodeling^73,74^. These findings nominate CAPN5-associated abomasum regulatory activity as a candidate regulatory association in post-domestication adaptation of European sheep lineages (**Extended Data Fig. 8b**). Together, these examples illustrate that SheepEpimap can integrate population-genetic selection signals with tissue-resolved regulatory mechanisms to prioritize candidate regulatory elements and genes associated with domestication and adaptation.

## Translational analysis of SheepEpimap in human complex traits and diseases

To evaluate the translational potential of the SheepEpimap for interpreting the genetic architecture of human complex traits, we mapped sheep tissue-specific active enhancers (EnhA) to the human genome using liftOver^75^ and then partitioned heritability across 37 human GWAS traits using LDSC^76,77^. Cross-species projection recaptured biological expectation of tissue–trait pairs, including uterus-associated enrichment for human multiple-birth phenotypes, heart-associated enrichment for pulse rate and brain-associated enrichment for neuroticism score (**Fig. 8a**)^78–80^. These results indicate that tissue-specific regulatory landscape in sheep can capture evolutionarily conserved features relevant to human complex traits and further highlight the value of sheep as translational large-animal models for human reproductive biology, cardiovascular physiology, and neurological disorders^11,81,82^.

**Fig. 8.**
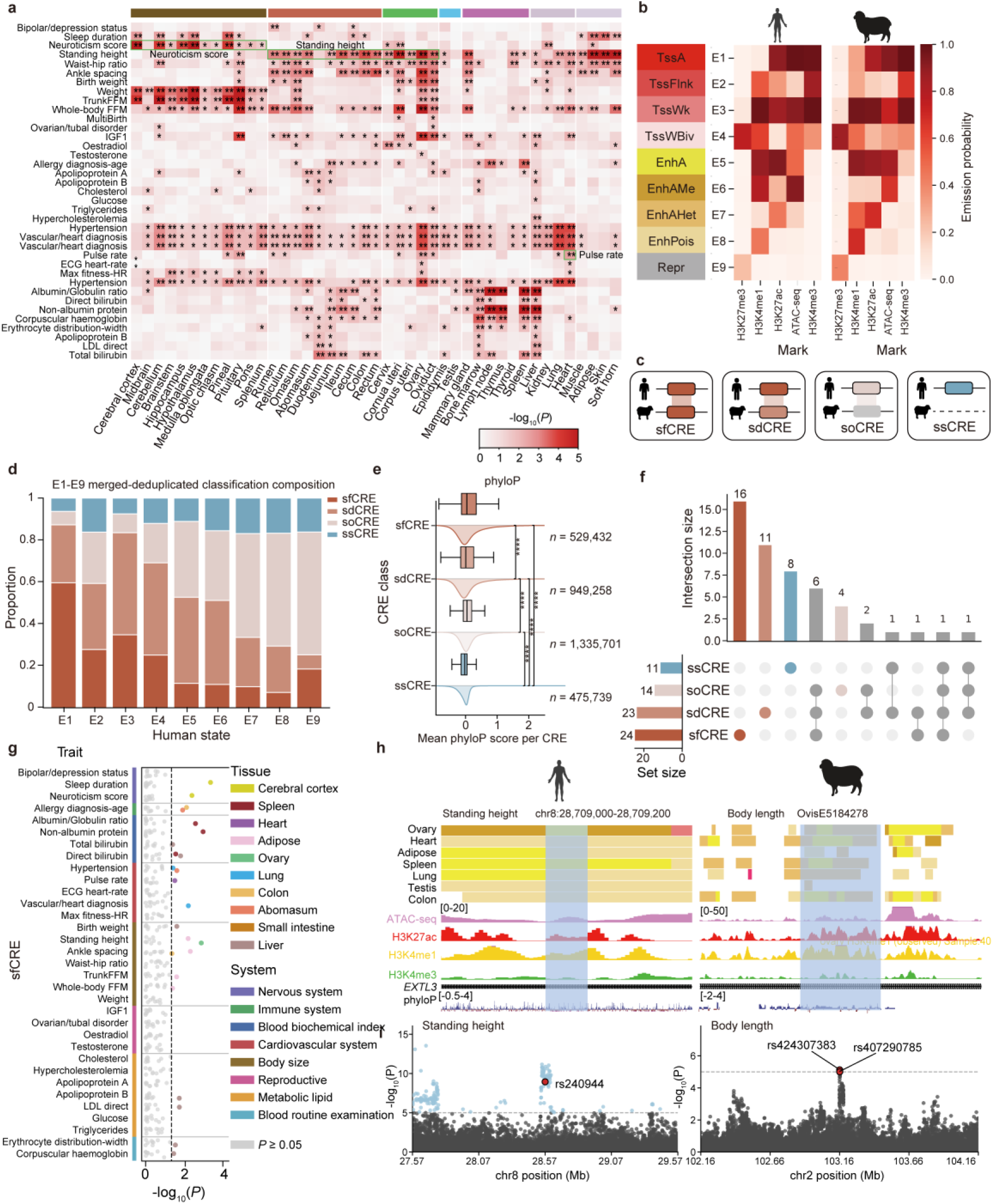
Cross-species conservation of sheep and human *cis*-regulatory elements informs human complex-trait architecture. a, Partitioned heritability enrichment of human GWAS traits in tissue-specific sheep EnhA annotations mapped to the human genome. Color intensity indicate −log_10_(*P*), and asterisks denote nominal significance levels. b, ChromHMM emission probabilities of matched chromatin states in human and sheep. Rows represent chromatin states, and columns represent chromatin accessibility and histone-mark assays. c, Classification scheme for human *cis*-regulatory elements based on sequence synteny and matched-tissue sheep chromatin states. d, Proportional composition of conserved and species-divergent *cis*-regulatory element classes across human chromatin states. e, Distribution of mean phyloP scores across *cis*-regulatory element classes. f, UpSet plot showing intersections of nominal trait–tissue enrichment signals across E5 *cis*-regulatory element conservation classes. g, Enrichment signals of sfCREs across human GWAS traits grouped by trait system. Colored points indicate nominally significant tissue–trait signals, grey points indicate non-significant tests, and the dashed line indicates nominal *P* = 0.05. h, Representative conserved regulatory locus showing matched human and sheep regulatory tracks for standing height and body length signals. i, Regional GWAS association signals at the corresponding human standing-height and sheep body-length loci.

To investigate the evolutionary basis of these cross-species enrichment patterns, we distinguished sequence conservation from conservation of regulatory activity. Across 12 matched sheep and human tissue pairs, we classified 3,290,130 human CREs into four distinct categories: 529,432 state-conserved functional CREs (sfCREs), 949,258 state-divergent functional CREs (sdCREs), 1,335,701 sequence-only conserved CREs (soCREs) and 475,739 species-specific CREs (ssCREs) (**Fig. 8b,c**). Across matched tissues, sfCREs represented a larger proportion of promoter states (E1–E4) than enhancer states (E5–E8) (36.4% versus 9.3%), consistent with stronger evolutionary constraint on promoter-associated regulatory elements (**Fig. 8d**). Among active enhancers, sequence-conserved CREs (sfCREs, sdCREs and soCREs), exhibited significantly higher phyloP, phastCons and GERP scores than ssCREs, with sfCREs showing the strongest sequence conservation (**Fig. 8e; Supplementary Fig. 16a,b**).

We next assessed human-trait heritability enrichment across the four CRE classes within tissue-specific EnhA elements. Of the 37 trait–tissue pairs tested, 24 (64.9%) showed significant heritability enrichment in sfCREs, consistent with previous reports^24^. This suggests that regulatory elements maintaining both sequence conservation and tissue-specific activity across mammalian species capture a substantial proportion of functional genetic signals underlying human complex traits (**Fig. 8f,g; Supplementary Fig. 16c-d; Supplementary Tables 26**).

Further locus-level comparative analyses identified conserved CREs overlapping association signals for human skeletal height and sheep body length. For the ovary–standing height association, the human height-associated variant rs240944 overlapped an ovary sfCRE syntenic to the sheep ovary enhancer OvisE5184278 at the *EXTL3* locus. Pathogenic *EXTL3* variants cause neuro-immuno-skeletal dysplasia characterized by short stature, supporting a potential role of this conserved locus in height-associated biology^83^ (**Fig. 8h**). For the pulse rate, rs9990137 overlapped a human heart sfCRE syntenic to sheep heart enhancer OvisE5158497, which was predicted to link to *SCN5A*, a cardiac sodium-channel gene essential for action-potential initiation and conduction^84^ (**Supplementary Figs. 17c** **and 18**). Similarly, the cortex– neuroticism association mapped to a cerebral cortex sfCRE associated with *CRHR1*, *MAPT* and *NSF*, genes implicated in neuroticism and brain-related traits (**Supplementary Figs. 17a** **and 18**).

Taken together, these results demonstrated that cross-species regulatory conservation can connect human trait-associated variants with tissue-matched regulatory landscape and prioritize candidate functional mechanisms. SheepEpimap therefore provides a comparative functional genomics ecosystem for investigating mammalian regulatory evolution, complex trait biology and translational disease models.

## Discussion

SheepEpimap integrates chromatin accessibility, four histone modifications and transcriptomic profiles from 43 adult sheep tissues, comprising 516 genome-wide datasets, to establish a systematic multi-tissue regulatory atlas of the sheep genome. Beyond defining an 11-state chromatin model, tissue-specific *cis*-regulatory elements, gene networks and allele-specific regulatory variation, this resource places chromatin states, tissue identity, gene-expression programs, regulatory sequence features and genetic variant interpretation within a unified framework. To facilitate broad community access and exploration of these regulatory annotations, we have developed a dedicated web portal at https://genome.ucsc.edu/s/mengzhu/SheepEpimap that enables interactive visualization and query of sheep regulatory elements across tissues. In doing so, SheepEpimap moves the sheep non-coding genome beyond a static catalog of regulatory DNA elements towards a tissue-informed framework for candidate regulatory mechanism discovery^19,20,85^. Our analyses indicate that tissue-specific active enhancers represent a major layer of regulatory specialization across sheep tissues. The biological processes, TF motifs and human or mouse orthologous phenotypes associated with tissue-specific EnhA-linked genes were concordant with the corresponding tissue functions, supporting an important role for enhancer activity in encoding tissue identity and functional specialization^7,86^.

Sequence-to-function modeling with ChromBPNet further increased the interpretability of this atlas^43^. ChromBPNet recovered canonical and *de novo* motifs and resolved sequence determinants of chromatin accessibility at base-pair resolution, providing a basis for scoring the potential regulatory effects of genetic variants at scale. This strategy complements conventional motif analysis and provides a sequence-level view of sheep *cis*-regulatory grammar¹². Gene regulatory networks constructed from motifs, enhancers, promoters and genes further connect regulatory elements to putative upstream transcription factors and downstream candidate target genes, offering a structured framework for prioritizing biologically plausible regulatory pathways. Allele-specific analyses add a variant-centered layer to this framework, showing that allelic imbalance can provide an informative signal for identifying *cis*-acting regulatory variants and prioritizing putative functional non-coding variation^87^. By integrating OMIA^62^ records, Animal QTLdb^63^ annotations, sheepGTEx^27^ eQTLs and enQTLs, complex-trait GWAS signals and selection scans, SheepEpimap helps refine association intervals into candidate regulatory elements, relevant tissue contexts and putative effector genes. Sheep– human comparative epigenomic analyses further suggest that sequence-conserved regulatory elements may carry shared regulatory information across species and provide a useful reference for interpreting the genetic architecture of human complex traits and disease-associated non-coding variation^88–92^. Together, SheepEpimap provides a systematic resource for investigating the contribution of non-coding regulatory variation to complex traits in sheep, and its importance for understanding the biology of domestication and breed differentiation, and for enhancing molecular breeding, while also offering a tissue-resolved reference for comparative functional genomics and large-animal models of human disease.

Several limitations should be considered when interpreting and applying SheepEpimap. First, the atlas is based primarily on bulk adult tissues and a limited number of individuals, which constrains its ability to resolve cell-type heterogeneity within complex tissues, developmental dynamics and population-scale regulatory diversity. Future integration of single- cell and spatial epigenomic and transcriptomic datasets will help refine tissue-level annotations to cell-type resolution and capture cellular heterogeneity and dynamic state transitions during development, physiological state change or disease progression^93–98^. Second, the enhancer– gene links, gene regulatory networks and ChromBPNet-predicted allelic effects generated in this study are inferred primarily from correlations, chromatin features and sequence models. They should therefore be interpreted as prioritized candidate regulatory relationships rather than direct evidence of causality. Larger sheep GWAS, fine-mapping datasets and molecular QTL maps will be needed to improve the resolution with which regulatory variants can be linked to complex traits. In parallel, CRISPR perturbation^99^, STARR-seq^100^, massively parallel reporter assays^101^ and allele-specific reporter systems^102^ will be important for functionally testing prioritized enhancers, TF motifs and candidate variants. Future integration of SheepEpimap with haplotype-resolved pangenomes^103^, structural-variation maps^3^, long-read transcriptomes^104^, single-cell omics and perturbation screens^105^ will enable more precise identification of causal regulatory variants and their target genes. More broadly, SheepEpimap offers new insight into how the mammalian regulatory genome is remodeled by domestication, human-directed selection and adaptive microevolution across ecologically diverse environments. It also substantially enhances the value of the sheep as a large-animal model for human physiology and disease.

## Online Methods

### Sample collection and animal ethics

Adult Small-Tailed Han sheep, including two rams and two ewes, were used for tissue collection. The ewes were subjected to estrus synchronization and slaughtered 24 h after sponge withdrawal. Forty-one tissues were collected from ewes, including cerebral cortex, hypothalamus, cerebellum, midbrain, splenium, pineal gland, pituitary, optic chiasm, hippocampus, pons, medulla oblongata, brainstem, rumen, reticulum, omasum, abomasum, duodenum, ileum, jejunum, cecum, colon, rectum, heart, liver, spleen, lung, kidney, muscle, adipose, bone marrow, thymus, skin, soft horn, lymph node, thyroid, ovary, corpus uteri, cornua uteri, cervix, oviduct and mammary gland. Testis and epididymis were collected from rams. In total, 43 tissue types were obtained. All tissues were dissected according to FAANG tissue-collection standards, snap-frozen in liquid nitrogen immediately after collection and stored at −80 °C until use. All animal procedures were approved by the Animal Ethics Committee of the Institute of Animal Sciences, Chinese Academy of Agricultural Sciences, under approval number IAS2024-129.

### Sequencing data processing and quality control

ATAC-seq and RNA-seq datasets were processed using the FAANG functional annotation pipeline developed at the University of California, Davis. CUT&Tag data were analyzed following the method described by Kaya-Okur et al.^106^. All sequencing reads were aligned to the sheep reference genome ARS-UI_Ramb_v2.0 (GCF_016772045.1). Data quality was assessed using the ENCODE cross-correlation QC^107^ tool phantompeakqualtools/run_spp.R. Samples were considered to have failed quality control if QualityTag < 0, if QualityTag = 0 and RSC < 1.1, or if RSC < 0.8. Adapter sequences and low-quality bases were removed using Trim Galore (v0.6.5) ^108^.

RNA-seq reads were aligned to the reference genome using STAR (v2.7.11a)^109^. ATAC- seq reads were aligned using Bowtie2 (v2.5.3)^110^. Alignments with MAPQ < 30 were removed using SAMtools (v1.18)^111^. Gene-level read counts from RNA-seq data were quantified using HTSeq-count (v2.0.4)^112^, and gene expression was normalized using edgeR (v4.0.2)^113^ and StringTie2 (v2.2.1)^114^. CUT&Tag PCR duplicates were removed using Picard (v3.1.1), and peaks were called using MACS2 (v2.2.9.1)^115^. Public Hi-C data were downloaded from Yan et al^56^ and processed using Juicer (v1.6)^116^ for topologically associating domain (TAD) identification.

Public WGBS datasets for muscle, ovary, skin and liver were downloaded from EBI^117–122^. Adapter trimming was performed using Trim Galore (v0.6.6), and preprocessing was performed using Bismark (v0.24.2)^123^. BAM files were converted to compressed beta and pat formats using wgbs_tools (v0.2.2)^124^, and homogeneous methylation blocks were identified using the segment command. Blocks containing fewer than four CpG sites were excluded from downstream analyses. For each methylation block, the mean methylation level across all CpG sites was calculated from the corresponding beta file.

## Chromatin-state annotation and genomic-feature enrichment

Chromatin states across 43 sheep tissues were inferred using ChromHMM (v1.26)^32^. For each tissue, five epigenomic features were used as input: ATAC, H3K4me3, H3K4me1, H3K27ac and H3K27me3. Low-quality datasets were imputed using ChromImpute (v1.0.5)^28^. Briefly, ComputeGlobalDist was first used to calculate global distances among datasets; GenerateTrainData was used to generate training features; Train was used to train mark- and sample-specific predictors; and Apply was used to generate imputed signal tracks. Following the original ChromImpute study^43^, Pearson correlation between observed and predicted read counts was used to evaluate imputation performance, and datasets meeting this analysis criterion were retained for downstream analyses.

For chromatin-state modelling, ChromHMM LearnModel was used to train models with 2–20 states. Following Gorkin et al.^125^, CompareModels was used to calculate emission-probability correlations among models. The optimal number of states was defined as the point at which the median emission-probability correlation reached a plateau in two or more replicate analyses. An 11-state model was selected for the sheep epigenome, and state labels were assigned following the approach of Jason et al.^126^. To enable sheep–human comparison, the same workflow was applied to 111 human epigenomes from ENCODE^127^ and Roadmap Epigenomics^7^.

## Identification and functional analysis of tissue-specific chromatin states

Chromatin-state intervals from 43 tissues were merged using BEDTools (v2.29.2)^128^ to generate a non-redundant set of regulatory elements. Chromatin-state variability and state transitions among tissues were then calculated. To analyse active enhancers across tissue types, the regulatory-region overlap across tissues (RRAT) was computed. For each tissue, an RRAT region was assigned a value of 1 if at least one EnhA overlapped the region and 0 otherwise. Tissue-specific regulatory elements (TSRs) were identified for all chromatin states except the quiescent state (Qui) using RRAT.

The GO enrichment analysis of TSR target genes was performed using clusterProfiler (v4.4.1)^129^. Motifs enriched in tissue-specific EnhA regions were identified using HOMER (v4.11)^130^, with FDR < 0.05 used as the significance threshold. Enriched motifs related to tissue function were defined as candidate motifs for each tissue. To evaluate the relationship between sheep tissue-specific EnhAs and human or mouse phenotypes, sheep tissue-specific EnhAs were converted to human genome coordinates using liftOver^75^. The converted human genomic intervals were then analyzed using GREAT^131^ for human and mouse phenotype enrichment, with parameters set to proximal upstream 2 kb, downstream 1 kb and distal extension 3 kb.

## Enhancer–gene pair prediction and functional analysis

Enhancer–target gene pairs were predicted by integrating H3K27ac, ATAC-seq and RNA-seq data across 43 sheep tissues. E5 enhancer regions annotated by ChromHMM in each tissue were merged to generate a non-redundant candidate enhancer set. H3K27ac and ATAC-seq read counts were quantified within candidate enhancer regions and normalized to CPM using edgeR. Gene expression was quantified as RNA-seq TPM.

For each gene, candidate enhancers located within ±500 kb of the TSS were retained. Pearson correlation coefficients between enhancer H3K27ac or ATAC-seq signal and gene expression across 43 tissues were calculated, and P values were adjusted using the Benjamini– Hochberg method to obtain q values. To determine the correlation threshold for H3K27ac enhancer–target gene pairs, ATAC-derived links were used as the reference. Accuracy, precision, recall, F1 score, Jaccard similarity and retained pair number were evaluated for H3K27ac- derived links across different Pearson’s *r* thresholds, and Pearson’s *r* = 0.3 was selected as the final threshold. High-confidence H3K27ac enhancer–target gene pairs were defined as enhancer–gene pairs with *q* value ≤ 0.05 and Pearson’s *r* > 0.3.

## ChromBPNet modeling of regulatory sequence features

ChromBPNet (v1.0)^43^ was used as a sequence-to-function model to learn regulatory sequence syntax of chromatin accessibility across 43 sheep tissues. Open chromatin regions (OCRs) with blacklist regions removed were used as input. OCRs were defined from replicate ENCODE narrowPeak calls, filtered using IDR (v2.0.3)^132^, required to show ≥50% reciprocal overlap between replicates and restricted to canonical chromosomes.

Chromosome-level data splits were generated using chrombpnet prep splits, with chr1, chr3 and chr6 used for testing, chr8 and chr20 for validation, and all remaining chromosomes for training. Model training was performed in two steps: first, a bias-only model was trained to capture Tn5 insertion bias; second, a bias-factorized accessibility model was trained to model chromatin accessibility after Tn5-bias correction. chrombpnet pred_bw tool was used to generate base-resolution bias-only and bias-corrected chromatin-accessibility predictions over tissue OCRs.

Following ChromBPNet standards^43^, prediction accuracy was evaluated using Pearson correlation, Spearman correlation and Jensen–Shannon distance (JSD). Models for which the Pearson correlation between observed and predicted read counts reached the analysis standard were retained for downstream analyses. ChromBPNet-derived contribution scores were then used to identify *de novo* motifs and generate a non-redundant cross-tissue motif set. Motif-centred footprint analyses were performed on motif instances using ATAC-seq cleavage tracks corrected by TOBIAS ATACorrect^47^. Statistical support was assessed using a matched-length accessible-region empirical null, and motif instances with nominal *P* < 0.05 were defined as footprint-supported motif instances.

## Construction of gene regulatory networks

To infer putative upstream and downstream regulatory relationships of enhancers, enhancer–gene pairs, promoter annotations and transcription-factor motif information were integrated to construct tissue-resolved gene regulatory networks. For the HOMER-based network, tissue-associated functional motifs were first extracted from HOMER motif-enrichment results for each tissue. Motif occurrences were scanned in enhancer regions using MEME FIMO (v5.5.9)^133^ with parameters --max-stored-scores 5000000 --thresh 0.001. FIMO enhancer-overlapping motif hits were used to define candidate TF-binding events and to connect TF motifs with motif-bound enhancers.

Enhancers were connected to candidate target genes using the predicted enhancer–gene pairs. For each tissue, motif-bound enhancers were matched with enhancer–gene pairs from the corresponding tissue, and only motif-bound enhancers with candidate target genes were retained to establish TF motif–enhancer–gene relationships. To include promoter-mediated regulatory paths, each target gene was further matched to its promoter annotation, defined as the 2-kb region upstream of the gene TSS. Motif-bound enhancers, enhancer-linked genes and corresponding promoter annotations were integrated to construct gene regulatory modules.

Network nodes were defined as four classes: motif, enhancer, promoter and gene. Network edges represented motif–enhancer, enhancer–gene and promoter–gene relationships. For each tissue–motif combination, node and edge tables were generated and then merged to construct the global regulatory network. HOMER-derived and ChromBPNet-derived motif sets were integrated using the same network-construction logic. For each regulatory pair, the method label was retained for comparison. Regulatory pairs identified by both methods were defined as shared pairs, whereas pairs identified by only one method were defined as HOMER-specific or ChromBPNet-specific pairs. Network nodes were assigned to tissue modules according to tissue labels, and representative tissue-associated subnetworks were extracted for global network visualization, method-specific comparison and locus-level regulatory interpretation.

### Identification of allele-specific events

To identify allele-specific regulatory and transcriptional events, a phased genotype reference dataset was first constructed for each individual. SNP calling was performed using the GATK (v4.6.2.0) workflow^134^. Single-sample GVCF files were generated using GATK, and joint genotyping was performed with GenotypeGVCFs to obtain raw VCF files. Raw VCF files were split by chromosome, followed by genotype imputation, haplotype phasing and quality-control filtering using Beagle (v5.4)^135^. The resulting phased VCF files were used as the genotype reference for allele-specific analyses.

To reduce reference-mapping bias, RNA-seq, ATAC-seq and CUT&Tag BAM files were corrected using WASP (v0.16.3) ^136^. Reads overlapping heterozygous SNPs were identified, the allele carried by each read was swapped to the alternative allele, and allele-swapped reads were realigned to the reference genome. Reads that failed to realign to their original genomic positions were removed. The WASP-corrected BAM files were used to count allelic reads.

At heterozygous SNPs, allelic read counts were quantified using GATK ASEReadCounter with default parameters. ASEReadCounter outputs were filtered using default criteria related to mapping quality, base quality, sequencing depth, overlapping paired-end reads and indels. For each site, allelic imbalance was tested using a two-sided binomial test under the null hypothesis of equal representation of the two alleles, with the expected allelic proportion set to 0.5. Sites were evaluated for significance only if both alleles were supported by at least three reads and the minor-allele fraction was ≥ 0.10. The *P* values were adjusted using the Benjamini–Hochberg method^137^, and sites with BH-FDR < 0.10 were defined as significant allele-specific events.

## Regulatory annotation and enrichment analysis of GWAS, QTL and OMIA variants

To evaluate the relationship between trait-associated variants and sheep regulatory elements, the 11-state chromatin annotation was integrated with GWAS summary statistics for 34 economically important complex traits in sheep^27^. These traits included body weight, body size, reproductive traits and wool-related traits (Supplementary Table 24). For each trait, GWAS results from multiple sheep populations were meta-analyzed using METAL (v2020-05-05)^138^. Given the limited statistical power of the available GWAS datasets, variants with *P* < 1 × 10⁻⁵ were defined as suggestive GWAS loci for downstream enrichment analyses.

Enrichment of significant GWAS loci in the 11 chromatin states and tissue-specific enhancers was evaluated using a genotype cyclical permutation test with 10,000 permutations. Differences in enrichment among chromatin states were assessed using a two-sided *t*-test, with the quiescent state “11 Qui” used as the reference state.

Sheep disease- and phenotype-associated variants were downloaded from OMIA^62^ and Animal QTLdb^63^ and used to evaluate overlap between known disease, phenotype or QTL records and regulatory elements. Genomic interval overlaps between OMIA or Animal QTLdb records and regulatory elements were determined using the intersect function in BEDTools (v2.29.2). Records overlapping regulatory elements were defined as regulatory-element-associated records and used for downstream locus-level interpretation.

To analyze the relationship between molecular QTLs and chromatin states, eQTL and enQTL loci were obtained from sheepGTEx and intersected with the 11-state chromatin annotation. For each QTL class, fold enrichment in each chromatin state was calculated as (C/A)/(B/D), where C is the number of QTL loci overlapping a given chromatin state, A is the total number of QTL loci, B is the genomic length covered by that chromatin state and D is the total analyzable genomic length.

## Identification and regulatory enrichment analysis of selection signals

Selection-signal data for four population comparisons were obtained from SheepGTEx and Li et al.^68^: modern Asian versus modern European sheep (EUR_CEA); ancient versus modern Asian sheep (Evolution CEA); ancient versus modern European sheep (Evolution EUR); and wild versus domestic sheep (Domestication). To identify genomic regions potentially affected by selection, pairwise fixation index (*F*_ST_) values were calculated for each comparison using 21,542,313 common SNPs from the SheepGTEx genotype reference panel with MAF ≥ 0.05. Pairwise *F*_ST_ was calculated using VCFtools (v0.1.16)^139^ with a 10-kb sliding window and 10-kb step size. For each population comparison, windows in the top 1% of the genome-wide empirical *F*_ST_ distribution were defined as candidate selective-sweep regions. For the wild-versus-domestic comparison, XP-CLR and nucleotide-diversity ratio (*π* ratio) analyses were performed as described by Li et al.^68^. XP-CLR scans used 0.5-cM windows spaced at 2- kb intervals, with up to 200 SNPs per window, and regions in the top 1% of both XP-CLR and ln(*π*_wild_/*π*_domestic_)/ln(2) values were retained as candidate selective sweeps.

Adjacent candidate regions were merged using BEDTools (v2.29.2)^128^, and candidate selective-sweep regions were functionally annotated using ANNOVAR^140^. To assess the robustness of selection signals, nucleotide diversity (*π*) and Tajima’s *D* were also calculated using VCFtools with a 10-kb window and 10-kb step size.

Regulatory enrichment of candidate-selected regions was assessed using the Genomic Association Tester (GAT)^141^. Candidate selective-sweep regions were tested for enrichment in chromatin states and in tissue-specific enhancer regions across 43 tissues. GAT simulations were repeated 10,000 times to estimate empirical *P* values. These *P* values were adjusted using the Benjamini–Hochberg method in R, and enrichment significance was assessed using the adjusted FDR.

## Cross-species CRE classification and heritability enrichment

Human chromatin states were re-annotated using ChromHMM with the same 11-state model and epigenomic inputs as used for sheep. Nine states with consistent cross-species emission probability profiles were retained for downstream analysis: E1–E9, corresponding to TssA, TssFlnk, TssWk, TssWBiv, EnhA, EnhAMe, EnhAHet, EnhPois and Repr. The E10 state, representing a species-divergent state, and the E11 state, corresponding to Qui, were excluded. For 12 matched human–sheep tissue pairs—adipose–adipose, colon–colon, cortex– cerebral cortex, heart–heart, liver–liver, lung–lung, muscle–muscle, ovary–ovary, small intestine–jejunum, spleen–spleen, stomach–abomasum and testis–testis—human chromatin intervals from each of the nine retained states (E1–E9) were converted from hg38 to sheep V3 genome coordinates using liftOver. The chain file hg38ToGCF_016772045.2 was used with minMatch = 0.1. Converted intervals were compared with merged sheep chromatin-state intervals in the matched tissue. Regions matching the same chromatin state in sheep were defined as state-conserved functional CREs (sfCREs). Regions matching any other E1–E9 state were defined as state-divergent functional CREs (sdCREs). Regions that did not match any of the E1–E9 states were defined as sequence-only conserved CREs (soCREs). CREs that failed liftOver conversion were defined as species-specific CREs (ssCREs). For the promoter states (E1–E4), interval midpoint overlap was used for classification; for the non-promoter states (E5– E9), BEDTools intersect -f 0.5 was used as the minimum overlap threshold.

Average phyloP, phastCons and GERP scores for the four CRE classes were calculated using bigWigAverageOverBed^142^, defined as the per-base average score across covered bases. Pairwise comparisons among the four CRE classes were performed using two-sided Mann– Whitney *U* tests, and *P* values were adjusted using the Benjamini–Hochberg method.

For each CRE class, tissue-specific heritability enrichment was analyzed using ldsc.py -- h2-cts in LDSC (v1.0.0)^77^. Sheep tissue-specific active enhancers were first converted to human genome coordinates using liftOver and then analyzed using ldsc.py --h2-cts to estimate partitioned heritability enrichment. The analysis used 1000 Genomes Phase 3 European allele frequencies, HM3 noMHC weights and the full baselineLD model, with the --overlap-annot option enabled. Enrichment *P* values were reported without multiple-testing correction.

## Data availability

The raw CUT&Tag data for H3K4me3, H3K4me1, H3K27ac and H3K27me3, together with the raw ATAC–seq and RNA-seq data generated across 43 sheep tissues, have been deposited in the NCBI Sequence Read Archive under BioProject accession PRJNA1237432. Processed datasets, including chromatin-state annotations, peak calls, bigWig signal tracks, enhancer–gene pairs, genome-wide transcription factor motif-instance annotations and related regulatory annotations, are available for visualization and download through the UCSC Genome Browser (https://genome.ucsc.edu/s/mengzhu/SheepEpimap). All other publicly available datasets analyzed in this study, including Hi-C, WGBS, SheepGTEx molecular QTL, OMIA, Animal QTLdb, ENCODE, Roadmap Epigenomics and human GWAS datasets, are available from the repositories and source publications cited in the Methods.

## Code availability

All custom scripts used for sequencing-data processing and quality control, epigenome imputation and chromatin-state annotation, regulatory-element characterization, enhancer– gene linking, super-enhancer and functional-enrichment analyses, ChromBPNet modeling and motif and footprint analyses, gene network construction, allele-specific analysis, integration of molecular QTL and GWAS signals, selection-sweep analysis, and cross-species comparative analyses are available through the SheepEpimap GitHub repository (https://github.com/SheepEpimap/sheep).

## Acknowledgements

This work was supported by grants from National Key R&D Program of China (2023YFF1001800 and 2022YFF1000100 to Z.P.), the National Natural Science Fund for Excellent Young Scientists Fund Program (Overseas) (2024-HY-01 to Z.P.), the Agricultural Science and Technology Innovation Program of China (CAAS-ZDRW202502 and ASTIP- IAS13 to M.C.), the Earmarked Fund for China Agriculture Research System of MOF and MARA (CARS-38-02 to M.C.), the China Agriculture Research System of MOF and MARA (CARS-38 to Z. Zhang), the Program of Anhui Provincial Key Laboratory of Livestock and Poultry Product Safety (AHXM26-01 to K.Z.), the Hainan Provincial Postdoctoral Research Funding (BH443441 to Z.M.). We thank all researchers who contributed to this study. We also acknowledge the Institute of Animal Science, Chinese Academy of Agricultural Sciences, Beijing, for providing access to high-performance computing resources.

## Author contributions

Zhang, M.C., D.G., Z.P. and L.F. conceived and designed the study. Z.M., S. Zhang, Hao Y., X.D., Q.M., Hao L., G. Zhang, X.H., Jianning H., X.G., S.N., Xiaoxu Z., M. Shan, J.Y., Xinyue L., Huaiqiang Y., Yangshuo H., Yuchen H., S.T., Xianfeng W. and Xiaosheng Z. collected samples and generated data. Z.M., Hao Y. and Q.M. prepared the CUT&Tag, ATAC–seq and RNA-seq libraries. Z.M. processed and analyzed the CUT&Tag, ATAC–seq and RNA-seq data and performed epigenome imputation, chromatin-state annotation and identification of tissue-specific regulatory elements. S. Zhang identified enhancer–gene pairs. L.W. and Xinyi L. identified super-enhancers. S. Zhang and F.L. performed ChromBPNet analyses and constructed the gene regulatory networks. L.W., Jiayi H. and P.Z. performed allele-specific analyses. X.S. and S. Zhang curated and processed the GWAS summary statistics and integrated the regulatory atlas with allele-specific, QTL and GWAS signals. Z. Zhang and M.G. performed the selection-signal analyses. Z. Zhuang contributed to the cross-species comparative analyses. Xudong W., Y.Y., K.Z., M.W., X. Lu, H. Zhou, G. Zhao, P.Z., Y.L., C.R., Xiaosheng Z., Y.J., W. Wu, R.X., R.P.M.A.C., O.M., D.E.M., K.G.D., E.L.C., L.F., Z. Zhang, M.C., D.G. and Z.P. contributed to the interpretation of results and manuscript discussions. Z.M., J.L., M.W. and X. Lu developed the SheepEpimap web portal. L.F., Z.M., Z. Zhang, M.C., D.G. and Z.P. wrote the manuscript. All authors reviewed, revised and approved the final version.

## Competing interests

The authors declare no competing interests.

---

**Extended Data Fig. 1.**
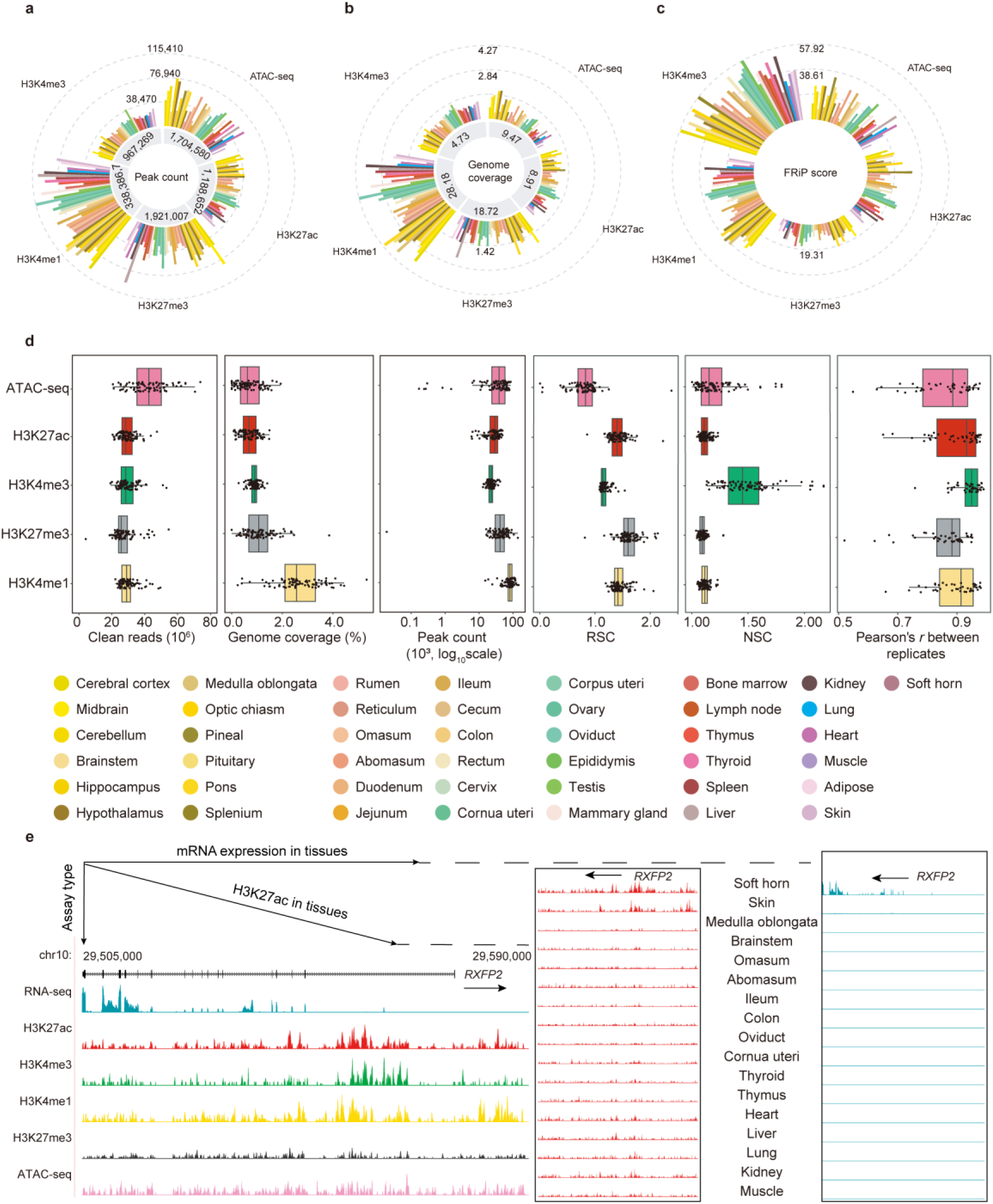
Overview and quality metrics of the integrated sheep multi-omics datasets. a, Peak counts for ATAC-seq and CUT&Tag profiles of H3K4me3, H3K27ac, H3K27me3 and H3K4me1 across tissues. b, Genome coverage of epigenomic peaks across tissues and assays. b, Fraction of reads in peaks (FRiP) scores for ATAC-seq and CUT&Tag datasets. d, Summary of dataset quality metrics, including clean reads, genome coverage, peak count, relative strand cross-correlation (RSC), normalized strand cross-correlation (NSC) and Pearson’s correlation between biological replicates. Points represent individual datasets and are coloured by tissue. e, Epigenetic signal at the RXFP2 locus according to different assays and in different tissues. Vertical scales were set to 0–50 for RNA-seq, H3K27ac and H3K4me3, 0–30 for H3K4me1 and ATAC-seq, and 0–80 for H3K27me3.

**Extended Data Fig. 2.**
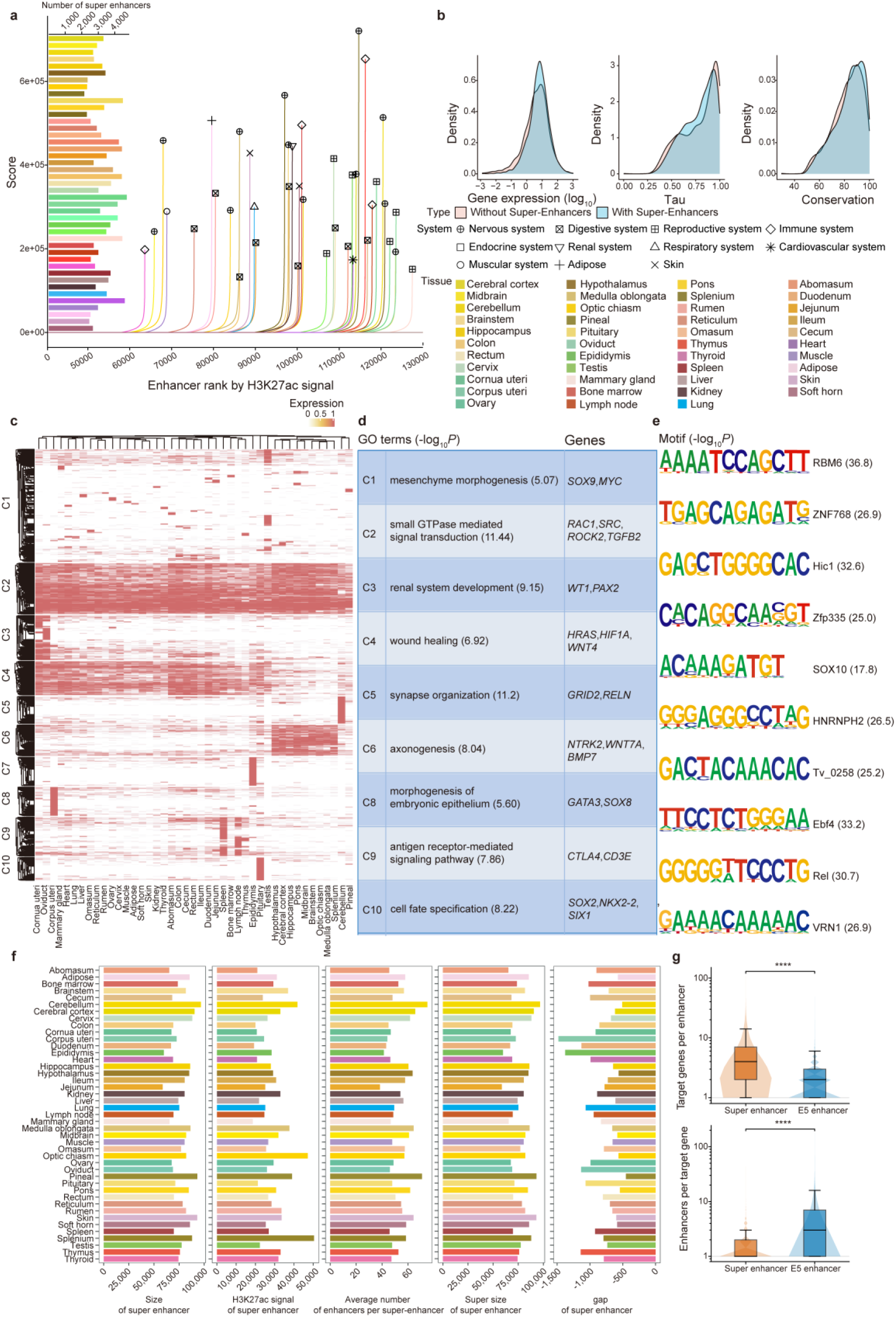
| Super-enhancer annotation and characterization. a, H3K27ac-based enhancer ranking curves and numbers of super-enhancers across tissues. b, Density distributions of gene expression, tissue specificity (Tau) and conservation for genes with and without super-enhancers. c, Heatmap of super-enhancer activity across tissues, grouped into ten clusters. d, Gene Ontology biological process terms and representative genes for the super- enhancer clusters. e, Motif enrichment for the super-enhancer clusters. f, Tissue-level summaries of super-enhancer size, H3K27ac signal, average number of constituent enhancers per super-enhancer, summed super-enhancer size and inter-enhancer gap size. g, Distributions of target genes per enhancer and enhancers per target gene for super-enhancers and E5 enhancers. P values were calculated using two-sided Mann–Whitney U tests; **** *P* < 0.0001.

**Extended Data Fig. 3.**
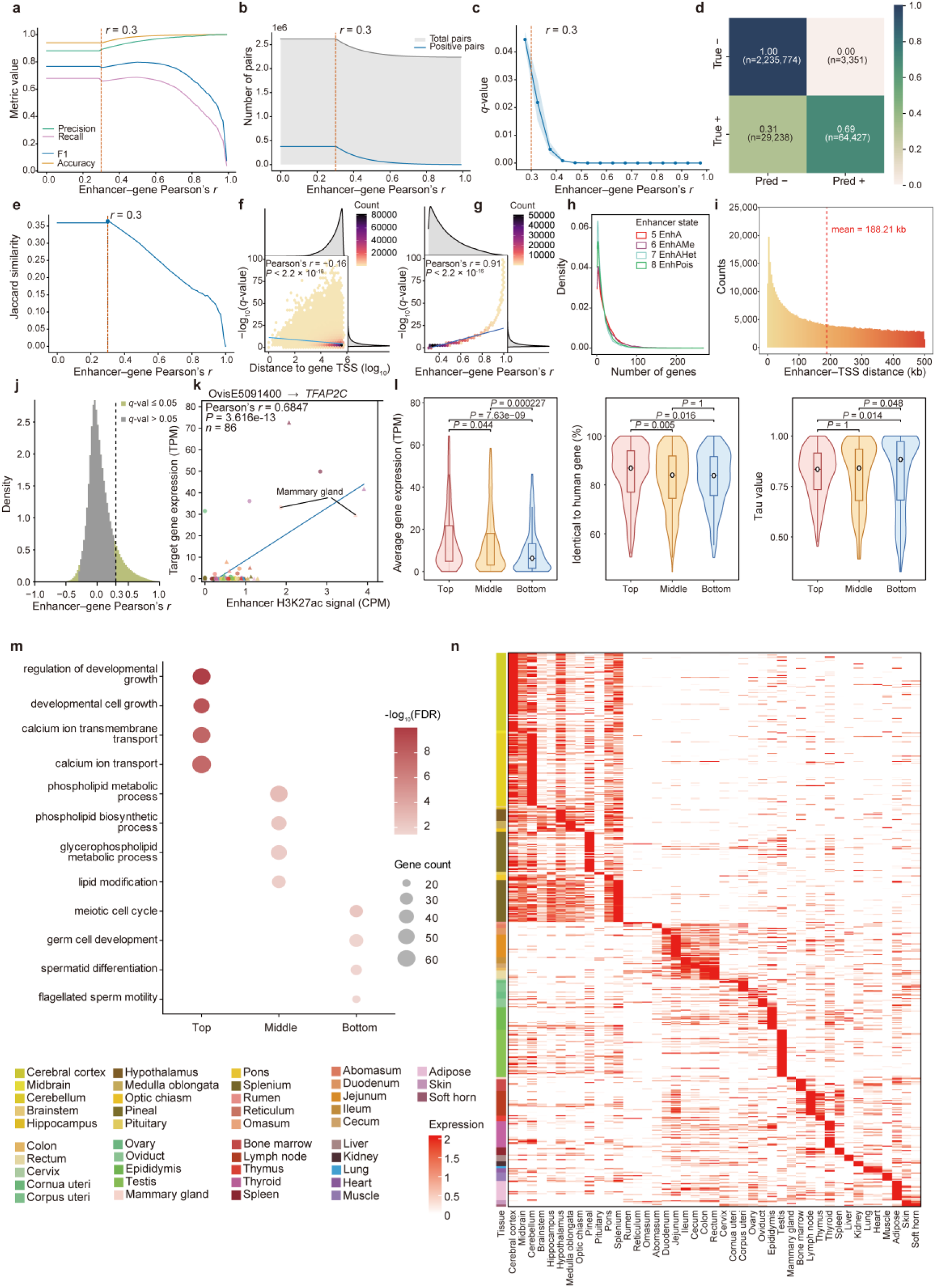
**Evaluation and characterization of enhancer–target gene pairs**. a, Classification metrics for H3K27ac-derived enhancer–target gene pairs relative to ATAC-derived links across enhancer–gene Pearson’s r thresholds. b, Number of retained enhancer– target gene pairs across Pearson’s r thresholds. c, Distribution of q-values across enhancer–gene Pearson’s r values. d, Confusion matrix for enhancer–target gene link classification. e, Jaccard similarity between H3K27ac-derived and ATAC-derived enhancer–target gene pairs across Pearson’s *r* thresholds. f, Relationship between enhancer–TSS distance and enhancer–target gene association significance. g, Relationship between enhancer–gene Pearson’s r and association significance. h, Density distribution of the number of linked genes per enhancer, stratified by enhancer state. i, Distribution of enhancer–TSS distances. j, Density distribution of enhancer–gene Pearson’s r for significant and non-significant enhancer–target gene pairs. k, Representative enhancer–target gene pair showing the correlation between enhancer H3K27ac signal and target-gene expression across tissues. l, Comparison of genes grouped by the number of linked enhancers, showing gene expression, human orthologue identity and tissue-specificity index Tau. m, Gene Ontology enrichment analysis of enhancer-linked gene groups. n, Heatmap showing tissue-specific expression patterns of enhancer-linked target genes across tissues.

**Extended Data Fig. 4.**
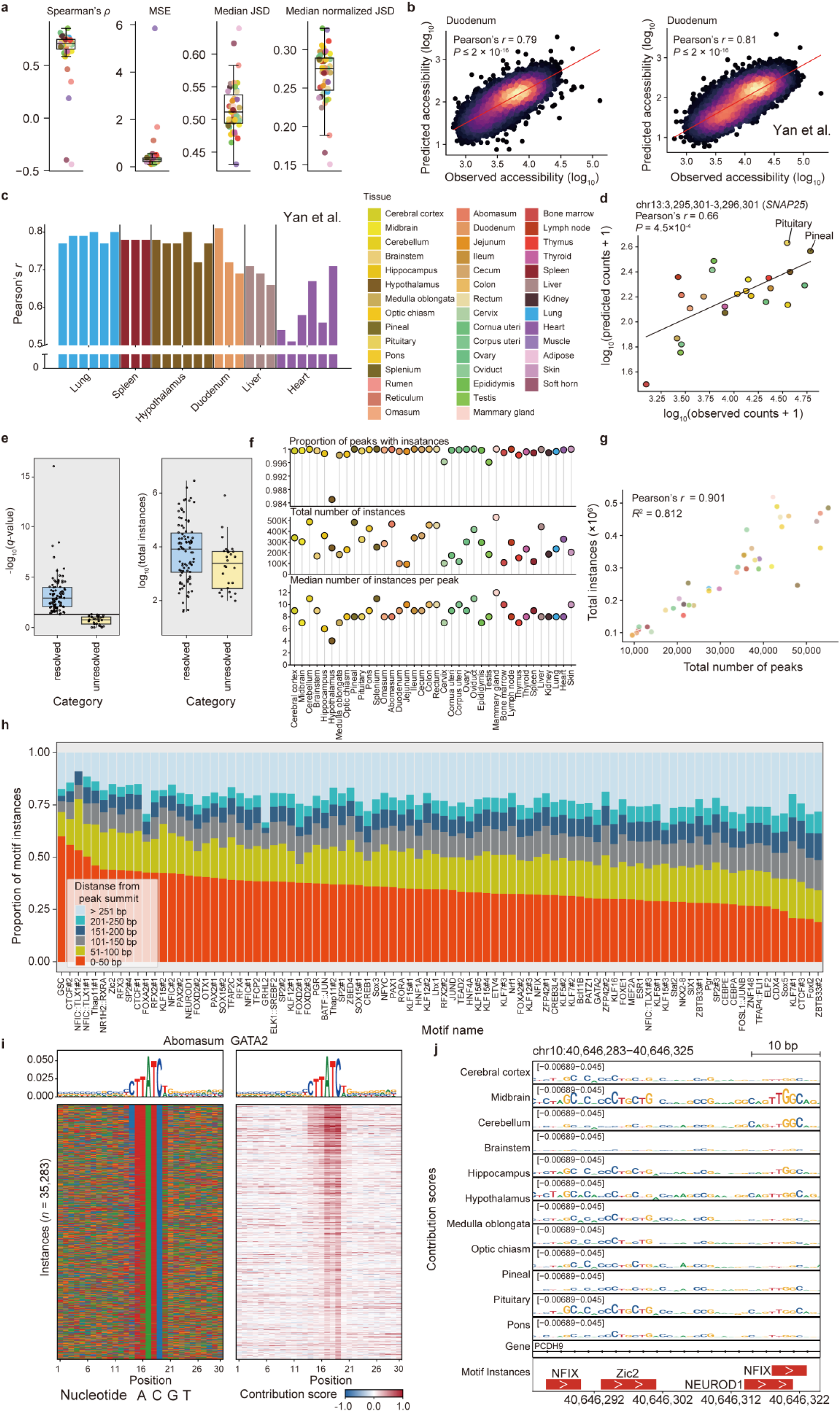
Performance evaluation of sheep multi-tissue ChromBPNet models and overview of motif-instance features. a, ChromBPNet model performance across held-out tissues, including Spearman’s ρ, mean squared error (MSE), median Jensen–Shannon divergence (JSD) and median normalized JSD. b, Observed and predicted log_10_-transformed ATAC-seq accessibility in duodenum, shown for the matched dataset generated in this study and the external dataset from Yan et al. c, Pearson’s r between observed and predicted accessibility in tissues overlapping with Yan et al. d, Observed and predicted log_10_(counts + 1) across tissues at a representative region near *SNAP25* on chr13. e, Resolved and unresolved motif categories summarized by motif-matching significance and total instance number. f, Motif-instance annotation across tissues, including peak coverage, total instance number and median number of instances per peak. g, Relationship between motif-instance number and ATAC-seq peak number across tissues. h, Motif-instance distribution by binned distance to the nearest peak summit. i, GATA2 motif instances in abomasum, shown with motif logo, nucleotide composition and contribution-score heatmaps. j, Contribution-score tracks and motif instances across neural tissues at a representative region near *PCDH9* on chr10.

**Extended Data Fig. 5.**
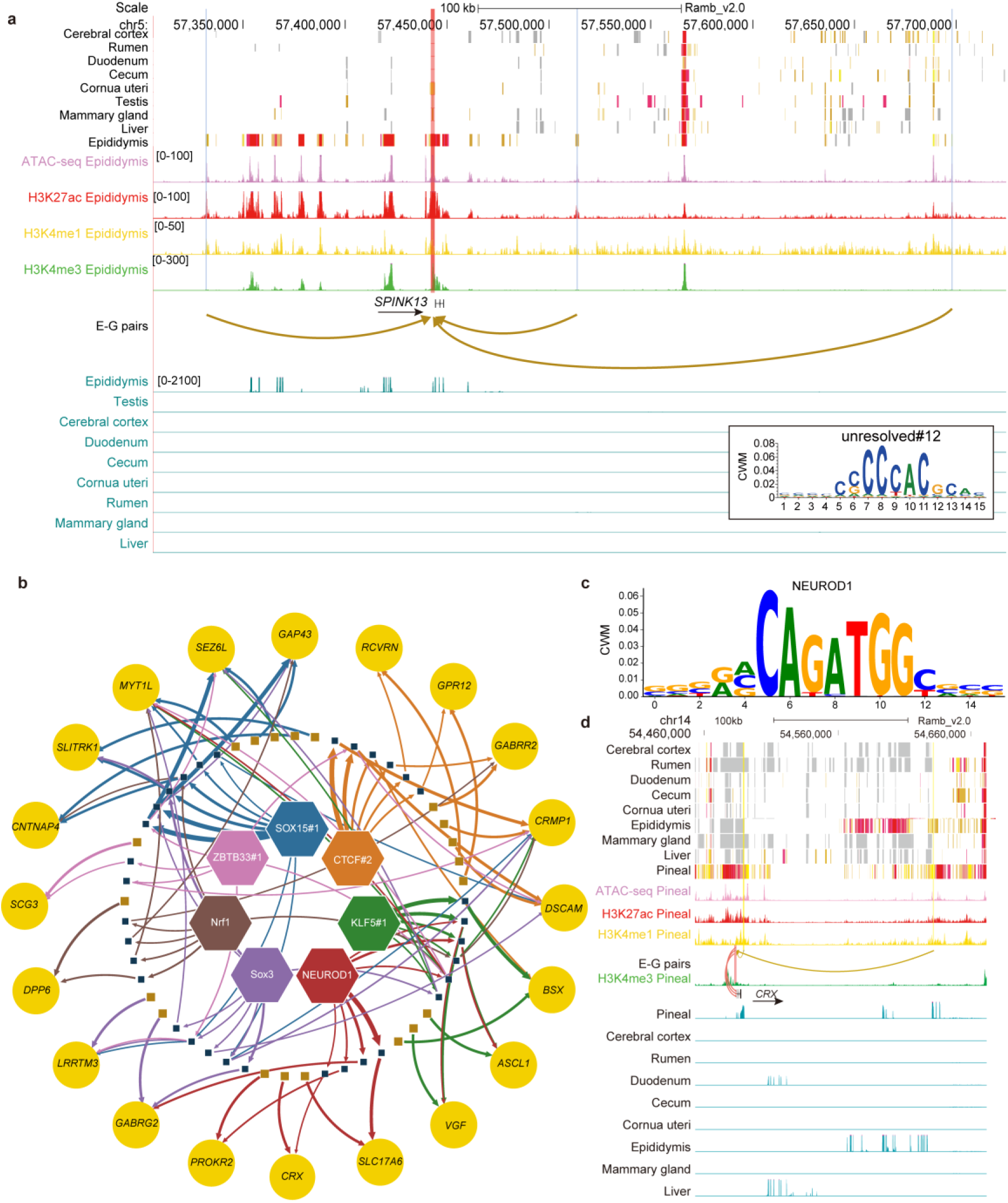
Additional tissue-associated ChromBPNet regulatory networks and representative loci. a, Epididymis-associated regulatory annotation of the *SPINK13* locus on chr5:57.35–57.70 Mb, showing chromatin states, ATAC-seq and histone-mark signals, enhancer–gene links, tissue-resolved RNA-seq expression and the unresolved motif 12 contribution weight matrix. b, Nervous-system-associated regulatory subnetwork connecting selected predictive motifs and candidate regulatory elements to target genes. c, NEUROD1 contribution weight matrix. d, Regulatory annotation of the *CRX* locus on chr14:54.46–54.66 Mb, showing chromatin states, pineal epigenomic signals, enhancer–gene pairs and tissue-resolved RNA-seq expression.

**Extended Data Fig. 6.**
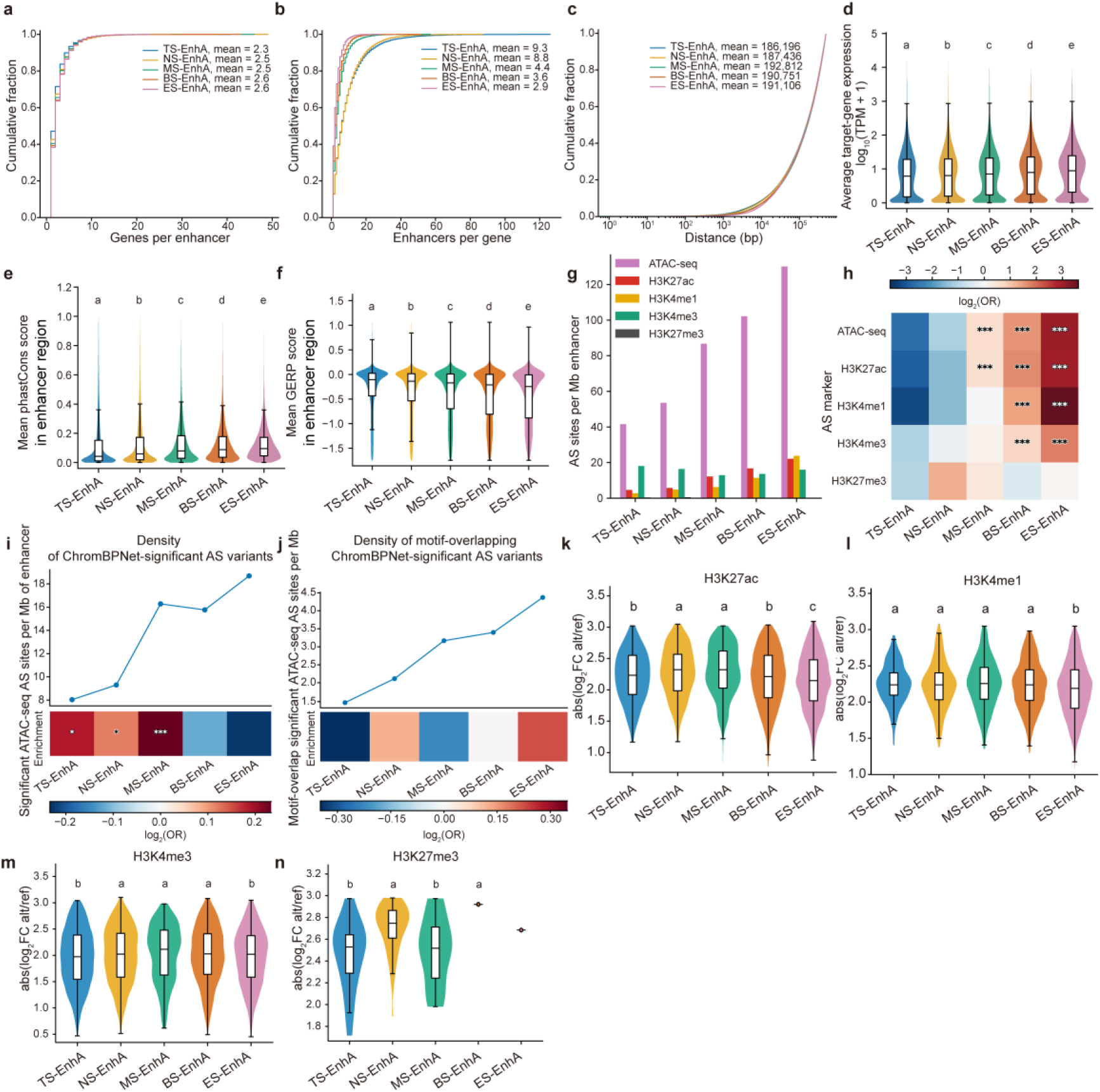
Target-gene connectivity, conservation and genetic-variation features of active enhancers with different tissue-sharing breadths. a–c, Cumulative distributions of target-gene connectivity features across active-enhancer groups, including the number of linked genes per enhancer, the number of enhancers linked to each gene and enhancer–target-gene distance. d, Average expression of target genes linked to active-enhancer groups. e, Mean phastCons conservation scores across active-enhancer groups. f, Mean GERP conservation scores across active-enhancer groups. g, Density of allele-specific sites from different assays across active-enhancer groups. h, Relative enrichment of allele-specific sites across active-enhancer groups and assays. i, Density and relative enrichment of ChromBPNet- significant ATAC-seq allele-specific variants across active-enhancer groups. j, Density and relative enrichment of motif-overlapping ChromBPNet-significant ATAC-seq allele-specific variants across active-enhancer groups. k–n, Effect-size distributions of allele-specific sites from H3K27ac, H3K4me1, H3K4me3 and H3K27me3 across active-enhancer groups.

**Extended Data Fig. 7.**
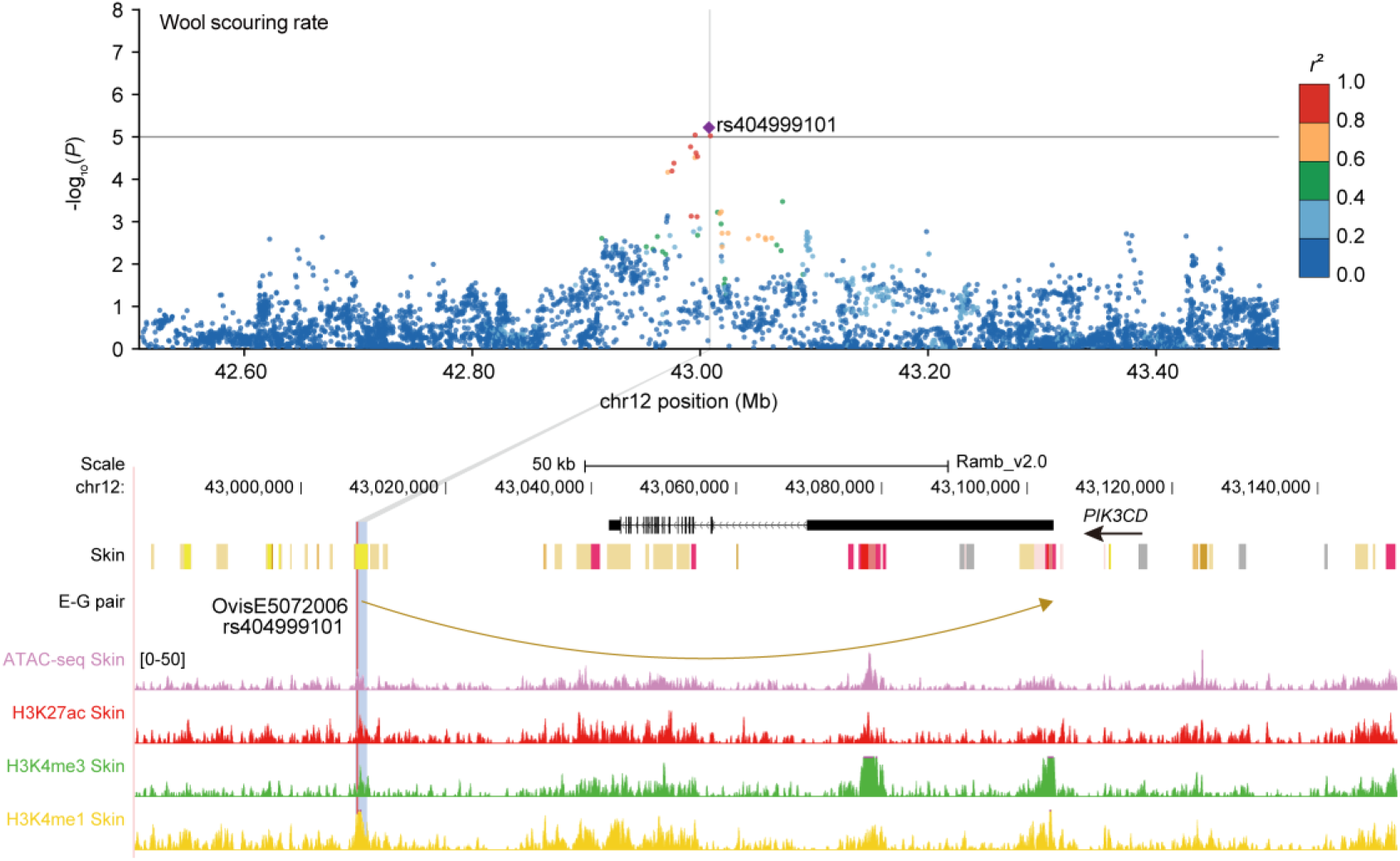
Regulatory annotation of additional candidate regulatory loci for sheep complex-trait GWAS signals. Regulatory annotation of the wool-scouring-rate-associated variant rs404999101 at the *PIK3CD* locus. The upper panel shows regional GWAS association signals, with rs404999101 highlighted; the y axis denotes −log10(*P*), and point color indicates linkage disequilibrium with the lead SNP (*r*²). Genome-browser tracks show *PIK3CD* gene annotation, skin chromatin states, enhancer–gene pairs, the SNP-containing enhancer element OvisE5072006 and skin epigenomic signals, including ATAC-seq, H3K27ac, H3K4me3 and H3K4me1. GWAS, genome-wide association study; E-G pair, enhancer–gene pair.

**Extended Data Fig. 8.**
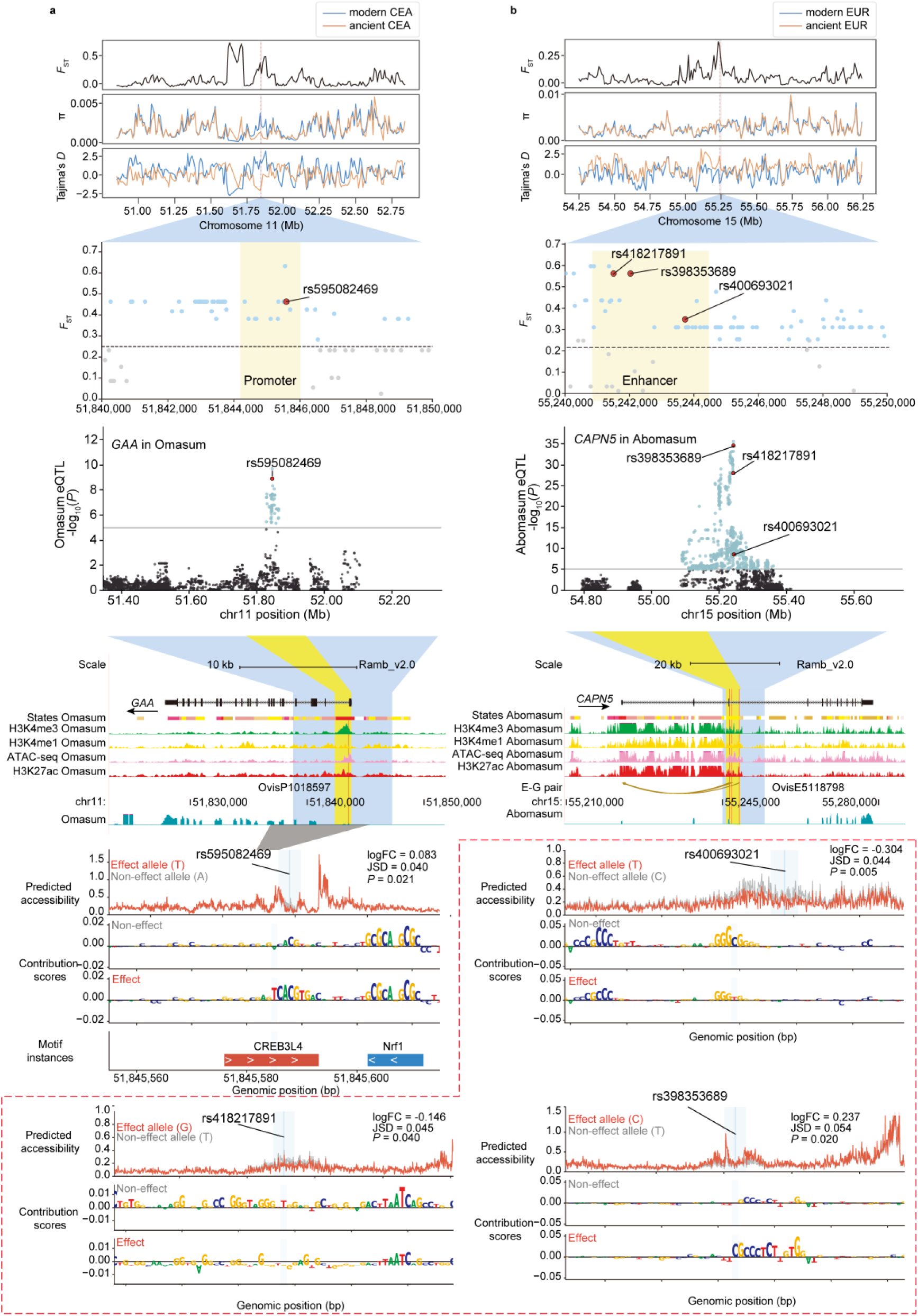
| Supplementary regulatory annotation of candidate selected variants. a, Regulatory annotation and ChromBPNet allele-effect prediction for the candidate selected variant rs595082469 at the *GAA* locus. The upper tracks show *F*_ST_, nucleotide diversity (π) and Tajima’s *D* signals across the surrounding chr11 region. The local *F*_ST_ panel highlights the promoter region containing rs595082469. The eQTL panel shows the omasum Eqtl association signal for *GAA*. Genome-browser tracks show *GAA* gene annotation, omasum chromatin states, promoter annotation, epigenomic signals and the candidate variant. The lower panels show ChromBPNet-predicted accessibility, base-resolution contribution scores and motif instances for the two alleles. b, Regulatory annotation and ChromBPNet allele-effect prediction for candidate selected variants near the *CAPN5* locus. The upper tracks show *F*_ST_, nucleotide diversity (π) and Tajima’s *D* signals across the surrounding chr15 region. The local *F*_ST_ panel highlights the enhancer region containing rs418217891, rs398353689 and rs400693021. The eQTL panel shows the abomasum eQTL association signal for *CAPN5*. Genome-browser tracks show *CAPN5* gene annotation, abomasum chromatin states, enhancer– gene pairs, epigenomic signals, RNA-seq signals and the candidate variants. The lower panels show ChromBPNet-predicted accessibility and base-resolution contribution scores for the two alleles at each variant.

## Notes

### Competing Interest Statement

The authors have declared no competing interest.

