## Supplementary note for "A multi-tissue epigenomic atlas links the sheep non-coding genome to domestication and complex traits"

The supplementary information contains:

**Supplementary Note**

**Supplementary Figures 1–18**

**Supplementary References**

**Supplementary Tables 1–26** (in a separate Excel file)

### Supplementary Note

#### RNA-seq, ATAC-seq and CUT&Tag library preparation and sequencing

Frozen tissues were used for RNA-seq, ATAC-seq and CUT&Tag library construction. CUT&Tag was performed for four histone modifications: H3K4me3, H3K27ac, H3K4me1 and H3K27me3.

For RNA-seq, total RNA was extracted using TRIzol reagent and RNA integrity was assessed. Samples with an RNA integrity number (RIN) of at least 6 were retained. Ribosomal RNA was depleted before sequencing, and strand-specific RNA-seq libraries were prepared using the dUTP method according to the Illumina protocol. Libraries were sequenced on an Illumina HiSeq X Ten platform in paired-end 150-bp mode.

For ATAC-seq, nuclei were isolated from frozen tissues using the Boyou nuclei isolation kit (52201-10). Briefly, approximately 5 mg of tissue was added to 0.5 ml lysis buffer, homogenized with a pestle until no visible tissue fragments remained, incubated on ice for 2–10 min, filtered to remove debris and resuspended in suspension buffer to obtain approximately 50,000 nuclei. ATAC-seq libraries were prepared using the Hyperactive ATAC-Seq Library Prep Kit for Illumina (TD711). Nuclei were incubated with the transposition reaction mixture at 37 °C for 30 min. DNA was recovered using ATAC DNA extraction beads, purified using ATAC DNA purification beads, amplified and subjected to two-step size selection. Qualified libraries were sequenced on an Illumina HiSeq X Ten platform in paired-end 150-bp mode.

For CUT&Tag, nuclei were isolated from frozen tissues using the same procedure as described for ATAC-seq, and approximately nuclei were used per reaction. Libraries were prepared using the Hyperactive Universal CUT&Tag Assay Kit for Illumina Pro (Vazyme, TD904) according to the manufacturer's instructions. Nuclei were immobilized on concanavalin A-coated magnetic beads and incubated overnight at 4 °C in 50 µl antibody buffer with one of the following affinity-purified rabbit polyclonal antibodies from Diagenode: H3K4me3 (0.4 µl, approximately 0.52 µg; cat. no. C15410003; lot A8034D), H3K4me1 (0.66 µl, approximately 0.99 µg; cat. no. C15410194; lot A1862D), H3K27ac (0.4 µl, approximately 1.12 µg; cat. no. C15410196; lot A1723-0041D) or H3K27me3 (0.9 µl, approximately 0.99 µg; cat. no. C15410195; lot A0824D). The nuclei–bead complexes were subsequently incubated with anti-rabbit secondary antibody in 50 µl Dig-wash buffer for 1 h at room temperature. After washing, the complexes were incubated with 2 µl pA/G-Tnp Pro diluted in 98 µl Dig-300 buffer for 1 h at room temperature. Tagmentation was performed in a 50-µl reaction containing 40 µl Dig-300 buffer and 10 µl 5× TTBL at 37 °C for 1 h. DNA fragments were purified using VAHTS DNA Clean Beads, quantified and used for library amplification. Qualified libraries were sequenced on an Illumina HiSeq X Ten platform in paired-end 150-bp mode. All four antibodies had been validated by the manufacturer for CUT&Tag using 50,000 K562 cells and showed modification-specific genomic enrichment. Antibody performance in sheep tissues was further assessed from the concordance between biological replicates and the expected genomic distribution of the corresponding histone modifications.

#### Overview of functional genomic data in the SheepEpimap project

The sequencing experiments yielded 14.88 billion raw reads, of which 13.99 billion (94.02%) were uniquely mapped and retained after alignment and filtering **(Supplementary Table 1)**. This effort represents a 3.78-fold increase in tissue coverage and a 4.62-fold increase in dataset volume compared to previous sheep epigenomic efforts[1]. The genome-wide average Pearson correlation between observed and imputed signals was 0.68, comparable to that reported for human[2] and cattle[3] datasets **(Supplementary Figs. 1 and 2; Supplementary Table 2)**. Gene expression levels showed weak positive correlations with active chromatin features and negative correlations with H3K27me3, supporting the overall data quality **(Fig. 1b; Supplementary Fig. 3)**. Compared with the previous version of the atlas[1], Protein-coding genes exhibited the most typical promoter-centered regulatory architecture, characterized by pronounced chromatin accessibility and enrichment of H3K4me3 and H3K27ac near transcription start sites (TSSs), whereas H3K4me1 signals were relatively weaker and more broadly distributed **(Supplementary Fig. 4a–f)**.

### Sample clustering

Sample clustering and correlation analyses were performed using deepTools (v3.5.0)[4]. For each sample, bigWig signal tracks were generated using bamCompare and standardized by Z-score transformation using scipy.stats.zscore in SciPy (v1.8.0)[5]. Standardized signal matrices were summarized with multiBigwigSummary, sample correlations were calculated with plotCorrelation, and principal component analysis was performed with plotPCA. Z-score-normalized signals across protein-coding genes, lncRNAs, pseudogenes, miRNAs, snRNAs and snoRNAs were calculated using deepTools computeMatrix scale-regions with parameters --afterRegionStartLength 2500 and --beforeRegionStartLength 2500.

### Evolutionary conservation analysis

Enrichment of chromatin states in genomic features, including exons, transcription start sites (TSSs), CpG islands, conserved sequence regions and VISTA enhancers, was calculated as (C/A)/(B/D), where A is the number of bases covered by a chromatin state, B is the number of bases covered by a genomic feature, C is the number of bases overlapping both the chromatin state and the genomic feature, and D is the total number of bases in the genome. phastCons and phyloP conservation scores for 100 vertebrates were downloaded from UCSC for hg38, and human and sheep GERP scores were downloaded from Ensembl. Conservation scores were converted from hg38 to sheep genome coordinates using UCSC liftOver. Experimentally validated human VISTA enhancers were downloaded from the VISTA Enhancer Browser, including enhancers annotated for all tissues, forebrain, midbrain, hindbrain, neural tube, limb and heart. Enrichment significance between the 11 chromatin states and human enhancers was assessed using Fisher's exact test.

### Enhancer-gene pair characterization

Based on high-confidence enhancer–gene links, the number of target genes connected to each enhancer, enhancer-to-TSS distance distributions and linking patterns across enhancer states were analyzed. Target genes were grouped according to the number of linked E5 enhancers: Top-group genes were linked to ≥56 enhancers, Middle-group genes were linked to 13–18 enhancers and Bottom-group genes were linked to ≤2 enhancers. Mean expression, human orthologue consistency and Tau index were compared among the three groups using the Wilcoxon rank-sum test. GO Biological Process enrichment analysis was performed separately for the three gene groups, and clustered heatmaps were used to visualize cross-tissue expression patterns of high-confidence enhancer target genes.

### Super-enhancer identification and functional enrichment

The four enhancer chromatin states, EnhA, EnhAMe, EnhAHet and EnhPois, were merged across tissues into a GFF file. Super-enhancers were identified using ROSE (v1.3.1)[6] based on H3K27ac signal for each sample in each tissue. Enhancer peaks located within 12.5 kb of one another were stitched into enhancer clusters. Read coverage was calculated for each peak, and signals were summed across all enhancers within each cluster. Putative super-enhancers were identified from the ROSE rank-order curve, with enhancers above the inflection point defined as super-enhancers. Tissue-specific super-enhancers were identified using the same approach as for tissue-specific regulatory elements.

Enrichment of super-enhancer states in genomic features was calculated as (C/A)/(B/D), where A is the length of super-enhancer states, B is the length of genomic features, C is the length of overlap between super-enhancer states and genomic features, and D is the total genomic length. Overlaps were determined using BEDTools (v2.29.2). Super-enhancer clustering was performed using ComplexHeatmap (v2.9.3)[7]. GO enrichment analysis of genes associated with super-enhancers was performed using clusterProfiler (v4.4.1). Motifs significantly enriched in super-enhancers were identified using HOMER (v4.11) with FDR < 0.05.

### Functional characteristics of tissue-specific chromatin states

GO enrichment analysis of tissue-specific EnhA-associated genes showed that digestive tissues were enriched for digestive, immune and epithelial-related processes, whereas reproductive tissues were enriched for embryonic development, sex differentiation and reproduction-related processes. Immune and muscle tissues were enriched for terms related to immune regulation and muscle function, respectively **(Fig. 2g; Supplementary Table 8)**. Motif enrichment analysis further revealed lineage-associated transcription factor signatures in tissue-specific EnhA elements from different tissue systems. EnhA elements in nervous system tissues were enriched for SOX3 and SOX5 motifs; those in muscle tissues were enriched for MEF2A and FHL1 motifs; those in digestive and epithelial-related tissues were enriched for KLF1, KLF5, and HNF4A motifs; those in reproductive tissues were enriched for GATA2 and PGR motifs; and those in immune tissues were enriched for SPI1 and RUNX3 motifs **(Fig. 2h; Supplementary Table 9)**.

### Construction of a sheep gene regulatory network

In the nervous system-associated subnetwork, candidate TF motifs, including NEUROD1 and members of the SOX family, formed putative regulatory relationships with multiple neural tissue-related genes[8,9]. At the *CRX* locus, NEUROD1 motif-associated regulatory connections were consistent with local chromatin interactions and tissue-specific epigenomic signals, further supporting the ability of this network to characterize regulatory relationships in neural tissues **(Extended Data Fig. 5b-d)**[10].

#### Construction of an atlas of allele-specific regulatory variants in sheep

Although a subset of AS SNPs was shared across multiple tissues, most AS loci exhibited strong tissue specificity Across omics layers, the proportion of AS loci detected in only one tissue ranged from 69.0% for RNA-seq to 99.4% for H3K4me1, indicating that allele-specific regulatory effects are jointly shaped by tissue context and chromatin state **(Supplementary Fig. 14f)**. Tissue-level sharing networks further revealed extensive but uneven sharing of AS signals among tissues, with strong connections between some tissues, whereas heart, testis, pons, and other tissues showed relatively independent allelic regulatory patterns **(Supplementary Fig. 14d,g)**. These results suggest that AS regulatory variants exhibit both cross-tissue sharing and pronounced tissue-specific regulatory features. These enrichment signals were observed across multiple traits, including body-measurement traits, horn length, and wool-related traits, suggesting that AS regulatory elements can provide an important basis for prioritizing candidate functional variants underlying complex traits **(Fig. 5n; Supplementary Fig. 15)**.

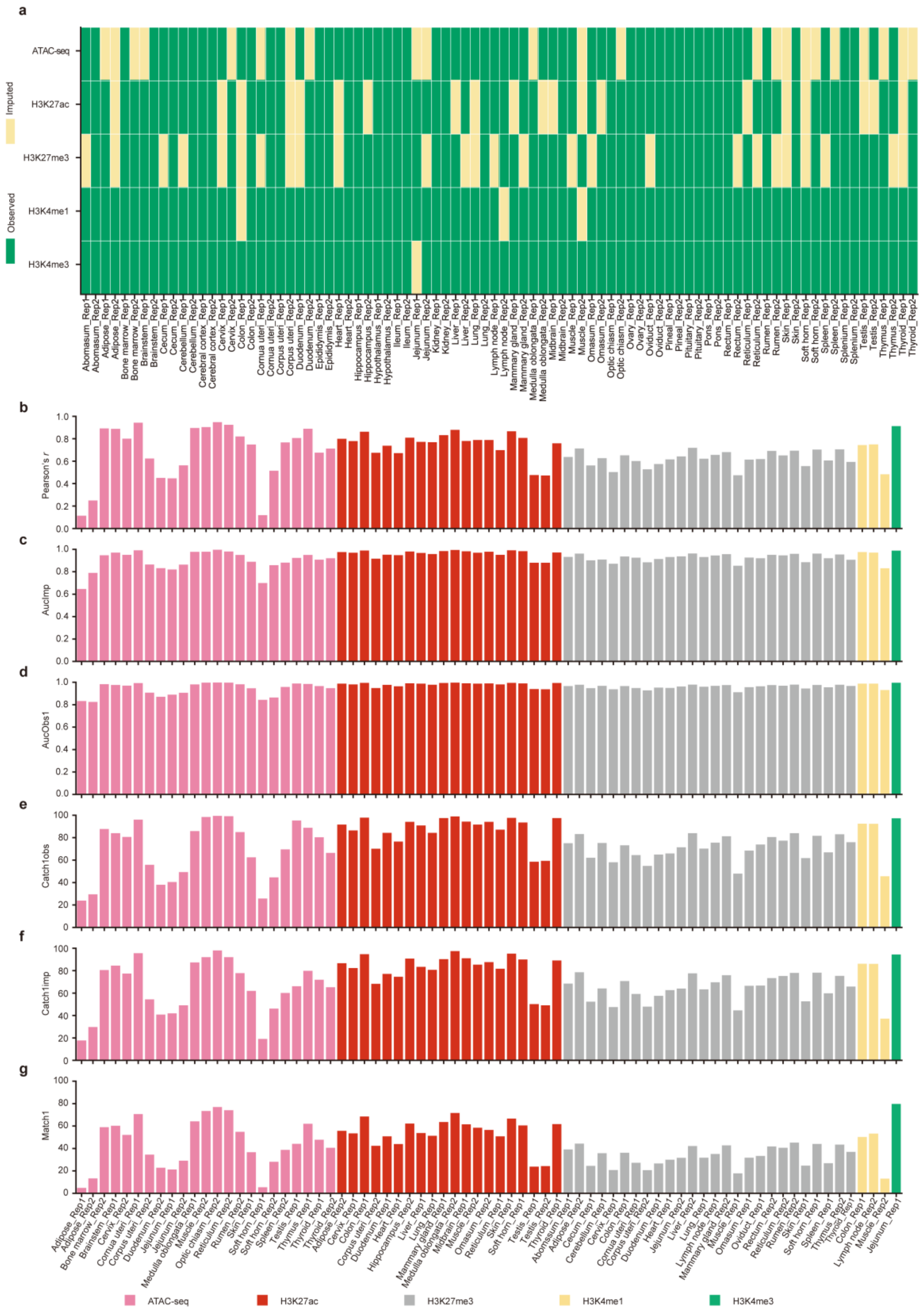


**Supplementary Fig. 1 Validation of epigenome imputation.** a, Overview of observed and imputed epigenomic datasets used for imputation validation across tissues, biological replicates and assays, including ATAC-seq, H3K27ac, H3K4me1, H3K27me3 and H3K4me3. **b–g,** Evaluation of imputation performance across held-out assay–tissue replicate combinations using six quality metrics. **b,** Pearson's *r* between observed and imputed signal tracks. **c,** AucImp, defined as the area under the receiver operating characteristic (ROC) curve for predicting the top 1% imputed signal using the full range of observed signal. **d,** AucObs1, defined as the area under the ROC curve for predicting the top 1% observed signal using the full range of imputed signal. **e,** Catch1obs, defined as the percentage of observed top 1% genomic bins recovered within the imputed top 5% bins. **f,** Catch1imp, defined as the percentage of imputed top 1% genomic bins recovered within the observed top 5% bins. **g,** Match1, defined as the percentage overlap between the observed top 1% and imputed top 1% genomic bins. Bars are colored by assay.

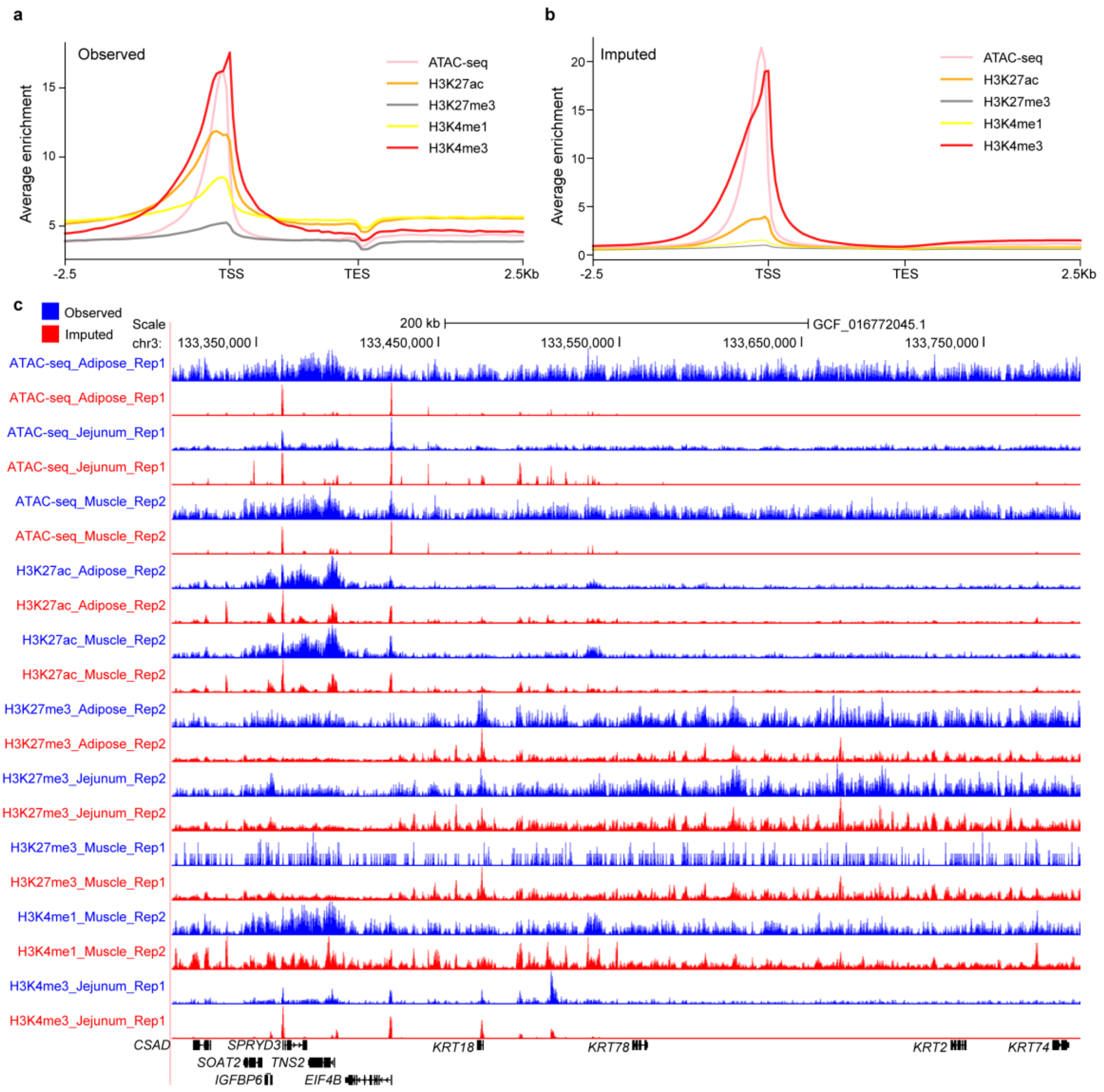


**Supplementary Fig. 2 Quality control of imputed epigenomic profiles for missing and low-quality datasets. a,b,** Average enrichment profiles of observed (**a**) and imputed (**b**) epigenomic signals across gene bodies from transcription start sites (TSS) to transcription end sites (TES), including 2.5-kb flanking regions upstream of TSSs and downstream of TESs. Profiles are shown for ATAC-seq and CUT&Tag histone-mark datasets, including H3K27ac, H3K27me3, H3K4me1 and H3K4me3. **c,** Genome browser snapshot comparing observed and imputed signal tracks across a representative genomic region on chr3:133.35–133.80 Mb. Observed and imputed signals are shown in blue and red, respectively.

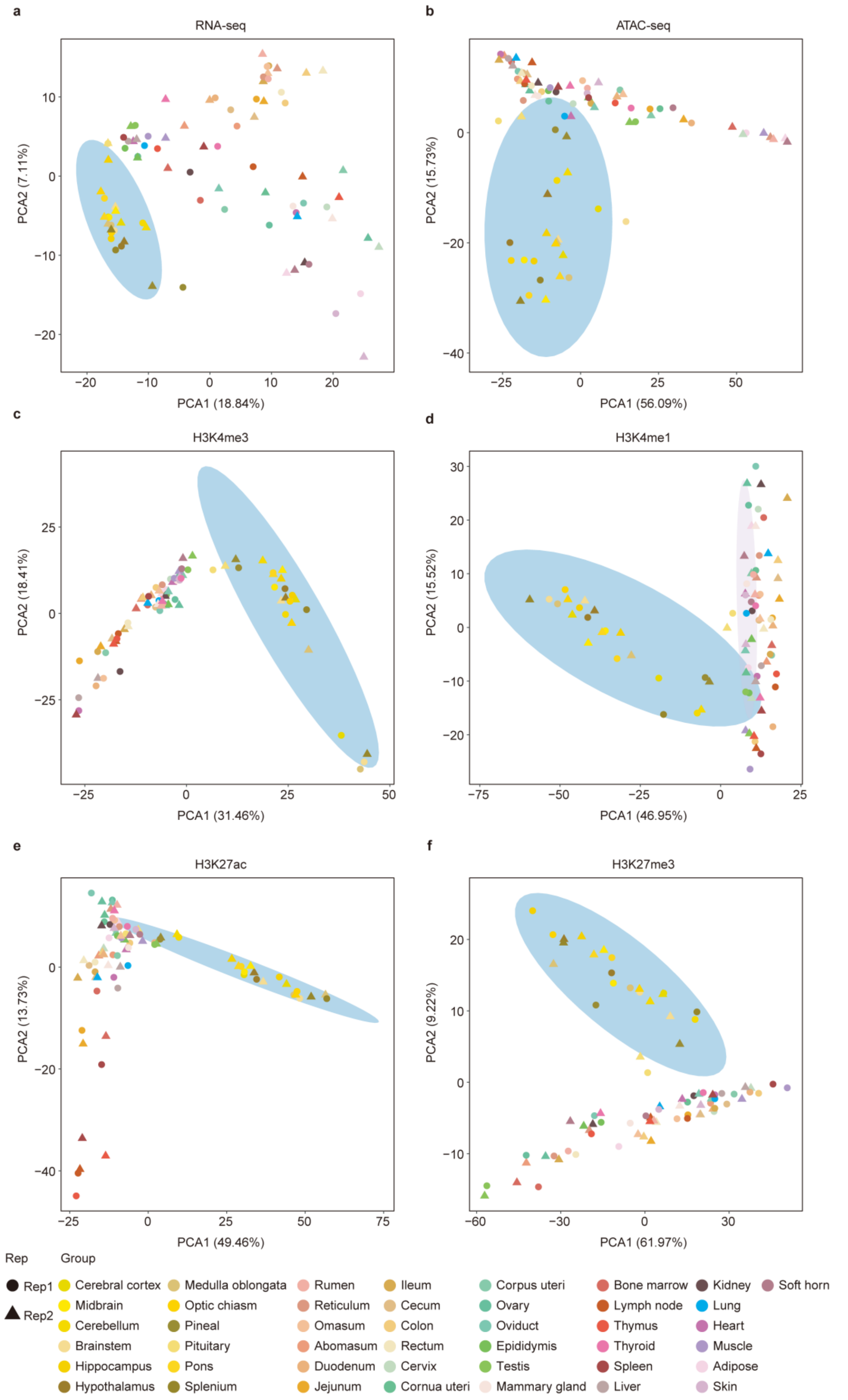


**Supplementary Fig. 3** Principal component analysis of epigenomic and transcriptomic profiles across 43 sheep tissues. a-f, Principal component analysis (PCA) of 43 sheep tissues based on five epigenomic marks and RNA-seq data. Colours and legend symbols are consistent across all six panels

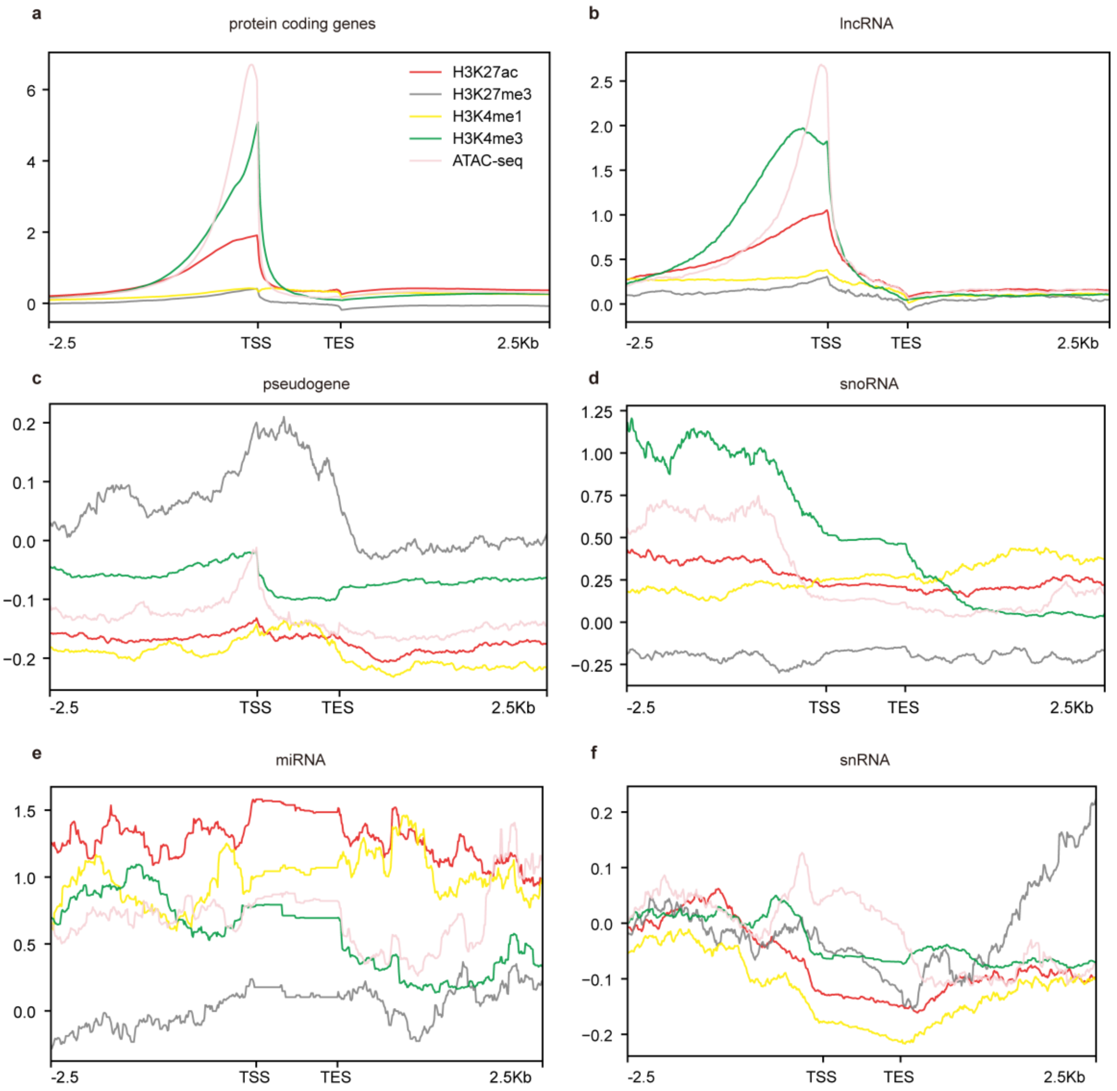


**Supplementary Fig. 4 Distribution of normalized epigenomic signals across gene classes.** a–f, Average normalized H3K27ac, H3K27me3, H3K4me1, H3K4me3 and ATAC-seq signals across protein-coding genes (a), long non-coding RNAs (b), pseudogenes (c), small nucleolar RNAs (d), microRNAs (e) and small nuclear RNAs (f), including 2.5-kb regions upstream of the transcription start site and downstream of the transcription end site. ATAC-seq, assay for transposase-accessible chromatin using sequencing; lncRNA, long non-coding RNA; miRNA, microRNA; snoRNA, small nucleolar RNA; snRNA, small nuclear RNA; TSS, transcription start site; TES, transcription end site.

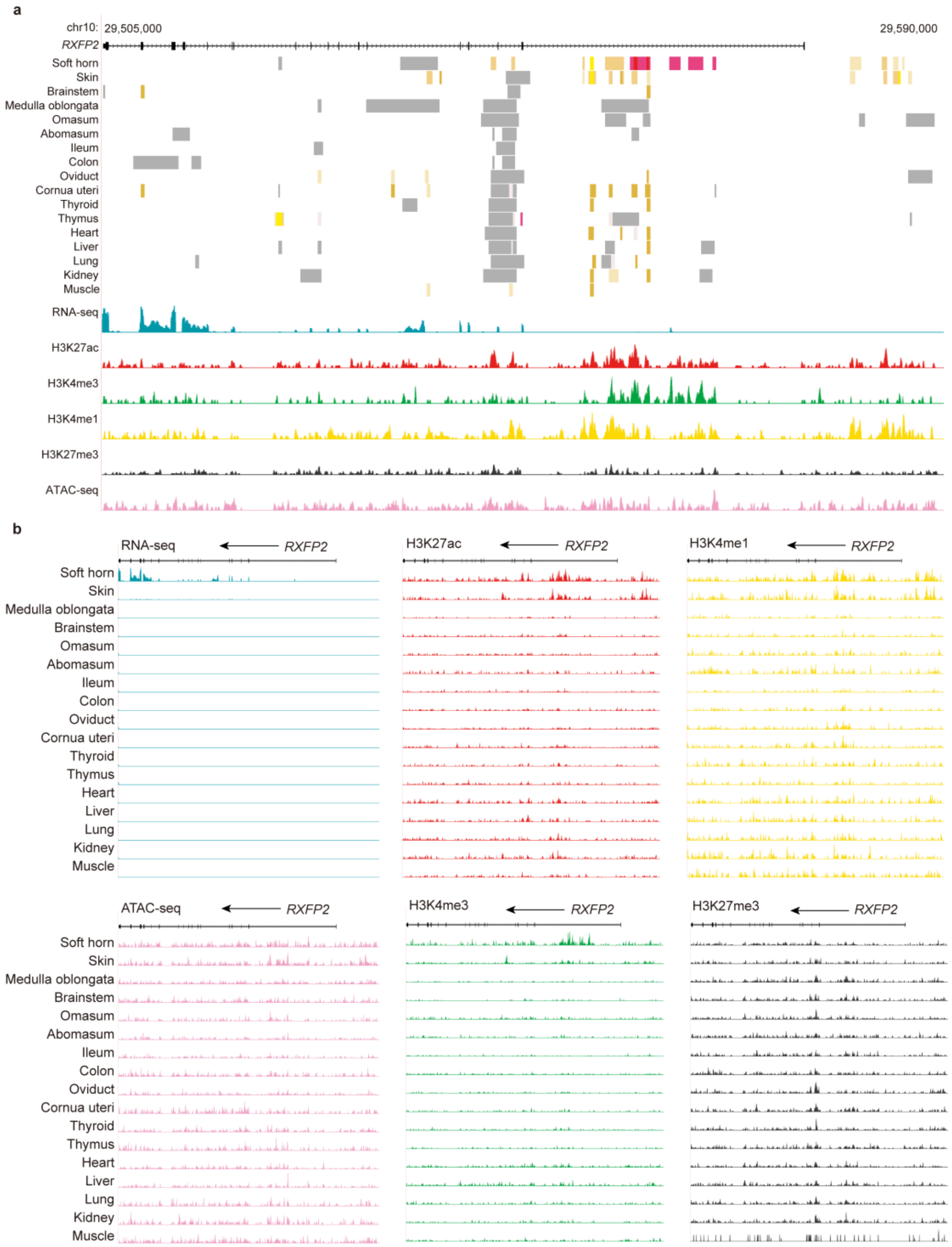


**Supplementary Fig. 5 Chromatin-state and multi-omics profiles at the RXFP2 locus.** a, Tissue-resolved chromatin-state annotations and averaged multi-omics signals across chr10:29,493,543–29,583,065. b, RNA-seq, H3K27ac, H3K4me1, ATAC-seq, H3K4me3 and H3K27me3 signal tracks across 17 representative tissues. Vertical scales were set to 0–50 for RNA-seq, H3K27ac and H3K4me3, 0–30 for H3K4me1 and ATAC-seq, and 0–80 for H3K27me3.

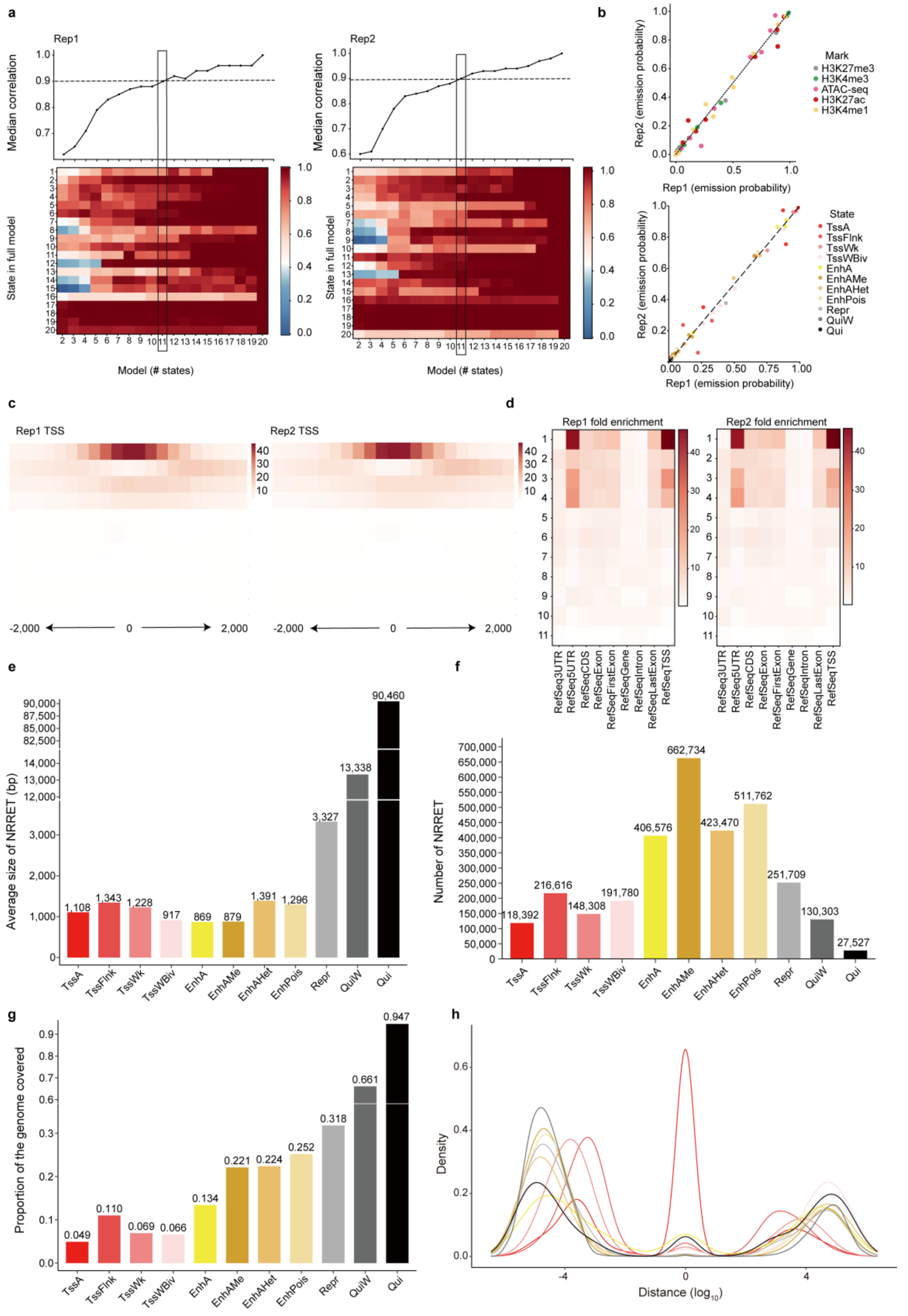


**Supplementary Fig. 6 ChromHMM model selection and chromatin-state characterization.** a, Comparison of ChromHMM models with different numbers of states in Rep1 and Rep2, with the 11-state model highlighted. b, Concordance of emission probabilities between Rep1 and Rep2, coloured by epigenomic mark or chromatin state. c, Enrichment profiles of chromatin states around transcription start sites in Rep1 and Rep2. d, Enrichment of chromatin states across RefSeq genomic features in Rep1 and Rep2. e, Average size of non-redundant regulatory elements across chromatin states. f, Number of non-redundant regulatory elements across chromatin states. g, Genome coverage of non-redundant regulatory elements across chromatin states. h, Distance distribution from non-redundant regulatory elements to transcription start sites across chromatin states.

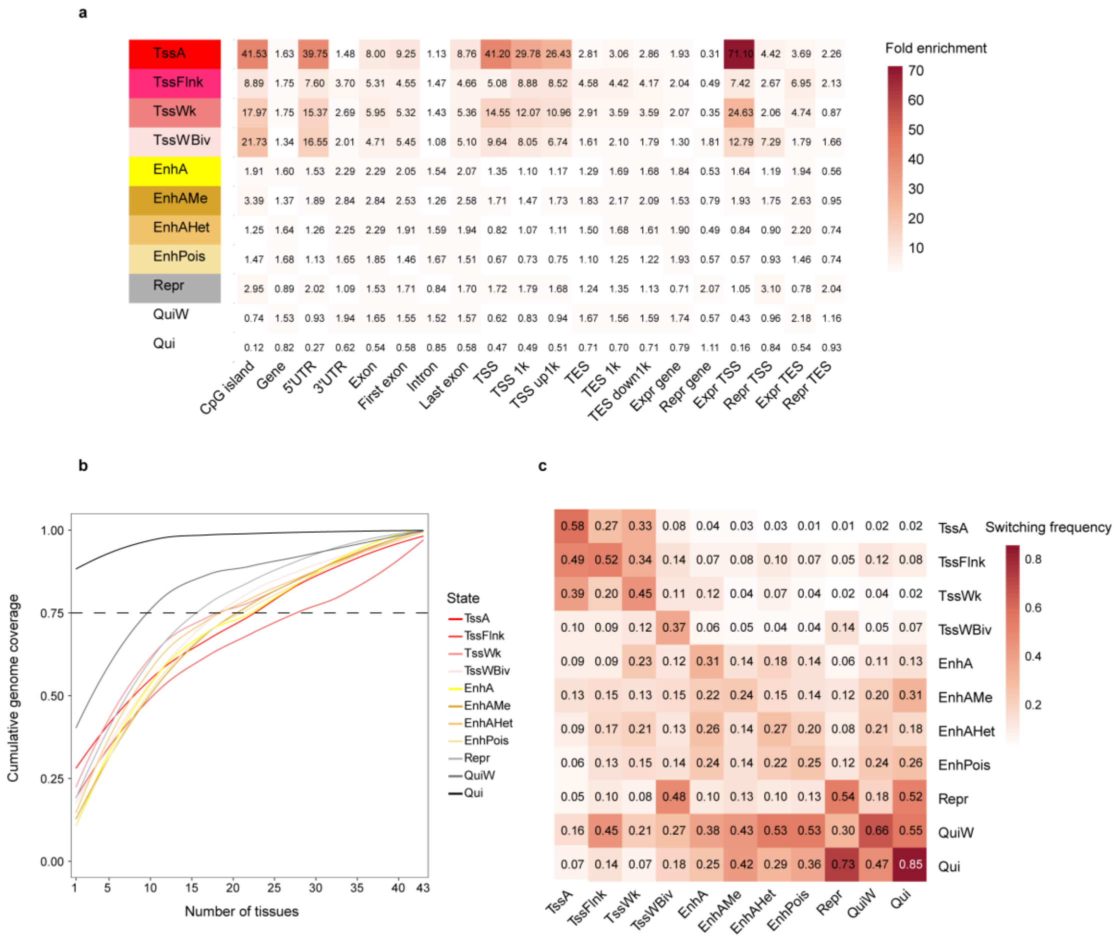


**Supplementary Fig. 7 Genomic enrichment, tissue sharing and switching patterns of chromatin states.** a, Fold enrichment of chromatin states across selected genomic, transcript and expression-based annotations. b, Tissue-sharing patterns of chromatin states, shown as cumulative genome coverage across increasing numbers of tissues. The dashed line indicates a cumulative genome coverage of 0.75. c, Pairwise chromatin-state switching frequency across tissues. Rows and columns represent chromatin states, and colour intensity indicates switching frequency.

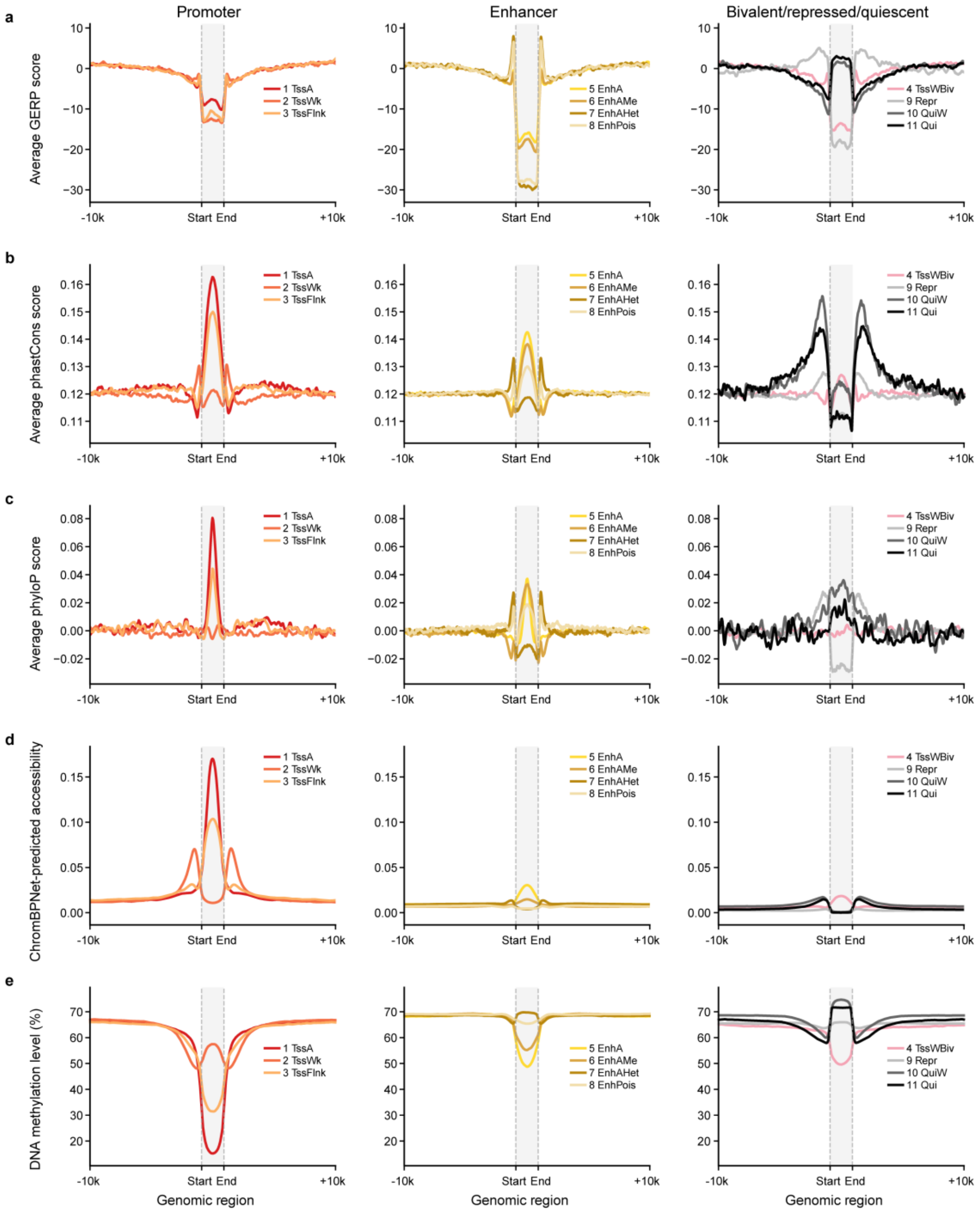


**Supplementary Fig. 8 Conservation, accessibility and DNA methylation profiles around chromatin states.** a–c, Average GERP, phastCons and phyloP conservation scores across chromatin-state regions and their flanking sequences. d, ChromBPNet-predicted accessibility profiles across chromatin-state regions and flanking sequences. e, DNA methylation profiles across chromatin-state regions and flanking sequences. Chromatin states are grouped into promoter, enhancer, and bivalent/repressed/quiescent categories. GERP, Genomic Evolutionary Rate Profiling.

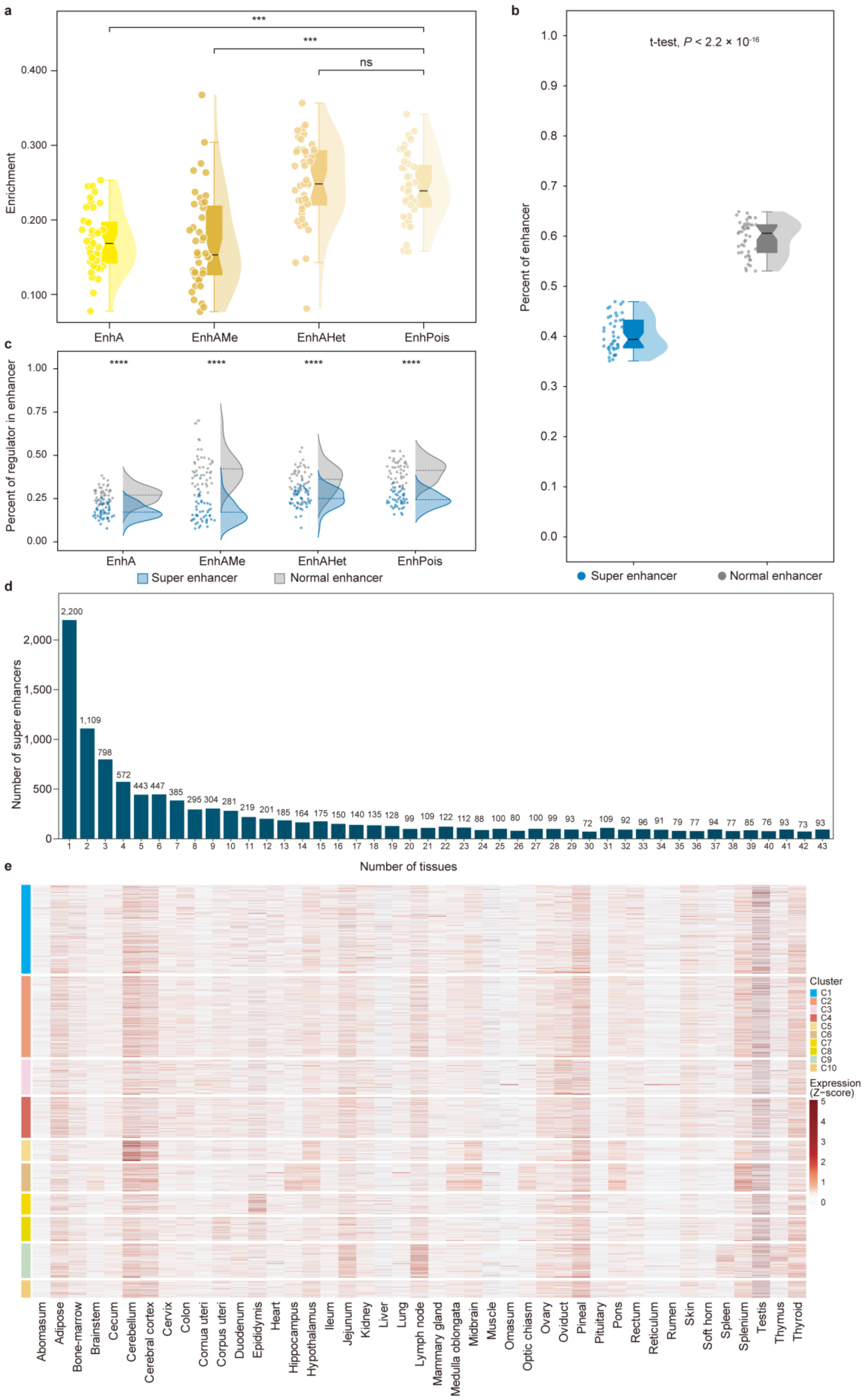
a
***
***
ns
Enrichment
0.400
0.300
0.200
0.100
EnhA
EnhAMe
EnhAHet
EnhPois
b
t-test, P < 2.2 × 10-16
Percent of enhancer
1.0
0.9
0.8
0.7
0.6
0.5
0.4
0.3
0.2
0.1
0.0
Super enhancer
Normal enhancer
c
Percent of regulator in enhancer
****
****
****
****
1.00
0.75
0.50
0.25
0.00
EnhA
EnhAMe
EnhAHet
EnhPois
Super enhancer
Normal enhancer
d
Number of super enhancers
2,000
1,500
1,000
500
0
2,200
1,109
798
572
443 447
385
295 304
281
219
201
185
164
175
150
140
135
128
99
109
122
112
88
100
80
100
99
93
72
109
92
96
91
79
77
94
77
85
76
93
73
93
1 2 3 4 5 6 7 8 9 10 11 12 13 14 15 16 17 18 19 20 21 22 23 24 25 26 27 28 29 30 31 32 33 34 35 36 37 38 39 40 41 42 43
Number of tissues
e
Cluster
C1
C2
C3
C4
C5
C6
C7
C8
C9
C10
Expression (Z-score)
5
4
3
2
1
0
Abomasum
Adipose
Bone-marrow
Brainstem
Cecum
Cerebellum
Cerebral cortex
Cervix
Colon
Cornua uteri
Corpus uteri
Duodenum
Epididymis
Heart
Hippocampus
Hypothalamus
Ileum
Jejunum
Kidney
Liver
Lung
Lymph node
Mammary gland
Medulla oblongata
Midbrain
Muscle
Omasum
Optic chiasm
Ovary
Oviduct
Pineal
Pituitary
Pons
Rectum
Reticulum
Rumen
Skin
Soft horn
Spleen
Splenium
Testis
Thymus
Thyroid

**Supplementary Fig. 9 Summary of super-enhancers in sheep.** a, Enrichment of enhancer-associated chromatin states within super-enhancers (SEs) across tissues, including EnhA, EnhAMe, EnhAHet and EnhPois. b, Comparison of the percentage of enhancer elements between SEs and normal enhancers. c, Distribution of the four enhancer-associated chromatin states within SEs and normal enhancers. d, Number of SEs shared across different numbers of tissues. e, Heatmap showing expression Z-scores of SE-associated target genes across tissues, grouped by SE cluster.

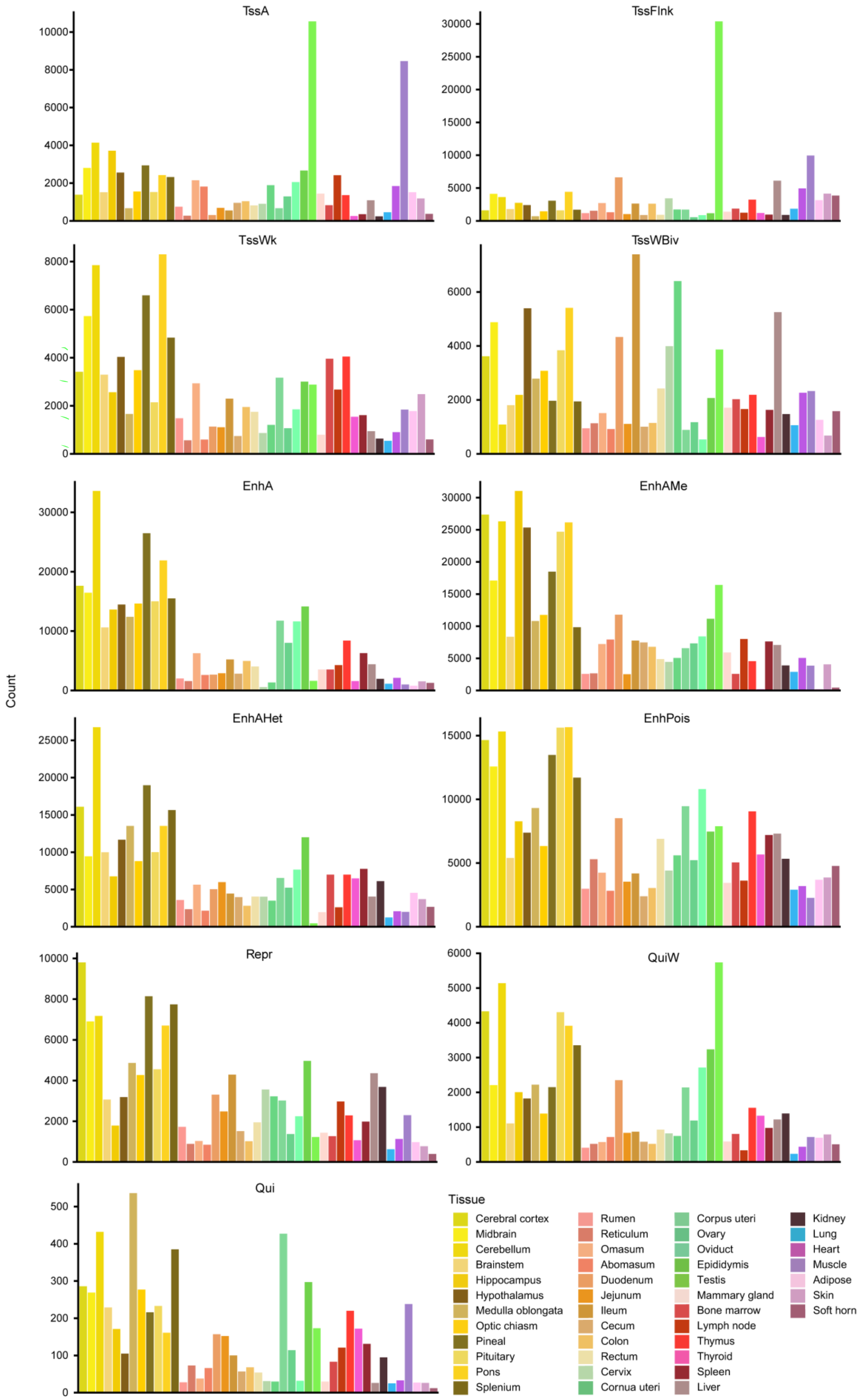
TssA
TssFlnk
TssWk
TssWBiv
EnhA
EnhAMe
EnhAHet
EnhPois
Repr
QuiW
Qui
Count
Tissue
Cerebral cortex
Midbrain
Cerebellum
Brainstem
Hippocampus
Hypothalamus
Medulla oblongata
Optic chiasm
Pineal
Pituitary
Pons
Splenium
Rumen
Reticulum
Omasum
Abomasum
Duodenum
Jejunum
Ileum
Cecum
Colon
Rectum
Cervix
Cornua uteri
Corpus uteri
Ovary
Oviduct
Epididymis
Testis
Mammary gland
Bone marrow
Lymph node
Thymus
Thyroid
Spleen
Liver
Kidney
Lung
Heart
Muscle
Adipose
Skin
Soft horn

**Supplementary Fig. 10 Distribution of tissue-specific regulatory elements across sheep tissues.** Bar plots showing the number of tissue-specific regulatory elements for each chromatin state across 43 sheep tissues.

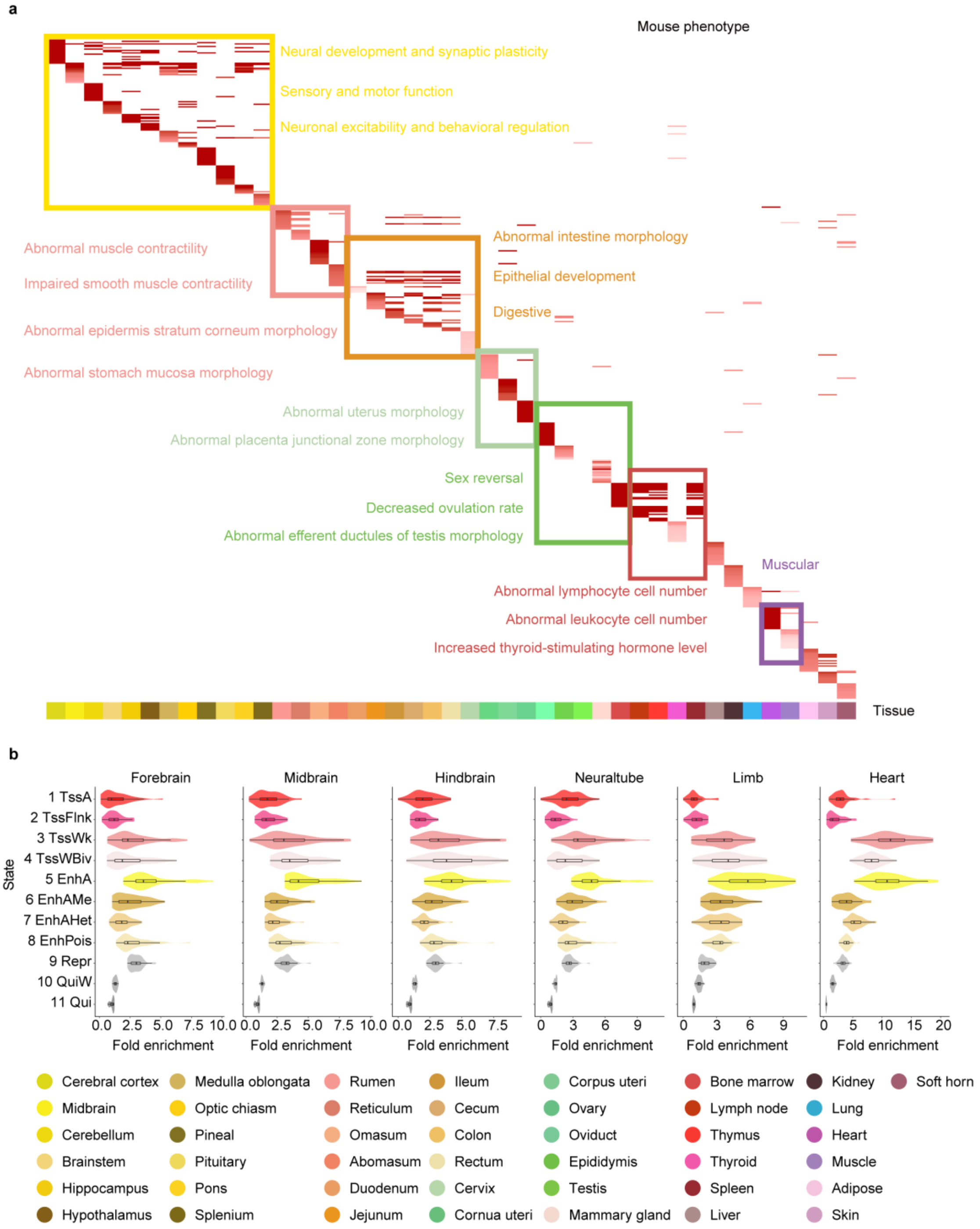


**Supplementary Fig. 11 Functional and developmental characterization of sheep regulatory elements.** a, Mouse phenotype enrichment based on mouse orthologues of genes associated with tissue-specific EnhA. Representative phenotype categories are indicated. b, Fold enrichment of VISTA enhancers across the 11 chromatin states in forebrain, midbrain, hindbrain, neural tube, limb and heart annotations.

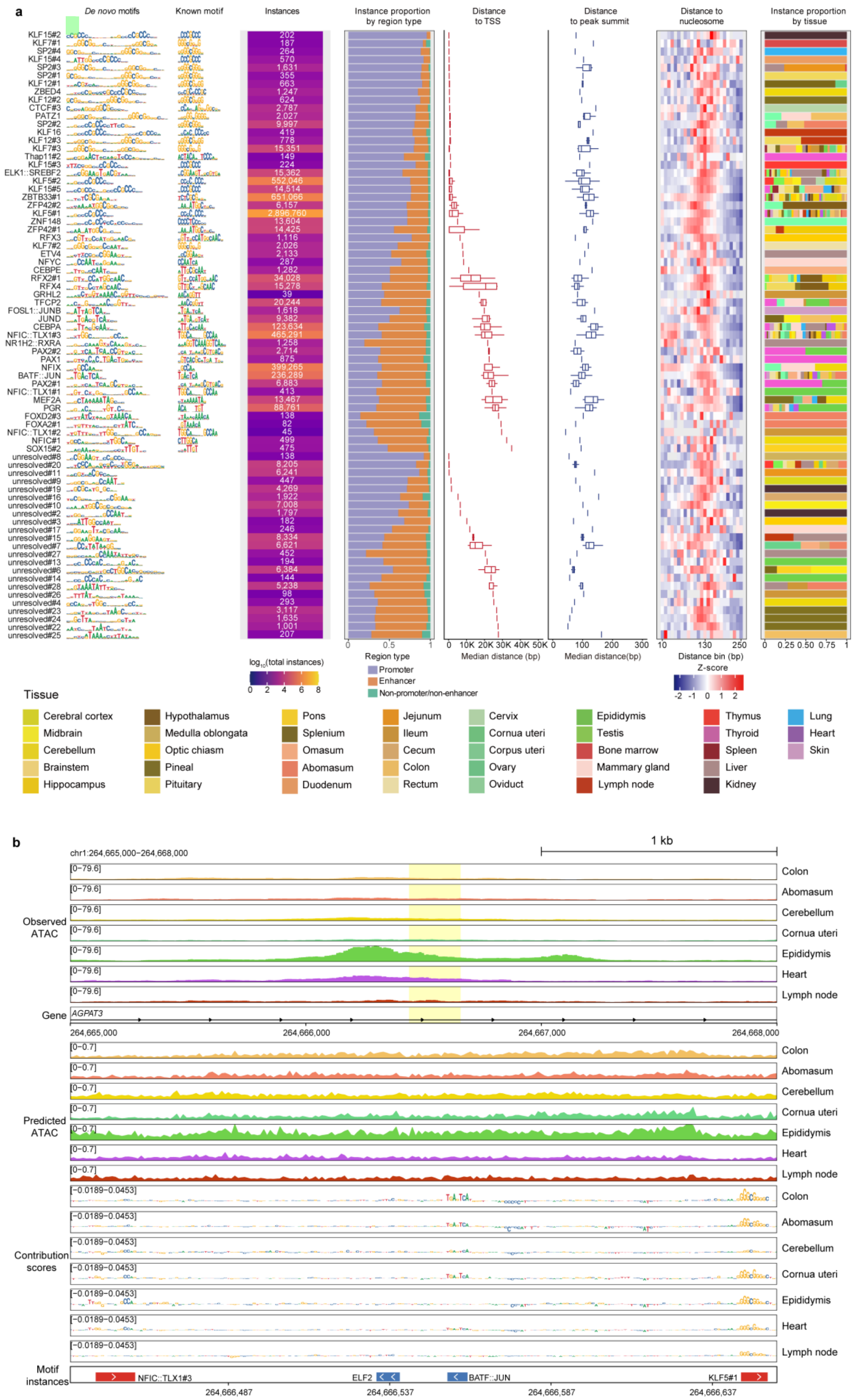
a
De novo motifs
Known motif
Instances
Instance proportion by region type
Distance to TSS
Distance to peak summit
Distance to nucleosome
Instance proportion by tissue
log10(total instances)
Region type
Promoter
Enhancer
Non-promoter/non-enhancer
Median distance (bp)
Median distance(bp)
Distance bin (bp)
Z-score
Tissue
Cerebral cortex
Midbrain
Cerebellum
Brainstem
Hippocampus
Hypothalamus
Medulla oblongata
Optic chiasm
Pineal
Pituitary
Pons
Splenium
Omasum
Abomasum
Duodenum
Jejunum
Ileum
Cecum
Colon
Rectum
Cervix
Cornua uteri
Corpus uteri
Ovary
Oviduct
Epididymis
Testis
Bone marrow
Mammary gland
Lymph node
Thymus
Thyroid
Spleen
Liver
Kidney
Lung
Heart
Skin
b
chr1:264,665,000–264,668,000
1 kb
Observed ATAC
Gene
AGPAT3
Predicted ATAC
Contribution scores
Motif instances
NFIC::TLX1#3
ELF2
BATF::JUN
KLF5#1
264,665,000
264,666,000
264,667,000
264,668,000
264,666,487
264,666,537
264,666,587
264,666,637

**Supplementary Fig. 12 Motif-instance features of 75 additional *de novo* motifs and a genome-browser example at the *AGPAT3* locus.** a, Summary of motif-instance features for 75 additional *de novo* motifs not shown in the main figure. Each row shows the *de novo* motif, closest known motif, total motif-instance number, genomic distribution, distances to the nearest TSS, ATAC-seq peak summit and nucleosome dyad, and tissue distribution. b, Genome-browser view of the *AGPAT3* locus on chr1:264,665,000–264,668,000 across representative tissues, showing observed ATAC-seq signal, ChromBPNet-predicted ATAC-seq signal, base-level contribution scores and predicted motif instances.

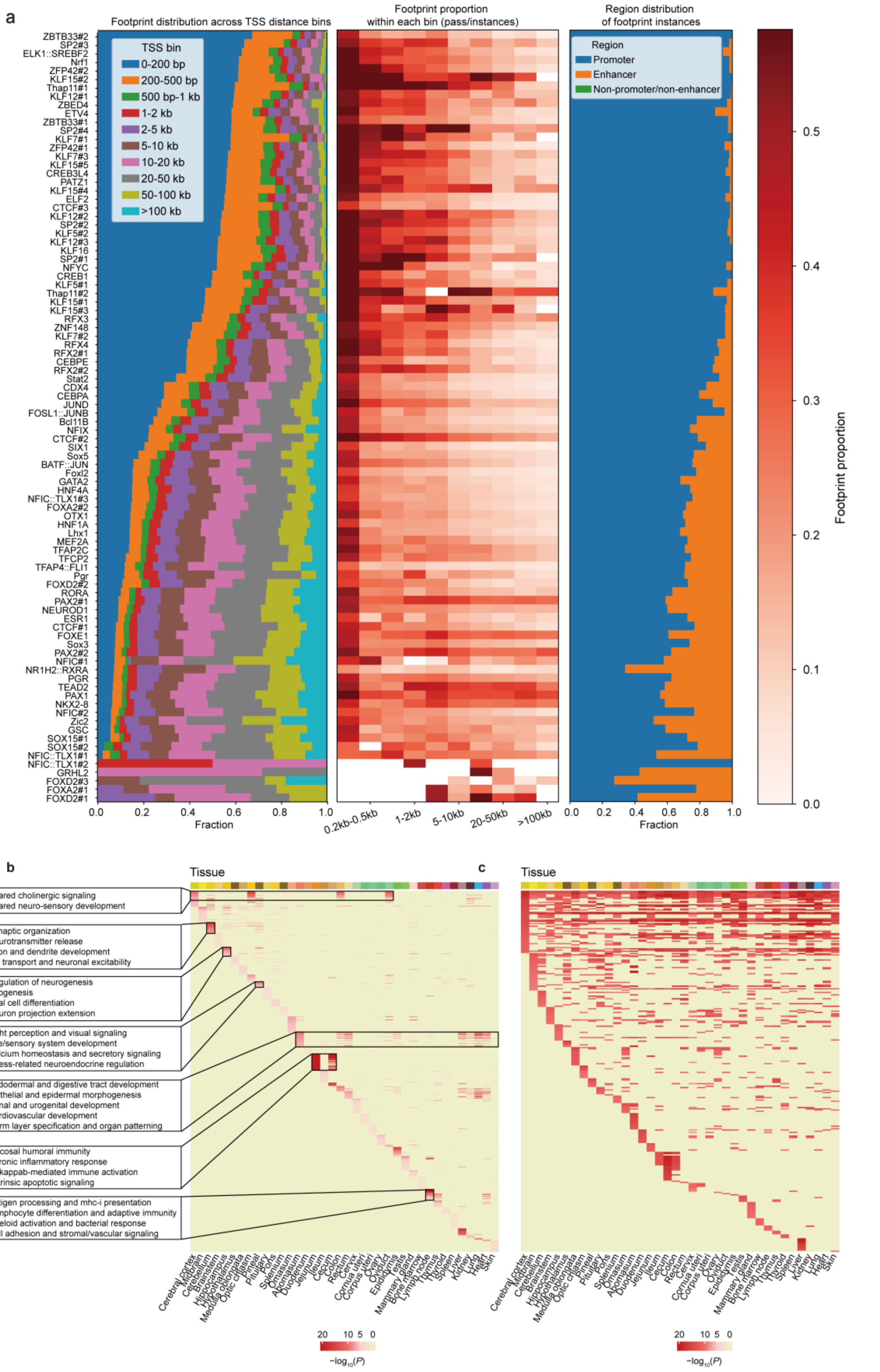


**Supplementary Fig. 13 Footprint-instance distributions and tissue-pair GO enrichment heatmaps of footprint-linked genes.** a, Summary of footprint-instance features for individual predictive motifs, including footprint-instance distribution across TSS-distance bins, footprint proportion within each bin and genomic distribution across promoter, enhancer and non-promoter/non-enhancer regions. b, Tissue-pair heatmap based on GO enrichments of genes linked to enhancer-associated footprint instances. Colour intensity indicates $-\log_{10}(P)$. c, Tissue-pair heatmap based on GO enrichments of nearest genes linked to promoter-associated footprint instances. Colour intensity indicates $-\log_{10}(P)$.

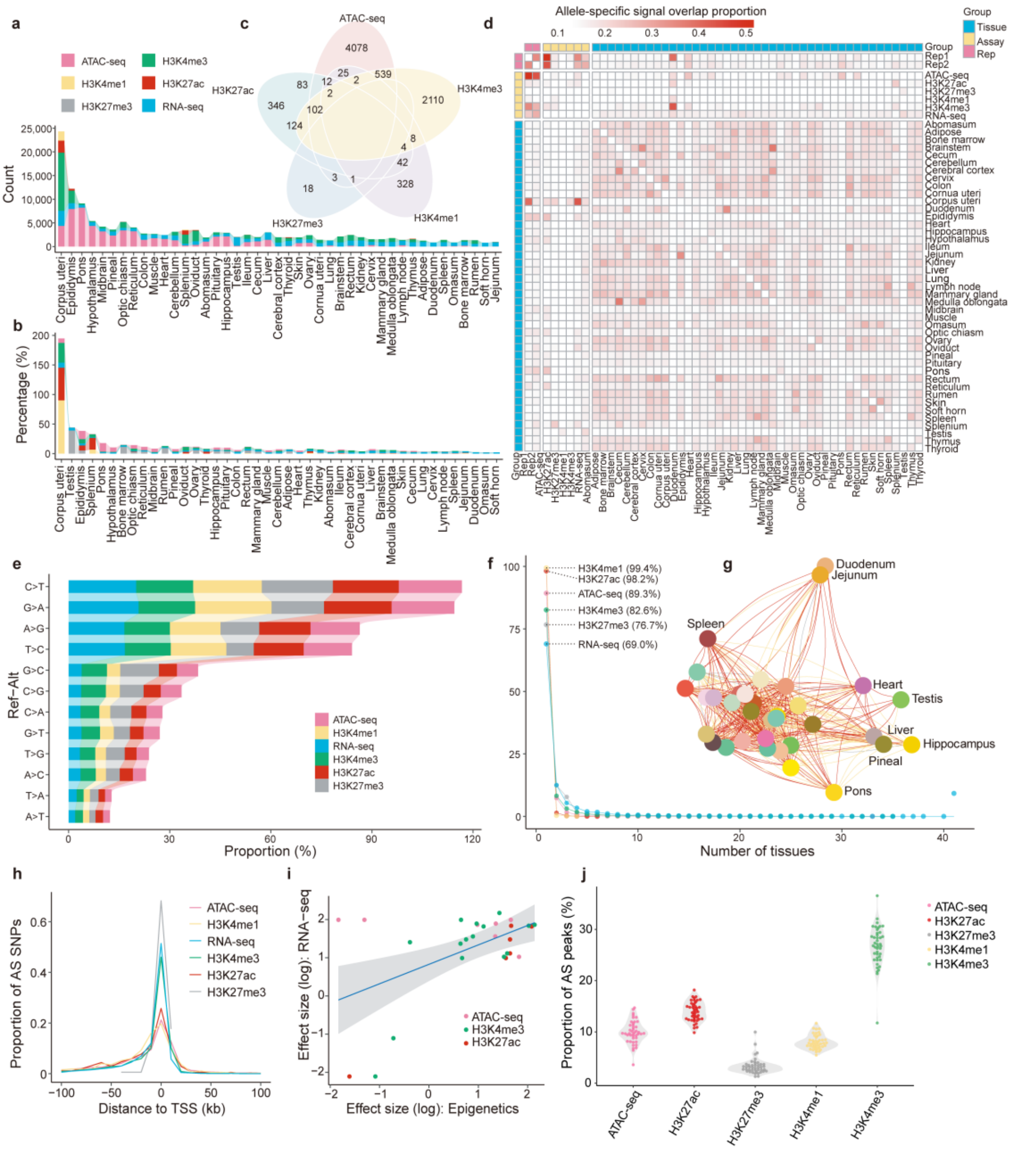


**Supplementary Fig. 14 Summary characteristics and cross-assay patterns of allele-specific signals.** a, Counts of allele-specific signals across tissues and assays. b, Percentage of allele-specific signals across tissues and assays. c, Overlap of allele-specific signals among ATAC-seq and histone-mark assays. d, Heatmap showing pairwise allele-specific signal overlap proportions across tissue, replicate and assay combinations. Row and column annotations indicate tissue, replicate and assay groups. e, References-to-alternative allele substitution spectrum across assays. f, Tissue-sharing patterns of allele-specific signals detected in each assay. g, Tissue-level network showing shared allele-specific regulatory signals among tissues.

h, Distribution of allele-specific SNPs relative to transcription start sites across assays. i, Concordance of allelic effect sizes between epigenomic and RNA-seq allele-specific signals. j, Proportion of allele-specific peaks across chromatin states.

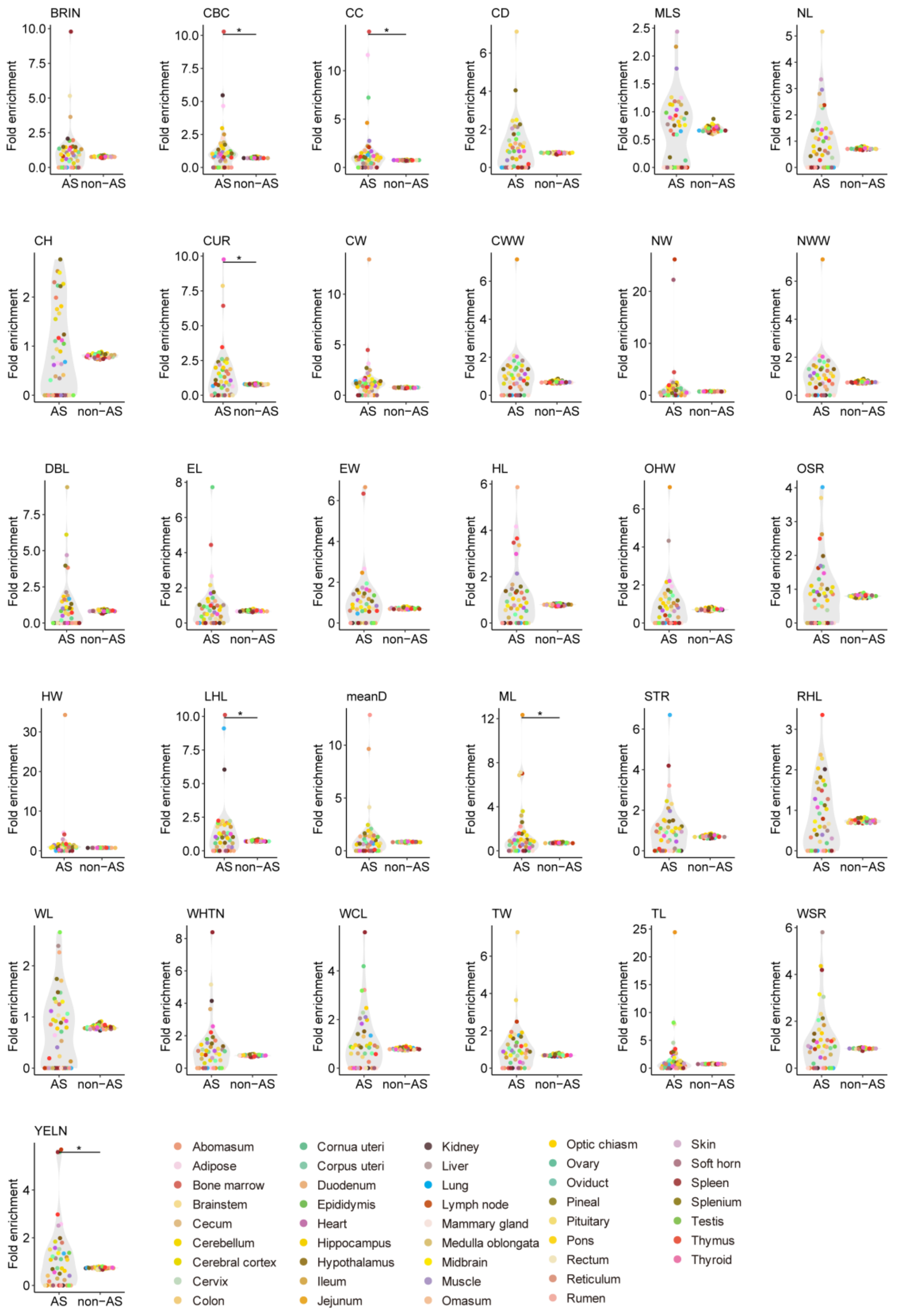


**Supplementary Fig. 15 Enrichment of GWAS signals in AS-containing and non-AS active enhancers.** Fold enrichment of GWAS loci for sheep complex traits in active enhancer elements containing allele-specific sites and active enhancer elements without allele-specific sites. Each panel represents one trait, and points represent tissue-level enrichment estimates. The x axis separates AS-containing EnhA and non-AS EnhA, and the y axis shows fold enrichment.

Asterisks indicate significant differences between groups.

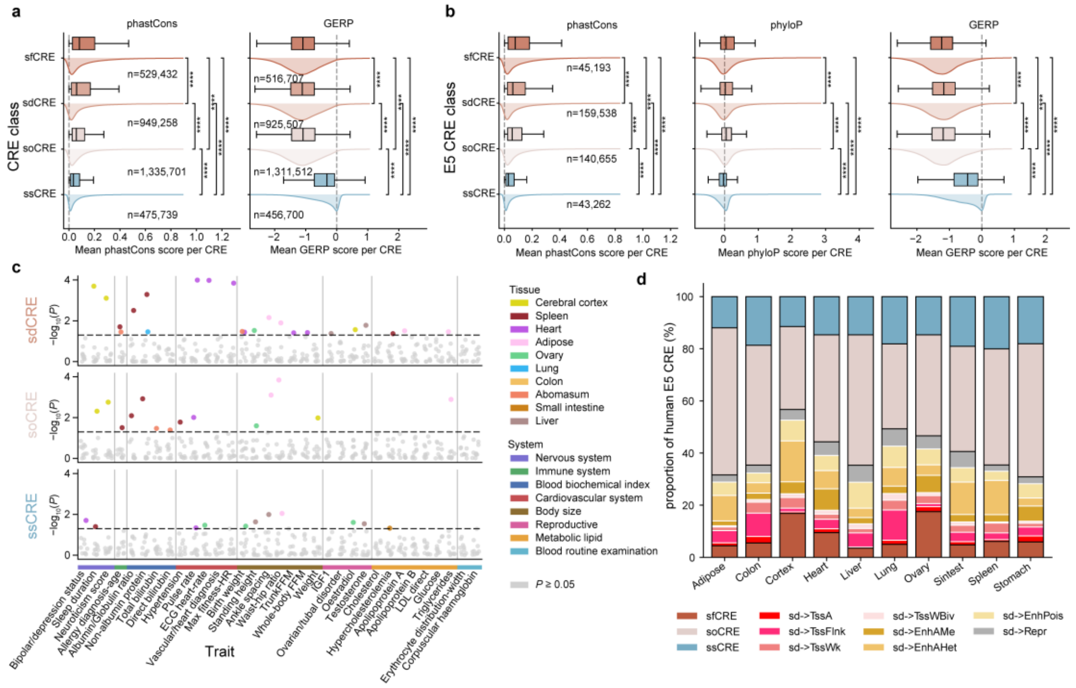


**Supplementary Fig. 16 Conservation scores and class-specific enrichment patterns of human–sheep cis-regulatory elements.** a, Distributions of mean phastCons and GERP scores across sfCRE, sdCRE, soCRE and ssCRE classes. Asterisks denote significance levels. b, Distributions of mean phastCons, phyloP and GERP scores for human E5 cis-regulatory elements stratified by sfCRE, sdCRE, soCRE and ssCRE classes. Asterisks denote significance levels. c, GWAS enrichment signals for sdCRE, soCRE and ssCRE classes across human traits and tissues. Coloured points indicate nominally significant tissue–trait signals, grey points indicate non-significant tests and the dashed line indicates nominal $P = 0.05$. d, Proportional composition of human E5 cis-regulatory elements across matched tissues, including sfCRE, soCRE and ssCRE classes, together with sdCREs assigned to sheep-side chromatin states.

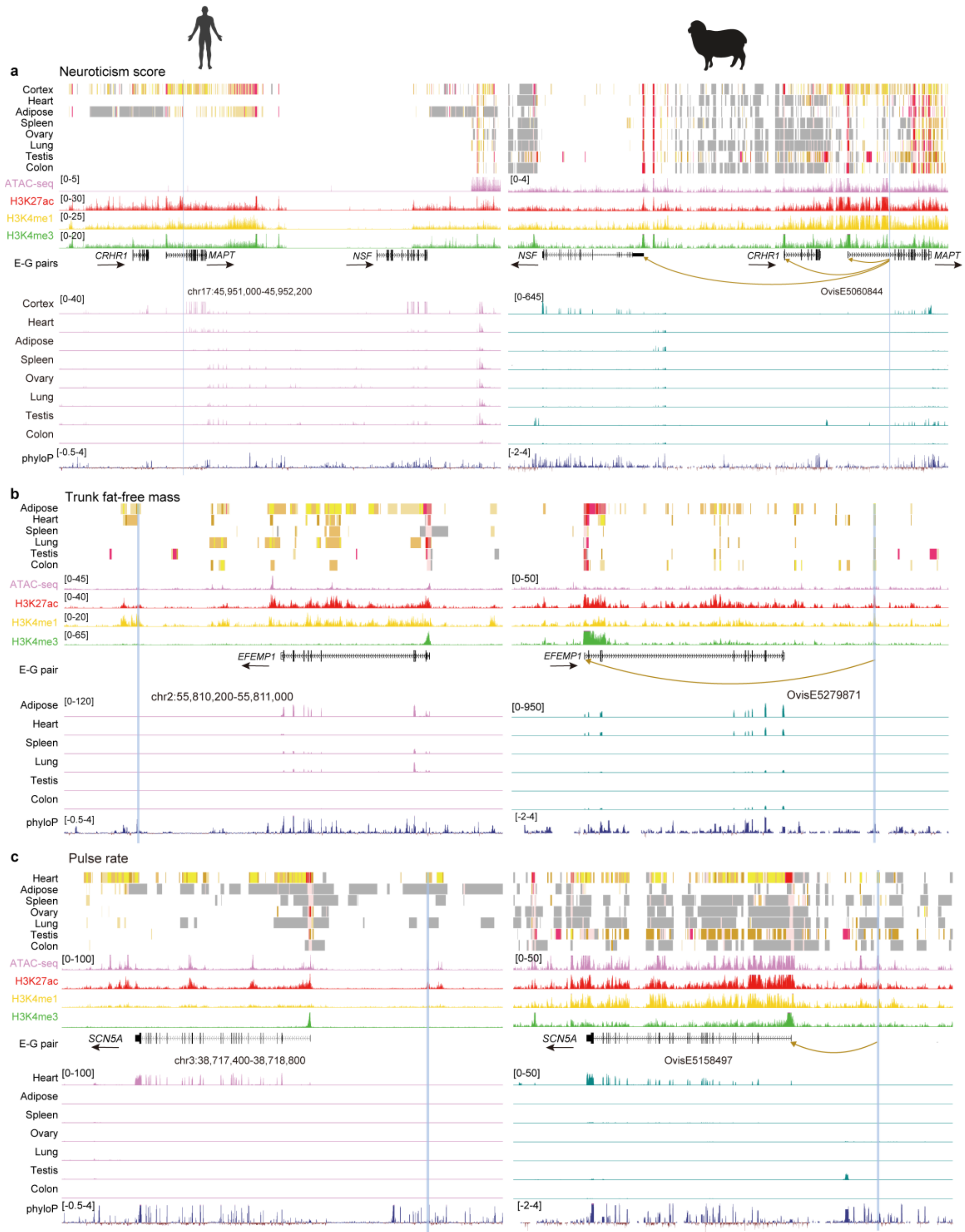


**Supplementary Fig. 17 Cross-species regulatory evidence for representative conserved GWAS-associated loci.** a, Locus-level regulatory annotation of the neuroticism-score-associated region near *CRHR1*, *MAPT* and *NSF*, showing matched human and sheep regulatory tracks and the corresponding sheep regulatory element OvisE5060844. b, Locus-level regulatory annotation of the trunk-fat-free-mass-associated region near *EFEMP1*, showing matched human and sheep regulatory tracks and the corresponding sheep regulatory element OvisE5279871. c, Locus-level regulatory annotation of the pulse-rate-associated region near *SCN5A*, showing matched human and sheep regulatory tracks and the corresponding sheep regulatory element OvisE5158497. Tracks include chromatin-state annotations, ATAC-seq, H3K27ac, H3K4me1, H3K4me3 and phyloP signals across selected tissues. GWAS, genome-wide association study.

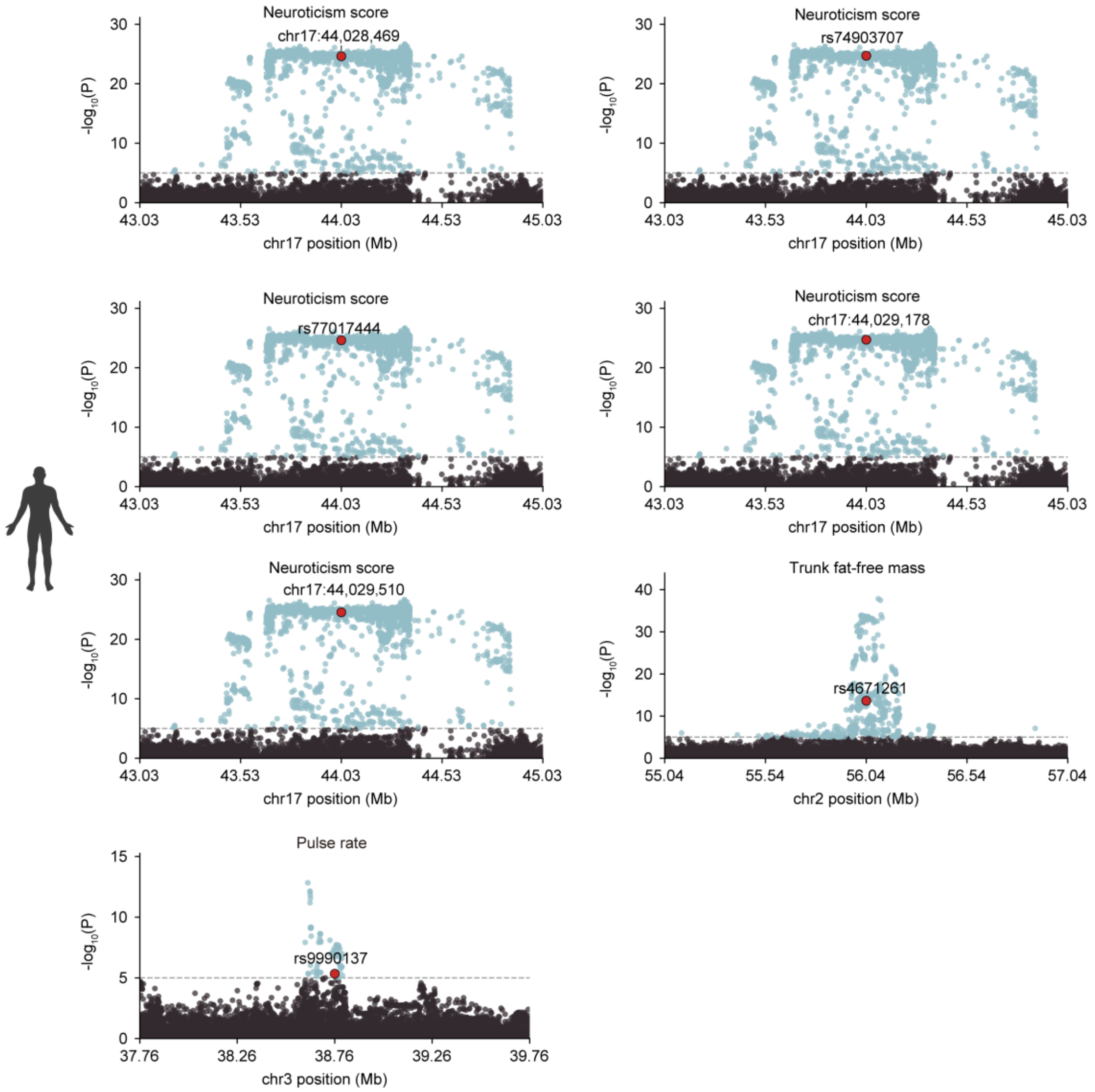


**Supplementary Fig. 18 Regional GWAS association signals overlapping conserved regulatory elements.** a, Regional association plots for representative human GWAS loci overlapping conserved regulatory elements, including neuroticism score loci on chr17, a trunk fat-free mass locus on chr2 and a pulse-rate locus on chr3. The x axis shows genomic position, and the y axis shows −log10($P$). Labelled variants indicate the representative GWAS variants highlighted for each locus.
